# The rate of evolution of postmating-prezygotic reproductive isolation in *Drosophila*

**DOI:** 10.1101/142059

**Authors:** David A. Turissini, Joseph A. McGirr, Sonali S. Patel, Jean R. David, Daniel R. Matute

## Abstract

Reproductive isolation (RI) is an intrinsic aspect of species, as described in the Biological Species Concept. For that reason, the identification of the precise traits and mechanisms of RI, and the rates at which they evolve, is crucial to understanding how species originate and persist. Nonetheless, precise measurements of the magnitude of reproductive isolation are rare. Previous work has measured the rates of evolution of prezygotic and postzygotic barriers to gene flow, yet no systematic analysis has carried out the study of the rates of evolution of postmating-prezygotic (PMPZ) barriers. We systematically measured the magnitude of two barriers to gene flow that act after mating occurs but before zygotic fertilization and also measured a premating (female mating rate in nonchoice experiments) and two postzygotic barriers (hybrid inviability and hybrid sterility) for all pairwise crosses of species within the *Drosophila melanogaster* subgroup. Our results indicate that PMPZ isolation evolves faster than hybrid inviability but slower than premating isolation. We also describe seven new interspecific hybrids in the group. Our findings open up a large repertoire of tools that will enable researchers to manipulate hybrids and explore the genetic basis of interspecific differentiation, reproductive isolation, and speciation.

## INTRODUCTION

Barriers to gene flow, or reproductive isolating mechanisms (RIMs), evolve as a byproduct of divergence between populations that accrue genetic differences over time. The process of speciation is thus the accumulation of RIMs. The strength of reproductive isolation dictates whether nascent species persist or whether they merge into a single lineage once they come into contact with each other. In cases where RI is absolute and no intermixing through hybridization is possible, speciation is complete. In other cases, RIMs are weak and can be overcome by gene flow, thus merging nascent species into a single lineage. There is an intermediate scenario, which is likely to be common, in which hybridization—and admixture —occurs but species persist. Therefore, the nature and magnitude of RIMs that evolve between groups and the rate at which they evolve are key factors influencing the origin of new species. The systematic identification of these barriers in a phylogenetic context (to infer their rates of evolution) is a prerequisite for understanding which barriers are important drivers of speciation and which result from post-speciation divergence.

Depending on when they occur in the reproductive cycle, RIMs may be classified as premating, postmating-prezygotic, or postzygotic (Orr and Presgraves 2000; Presgraves 2010). Premating RIMs encompass all the biological traits that preclude populations from encountering or mating with each other. Niche specificity, habitat preferences, reproductive timing, and mate choice are all examples of premating barriers. A second type of barrier that acts after mating but before a zygote is formed (i.e. postmating prezygotic [PMPZ] barriers) involves discordant interactions between gametes or between the female reproductive tract and components of the male seminal fluid. Gametic interactions include the physical and chemical cues that allow for mutual gametic recognition and eventual fusion into a zygote. Gametic incompatibilities may arise if these cues are incompatible between gametes from different species, thereby restricting gene flow. In organisms with internal fertilization, less is known about the evolution and prevalence of PMPZ RIMs compared to premating or postzygotic mechanisms (i.e. fitness reductions seen in interspecific hybrid individuals and not in the pure species (Dobzhansky 1937; Coyne and Orr 2004)) (but see Birkhead and Pizzari 2002; Sweigart 2010; Larson et al. 2012).

Several meta-analyses have inferred the rate at which RIMs evolve over time and most have found a positive relationship between their strength and genetic distance (reviewed in Edmands 2002). These results show that premating isolation usually evolves before postzygotic isolation in *Drosophila*, amphibians, and certain groups of plants and fish (reviewed in Coyne and Orr 2004). This body of work has led to the widespread notion that prezygotic isolation is necessary to initiate speciation (e.g., Abbot et al. 2009, Seehausen et al. 2014 among many others). However, premating and postzygotic isolation have similar rates of evolution in some plant genera (reviewed in Widmer et al. 2009), and in copepods, postzygotic isolation evolves before prezygotic isolation (Ganz and Burton 1995, Palmer and Edmands 2000, Edmands et al. 2009). Clearly, more comparative work, in terms of traits and taxa, is needed before a conclusion on what RIMs (if any) are responsible for setting the process of speciation in motion.

In contrast to studies that measure the strength and rates of evolution of premating and postzygotic isolation, the evolutionary rates of PMPZ isolation have rarely been investigated, with the notable exception of plant taxa. In orchids and *Fragaria*, there is no apparent correlation between the magnitude of prezygotic isolation (either premating isolation or post-pollination prezygotic, the equivalent of PMPZ) and genetic distance (Scopece et al. 2007; Scopece et al. 2008; Nosrati et al. 2011). In *Glycine* (Fabaceae) and *Silene* (Caryophyllaceae), post-pollination prezygotic and postzygotic isolation both increase monotonically with divergence time and at similar rates (Moyle et al. 2004). In Chilean Bellflowers (*Nolana*, Solanaceae), postzygotic isolation evolves faster than post-pollination prezygotic isolation (Jewell et al. 2012). The results from these five taxa suggest that post-pollination prezygotic isolation is important but heterogeneous across groups.

In the case of animals, even fewer studies have explored the effect of genetic distance on the magnitude of PMPZ isolation. This is surprising because this type of barrier seems to be common (e.g. Fricke and Arnqvist 2004, Mendelson et al. 2007, Dopman et al. 2010). Gametic incompatibilities are crucial in maintaining species boundaries in sea urchins of the genus *Echinometra*. Qualitative measurements revealed no apparent increase in the magnitude of gametic isolation in two species pairs (Lessios and Cunningham 1990). A second study measured the magnitude of conspecific sperm precedence in two pairs of species of *Drosophila* and suggested that this type of gametic barrier evolves after premating isolation but allowed for no comparison with other barriers (Dixon et al. 2003). Finally, a comparative analysis of in vitro fertilization rates (i.e., percentage of fertilized eggs) in toads revealed no effect of the level of genetic distance between the parental species on gametic interactions (Malone and Fontenot 2008). These disparate conclusions indicate that a more systematic approach is needed to measure the rate of evolution of these traits.

We measured the rate of evolution of reproductive isolation in a common environment for all possible hybridizations between all 9 species of the *Drosophila melanogaster* species subgroup. We provide fine scale measurements of two PMPZ RIMs: non-competitive gametic isolation (i.e., the number of eggs a female lays after a heterospecific mating) and conspecific sperm precedence (i.e., the number of individuals a conspecific male sires after mating with a female that also mated with a heterospecific male). We also improve upon previous summaries of premating and postzygotic isolation in the *melanogaster* subgroup by attempting all possible hybridizations in the group, measuring the magnitude of these barriers in a controlled laboratory environment, and incorporating genome-wide information to quantify genetic distance between species.

Our results show that PMPZ barriers evolve faster than postzygotic RIMs but slightly slower than premating RIMs. Overall, we show that PMPZ RIMs might have important evolutionary consequences in initiating speciation and in the persistence of new species.

## RESULTS

Our goal was to quantify the magnitude of four mechanisms of reproductive isolation—premating isolation, non-competitive gametic isolation, conspecific sperm precedence, and postzygotic isolation—in a controlled laboratory environment for all possible crosses between species of the *Drosophila melanogaster* species subgroup. The indexes we used to measure the magnitude of each RIM are shown in Table 1. We report our results for each barrier first, and then compare their rates of evolution using a phylogenetic comparative approach. Finally, we incorporate results from other species groups of the *Drosophila* genus to conduct a phylogenetic comparison that corrects for phylogenetic non-independence.

**TABLE 1.**
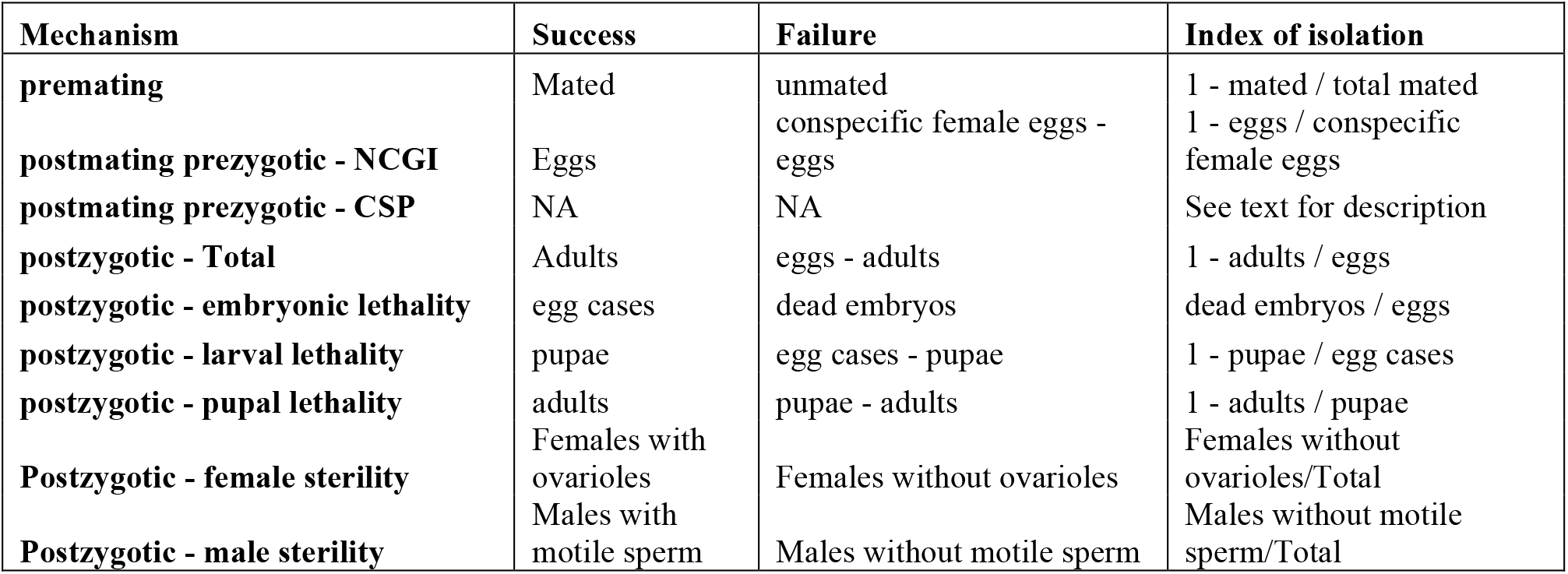
Reproductive isolation barriers studied in this report.

| Mechanism | Success | Failure | Index of isolation |
| --- | --- | --- | --- |
| **premating** | Mated | unmated | 1 - mated / total mated |
| **postmating prezygotic - NCGI** | Eggs | conspecific female eggs - eggs | 1 - eggs / conspecific female eggs |
| **postmating prezygotic - CSP** | NA | NA | See text for description |
| **postzygotic - Total** | Adults | eggs - adults | 1 - adults / eggs |
| **postzygotic - embryonic lethality** | egg cases | dead embryos | dead embryos / eggs |
| **postzygotic - larval lethality** | pupae | egg cases - pupae | 1 - pupae / egg cases |
| **postzygotic - pupal lethality** | adults | pupae - adults | 1 - adults / pupae |
| **Postzygotic - female sterility** | Females with ovarioles | Females without ovarioles | Females without ovarioles/Total |
| **Postzygotic - male sterility** | Males with motile sperm | Males without motile sperm | Males without motile sperm/Total |

### Premating isolation

For each of the 72 possible pairwise combinations of species in the *melanogaster* subgroup, we conducted 24-hour non-choice mating experiments. We estimated the magnitude of behavioral isolation by dissecting females and counting how many were inseminated after 24 hours of being housed with males of another species. As a proxy for mating propensity for the females of each species—which may vary among species and experimental blocks—we measured differences in insemination rates among conspecific crosses. The insemination rate did not differ across conspecific crosses, and was always above 90% (N = 5 one-hundred en masse matings per cross; cross effect, Linear model: F_8,36_ = 0.3658, P = 0.9318; Figure S1). Next, we scored the proportion of females that mated in interspecific crosses. We found that in 33 interspecific pairs, premating isolation is not complete, and insemination occurs (i.e., we found at least one inseminated female). In the other 39 possible interspecific crosses, premating isolation seemed to be complete (i.e., we found no inseminated females), which prevented the study of postmating isolation (see below).

Next, we compared the proportion of inseminated females among heterospecific crosses (Figure S1). We found significant heterogeneity in the proportion of females that accepted heterospecific males (range: 0%−37%; linear model—LM—: F_71,288_ = 35.471, P < 1 × 10^−15^). This heterogeneity persisted when only the 33 different interspecific crosses for which mating occurred were included in the linear model (range: 0.5%−37%; LM: F_27,80_ = 19.624, P < 1 × 10^−15^) This analysis revealed two general patterns. Even though this comparison is not phylogenetically independent, pairwise comparisons revealed that not surprisingly, less diverged species pairs are more prone to hybridize than those that are more diverged (all linear contrasts in Table S1). Second, we found that no female or male genotype were more prone to hybridize with heterospecifics than others (Linear contrasts; all mother levels: t_288_> −0.336, P > 0.737; all father levels: t_288_> −0.237, P > 0.813). The latter result indicates that even though the magnitude of premating isolation differs between pairs, this heterogeneity cannot be attributed to promiscuity of any of the studied species.

### Non-competitive gametic isolation (NCGI)

We next measured PMPZ isolation using singly mated females. In these crosses, the number of eggs laid by females inseminated by interspecific males relative to eggs laid by heterospecifically inseminated females is a proxy for non-competitive gametic isolation, a form of reproductive isolation (Wade et al. 1994,). While this measurement includes unfertilized eggs, it remains a reliable proxy of sperm retention and survival (Price et al. 2001, Matute 2010, Sagga and Civetta 2011). We attempted all eight conspecific crosses and 26 interspecific crosses by conducting 1,000 no-choice mating trials per cross type. (7 crosses only produced heterospecifically mated females en masse—described above—and not when watched individually.) We obtained between 3 and 28 females per heterospecific cross. As expected, pure species crosses vary in the number of eggs an inseminated female lays after mating with males of their own type (N ≥ 3 mated females per cross; LM; F_6,137_ = 38.784; P < 1 × 10^−15^). We normalized the egg counts of females mated with heterospecific males by the average number of eggs laid by a female of that species following conspecific mating. This constitutes a proxy of the maximum number of eggs a female of a given species can produce (i.e., conspecific fertility). Divergence from this average is our measure of NCGI. Levels of NCGI for each cross are shown in Figure S2. We found substantial heterogeneity in the magnitude of NCGI in heterospecific crosses (LM, F_13, 357_ = 44.338; P < 1 × 10^−15^). Pairwise comparisons show that crosses between divergent species tend to produce fewer eggs than crosses between younger species (Table S2). These results provide evidence that postmating interactions between gametes and/or between ejaculate (sperm + seminal fluid) and the female reproductive tract have diverged in distantly related species resulting in fewer eggs.

### Conspecific sperm precedence

In species that show little NCGI, competitive interactions between heterospecific sperm may still constitute an important RIM (citations). We thus measured conspecific sperm precedence (CSP) using doubly-mated females. We obtained progeny in 32 out of 144 possible interspecific crosses (Table S3). We scored the identity of the progeny sired by a female that mated to two males, one conspecific and one heterospecific. We first explored whether there was heterogeneity in the magnitude of CSP in different crosses. We found a similar pattern as the one observed for both premating isolation and NCGI in which the magnitude of CSP differs across interspecific crosses of *Drosophila* (range of sample size: [1,16] doubly mated females; F_1,234_= 29.805, P < 1 × 10^−15^). Levels of CSP for each cross are shown in Figure S3.

Our proposed index of CSP (I_CSP_) should be bounded between 0 and 1, where 0 indicates no sperm precedence and 1 indicates complete conspecific sperm precedence. Nonetheless, we found two major exceptions to this range. *Drosophila santomea* females mated to *D. yakuba* and then *D. santomea* males (in that order) produced a large proportion (~50%) of *yak/san* hybrids and *CC*_*C*1_ is lower than *HC*_*C*1_ (i.e., in pure species double-matings, females sire few progeny from the first mating; in interspecific matings, females sire an unexpectedly large number of hybrid progeny but only when the interspecific male was first in the order of the mating; see Methods for a full description). This lead to a I_CSP_ value of −2.56. In a less extreme, yet similar case, *D. yakuba* females mated to *D. santomea* and then *D. yakuba* males (in that order) produce more *yak/san* hybrids than expected (similar to the case outlined immediately above) which leads to I_CSP_ value of −0.118. There is significant heterogeneity among crosses either including (Linear Model, F_19,234_= 5.361, P=1.396 × 10^−13^) or excluding these two cases (Linear Model, F_19,210_= 4.1899, P=6.758× 10^−12^). The biological implications of a negative index of sperm precedence are challenging to interpret; because of their uniqueness among the crosses, these two crosses were excluded from any further analyses. Linear contrasts are shown in Table S4 and show that CSP is stronger in crosses between species that are long diverged than in closely related species (i.e., those with a a synonymous substitution per synonymous site rate —Ks— >10%).

### Postzygotic Isolation: hybrid inviability

We followed the fate of fertilized eggs from interspecific crosses through each developmental stage and assessed whether they produced larvae, pupae, and ultimately viable adults. Out of those 33 crosses for which premating isolation is not complete, 32 of them produce viable adult hybrids of at least one sex. Among these 33 crosses, we find seven previously undescribed hybridizations, mostly between highly divergent species of the *yakuba* species complex and the *melanogaster/simulans* clade. The list of hybridizations that produced progeny is shown in Table 2. We found that hybrid inviability is rare in the *D. melanogaster* subgroup, even after 15 million years of divergence. Of all crosses, only ♀ *D. santomea* × ♂ *D. sechellia* showed complete hybrid inviability (Linear contrasts in Table S5). In this cross, half of the progeny died as embryos, and the other half died as pupae. Dissection of these pupae revealed that all individuals had testes (N=34) suggesting (but not confirming) that females died at an earlier developmental stage, possibly as embryos. We found no difference among pure species in their viability (mean = 88.8%; range: 83.9%− 92.0%; F_8,36_ = 1.3119; P = 0.2689), but found extensive heterogeneity in the magnitude of hybrid viability (LM, F_32,132_ = 42.057; P < 1 × 10^−10^, Figure S4). We further dissected the source of this heterogeneity by quantifying inviability in the three developmental stages, and the developmental stages have different viabilities (Table S5). Notably, embryonic lethality rates are not correlated with either larval or pupal lethality rates (embryo vs larvae: ρ= −0.1172, P = 0.1338; embryo vs. pupae: ρ= −0.1718, P = 0.0293). Larval and pupal viability are correlated, suggesting that crosses that show larval lethality are also likely to show pupal lethality (ρ= 0.3806, P = 6.354 × 10^−7^).

**TABLE 2.**
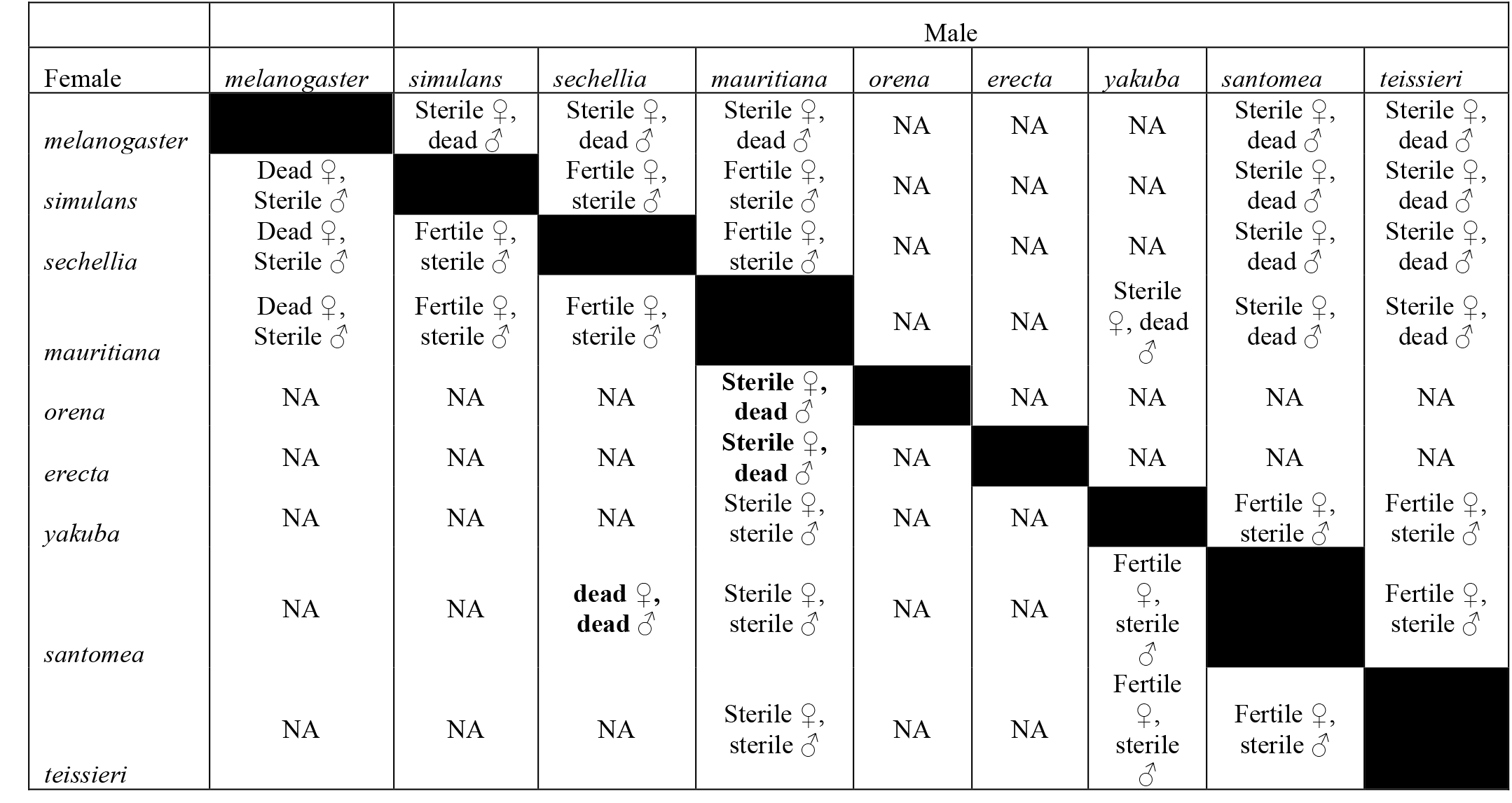
Postzygotic isolation in the *melanogaster* species group. In the majority of crosses for which we observed interspecific matings (i.e., inseminated females), we obtained viable interspecific hybrids. Black cells mark conspecific crosses which produce fertile progeny of both sexes. NA indicates crosses for which we obtained no progeny (i.e., behavioral isolation was complete). Thirty-one hybridizations produced viable progeny out of 72 possible interspecific crosses in the melanogaster species subgroup. Only one cross (♀*D. santomea* × ♂*D. sechellia*). We failed to obtain inseminated females from other 39 crosses.

### Postzygotic Isolation: hybrid sterility

Hybrid female fertility is a largely binomial trait; all females from a single cross were either sterile or all were fertile. Clearly, there is heterogeneity among crosses for hybrid female fertility, as eight crosses produced over 99% fertile F1 females, while sixteen produced only sterile F1 females. The only notable exceptions to this pattern were F1 female hybrids between the divergent species *D. santomea* and *D. teissieri*, and *D. yakuba* and *D. teissieri* which produced ~94% fertile females. These two species pairs are two of the most divergent crosses in *Drosophila* to produce F1 fertile females (Turissini et al. 2015); our findings indicate that not all F1 females from these crosses are fertile, suggesting differential penetrance of the hybrid incompatibilities that eventually lead to ovariole production. Hybrid male fertility was homogeneous as hybrid males were consistently infertile in all interspecific crosses (Table 2).

### Rate of evolution of reproductive isolating mechanisms

We evaluated the rate at which PMPZ isolation (which has rarely been measured in animals) evolves compared to premating and postzygotic isolation. To do this, we tested whether the genetic distance between the parental species influenced the magnitude of reproductive isolation between them. K_s_, the number of per site synonymous substitutions between a pair of species was used as a proxy genetic distance (and therefore divergence time; Table S6), and π_s_, the per synonymous site nucleotide diversity was used as the average phylogenetic distance between individuals of the same species (Table 3). It is worth noting that Ks is a proxy of divergence and it can be slightly affected by codon bias, population size differences, and mutational saturation, especially between divergent species (Akashi and Eyre-Walker 1998, Comeron and Aguade 1998).

**TABLE 3.**
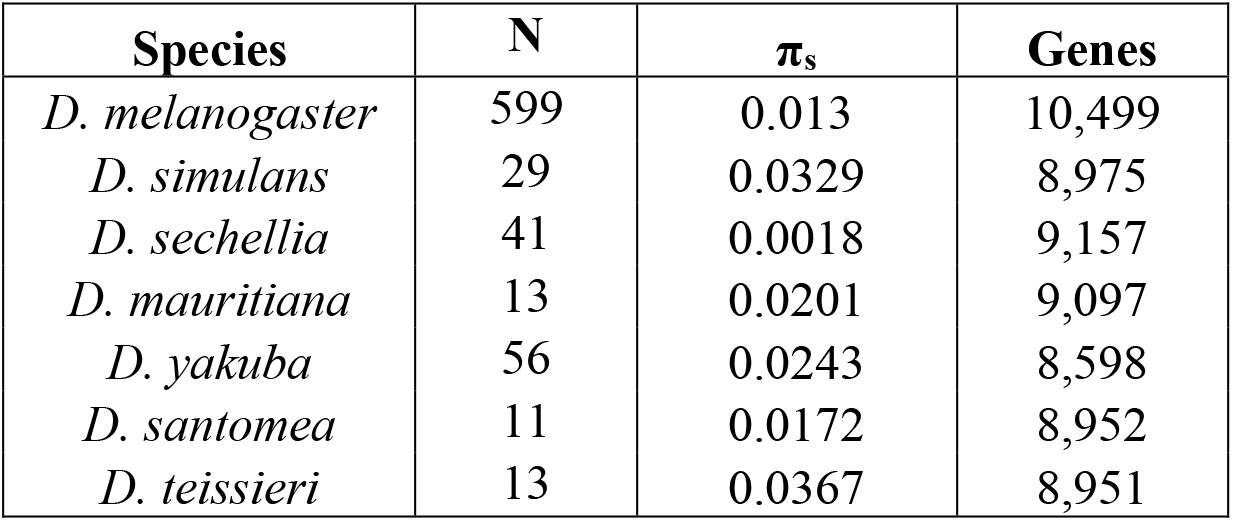
Within species nucleotide diversity. Average heterozygosity values across the whole genome based on synonymous sites (π_s_) values. N represents the number of sequenced lines per species. Since polymorphism data was unavailable for *D. erecta* and *D. orena*, the average of the 7 other species (0.0208) was used in Figure 1. Ks, the genetic distance between species, was calculated with 8,923 genes (Table S6).

| Species | N | $\pi_s$ | Genes |
| --- | --- | --- | --- |
| *D. melanogaster* | 599 | 0.013 | 10,499 |
| *D. simulans* | 29 | 0.0329 | 8,975 |
| *D. sechellia* | 41 | 0.0018 | 9,157 |
| *D. mauritiana* | 13 | 0.0201 | 9,097 |
| *D. yakuba* | 56 | 0.0243 | 8,598 |
| *D. santomea* | 11 | 0.0172 | 8,952 |
| *D. teissieri* | 13 | 0.0367 | 8,951 |

As expected, the magnitude of all types of reproductive isolation scales positively with divergence time. A logistic regression for each of the RIMs showed a strong positive relationship between the magnitude of reproductive isolation and the genetic distance between the parentals (Figure 1). The fit of each of these regressions is shown in Table S7. The increase in premating isolation (Figure 1, red lines) is rapid and (almost) complete at K_s_ >=10% between the hybridizing species. The two mechanisms of PMPZ also follow a similar pattern. The magnitude of both NCGI and CSP is almost complete between species with K_s_ >= 12%. This is in contrast to hybrid inviability, which also scales positively with divergence but evolves more slowly; hybrid inviability is complete in only one of the possible crosses in the *melanogaster* species subgroup (♀*D. santomea* × ♂*D. sechellia*; Figure 1, blue lines).

**FIGURE 1.**
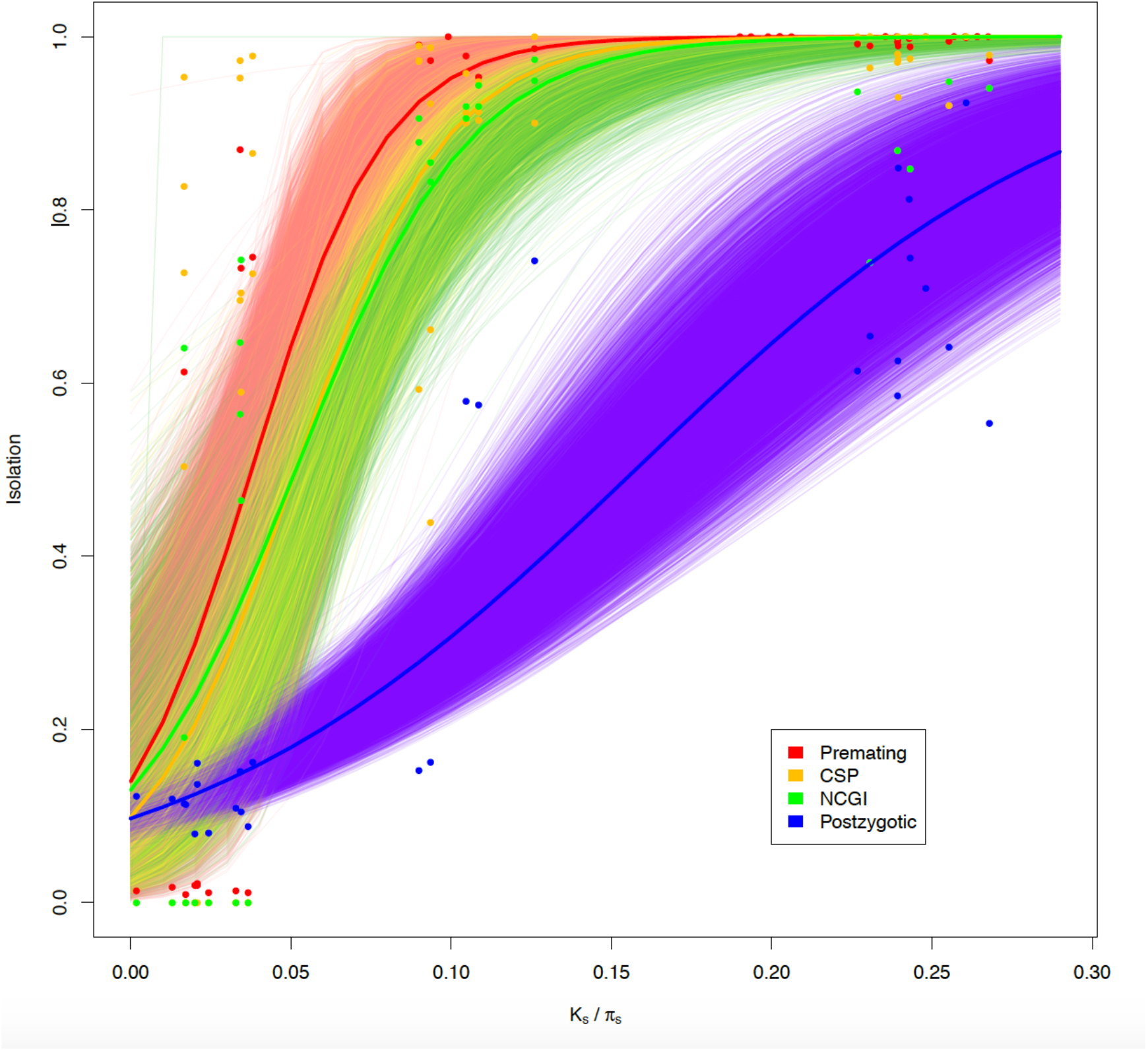
Premating, conspecific sperm precedence, non-competitive gametic isolation, and postzygotic isolation show a strong phylogenetic signal. Proxies of premating isolation (red), conspecific sperm precedence (CSP, yellow), non-competitive gametic isolation (NCGI, green), and postzygotic isolation (blue) were regressed against phylogenetic distance (K_s_ between species and π_s_ within species). The four types of isolation increase with genetic distance, and premating isolation evolves faster than hybrid inviability. The thick red, yellow, green, and blue lines represent fitted logistic regressions for the premating and postzygotic data respectively. The thinner lines of each of the four colors are the regressions for each of 10,000 bootstrap resamplings of the data.

We tested whether any of the four RIMs evolved more quickly than others. (Due to the perfect separation of values along Ks in hybrid sterility, we analyzed this trait separately; see below.) We performed 10,000 bootstrap iterations to assess variation in the effect of divergence time on the strength of each of the four RIMs. Threshold_Ks, the genetic distance at which 95% of RI is achieved (i.e., any of the four indexes of RI equals 0.95), determines how quickly the logistic regression approaches 1, and constitutes a measurement of how fast a RIM evolves. This measurement in bootstrapped datasets was used as a metric for pairwise comparisons. This approach revealed that of all four types of RI, premating isolation evolves quickest followed by the two types of premating-postzygotic isolation (Figure 2; Table S8). We found that of the two PMPZ barriers, CSP evolves faster than non-competitive gametic isolation (Table S8). All prezygotic barriers evolve quicker than postzygotic isolation (Table S8). The relative ranking does not change regardless of the value of the threshold as long as RI > 0.2.

**FIGURE 2.**
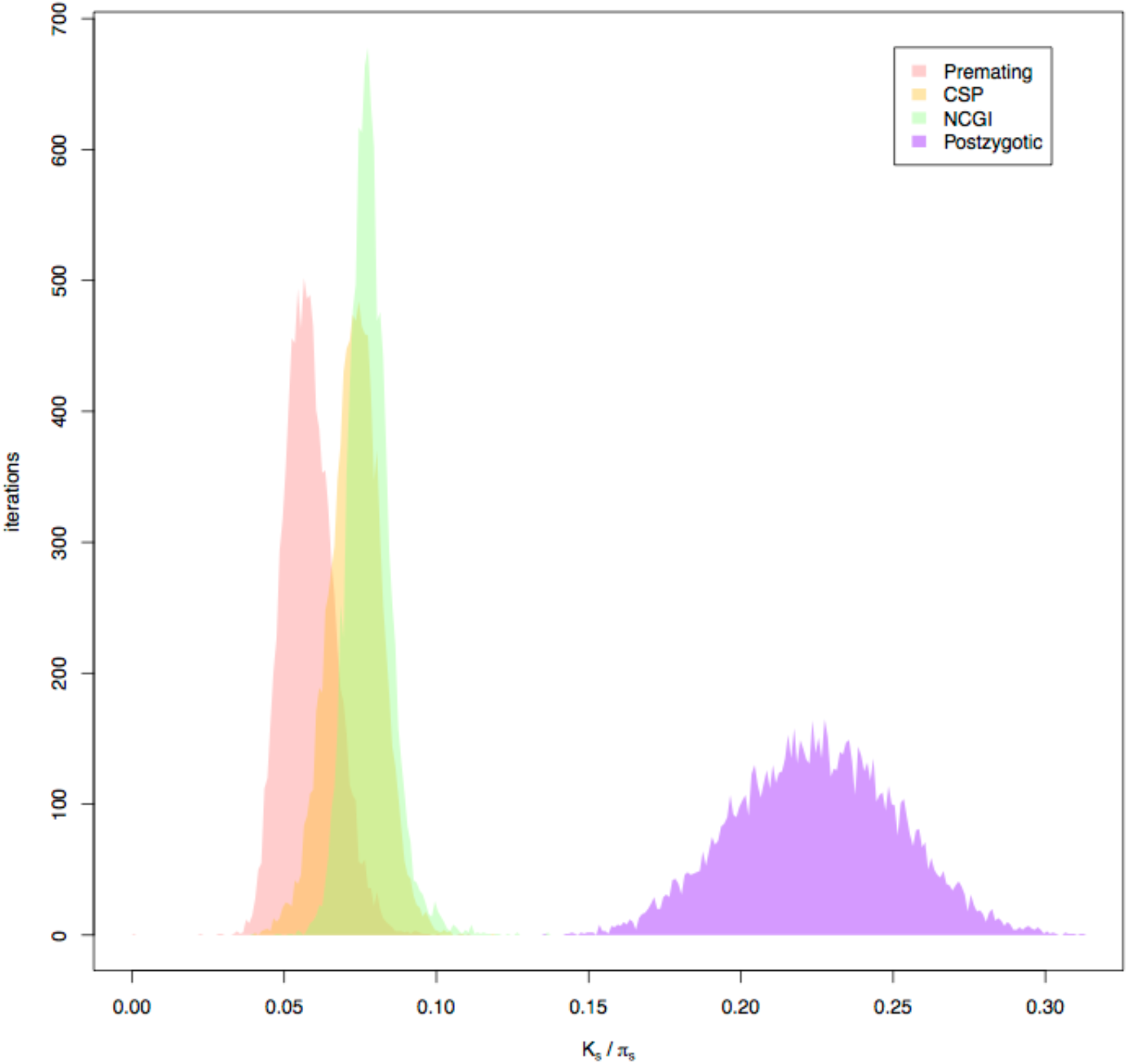
Premating, NCGI, CSP and postzygotic isolation evolve at different rates in the *melanogaster* species subgroup. Threshold_Ks indicates the Ks value for which a given RI barrier achieves a value of 0.95. To assess significance, we compared distributions of bootstrapped values of Threshold_Ks, which determines how quickly isolation approaches 1. All distributions differ from each other in pairwise comparisons. Premating isolation values are red, conspecific sperm precedence (CSP) are yellow, non-competitive gametic isolation (NCGI) are green, and postzygotic values are blue.

We also compared the rates of evolution of hybrid sterility and of hybrid inviability. We analyzed the two sexes separately. First, we compared the rate of evolution of female inviability with that of male inviability. For the former, we assumed that near-complete hybrid female inviability evolved at Ks = 0.25, a very conservative lower bound limit of the genetic distance required for the trait to evolve (Figure 3C). Hybrid male inviability (95% complete) occurs at Ks ~0.15. (This result is identical regardless of how hybrid sterility is measured.) Clearly, hybrid male inviability evolves faster than hybrid female inviability (t test one sample; t_999_= 8385.2, P < 0.0001).

**FIGURE 3.**
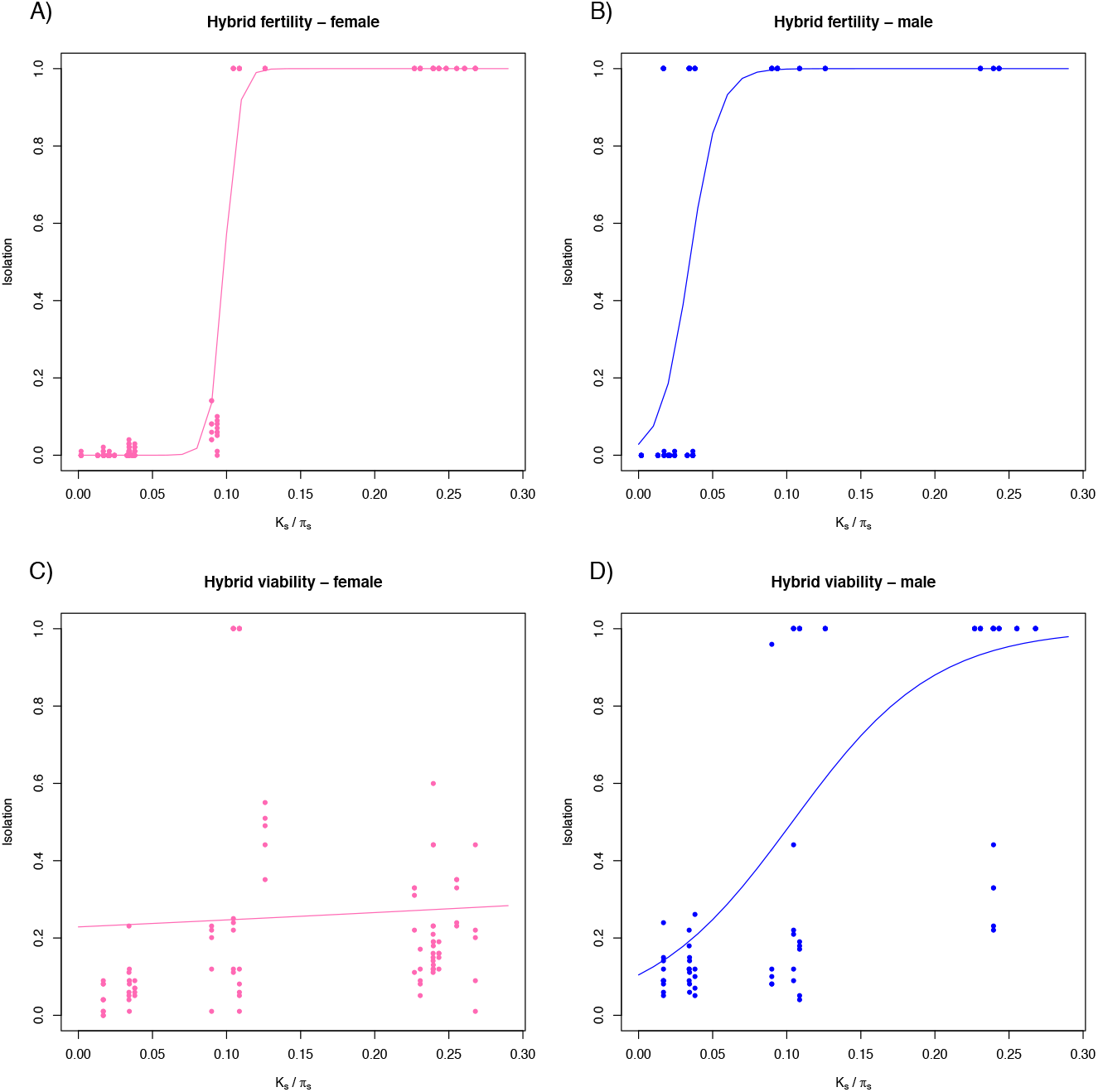
Hybrid sterility evolves faster than hybrid inviability. Values of female and male inviability and female and male sterility were regressed against phylogenetic distance (K_s_ between species and π_s_ within species). The four types of isolation increase with genetic distance. In both sexes, fertility evolves faster than hybrid inviability. Given the perfect separation f values along the x-axis (K_S_/π_S_), these RIMs were not directly compared with other RIMs.

Second, we compared the rates of evolution of hybrid male sterility and of hybrid male inviability. Hybrid male inviability (95% complete) evolves at Ks ~0.15. An equivalent strength of hybrid male sterility takes less divergence to occur (Ks ~0.05). Not surprisingly, we found that hybrid male inviability evolves slower than hybrid male sterility (Wilcoxon sign test: W = 0, P < 1.0 ×10^−15^). A similar comparison in females revealed that female sterility evolves faster than female inviability (even when assuming the lower boundary of possible values for genetic divergence to achieve 95% of the maximum hybrid female inviability; (t test one sample; t_999_= −4706.5, P < 0.0001). It is clear that these three RIMs evolve at different rates (Figure 4). These results confirm the largely accepted, but untested, hypothesis that hybrid sterility evolves faster than hybrid inviability in both sexes.

**FIGURE 4.**
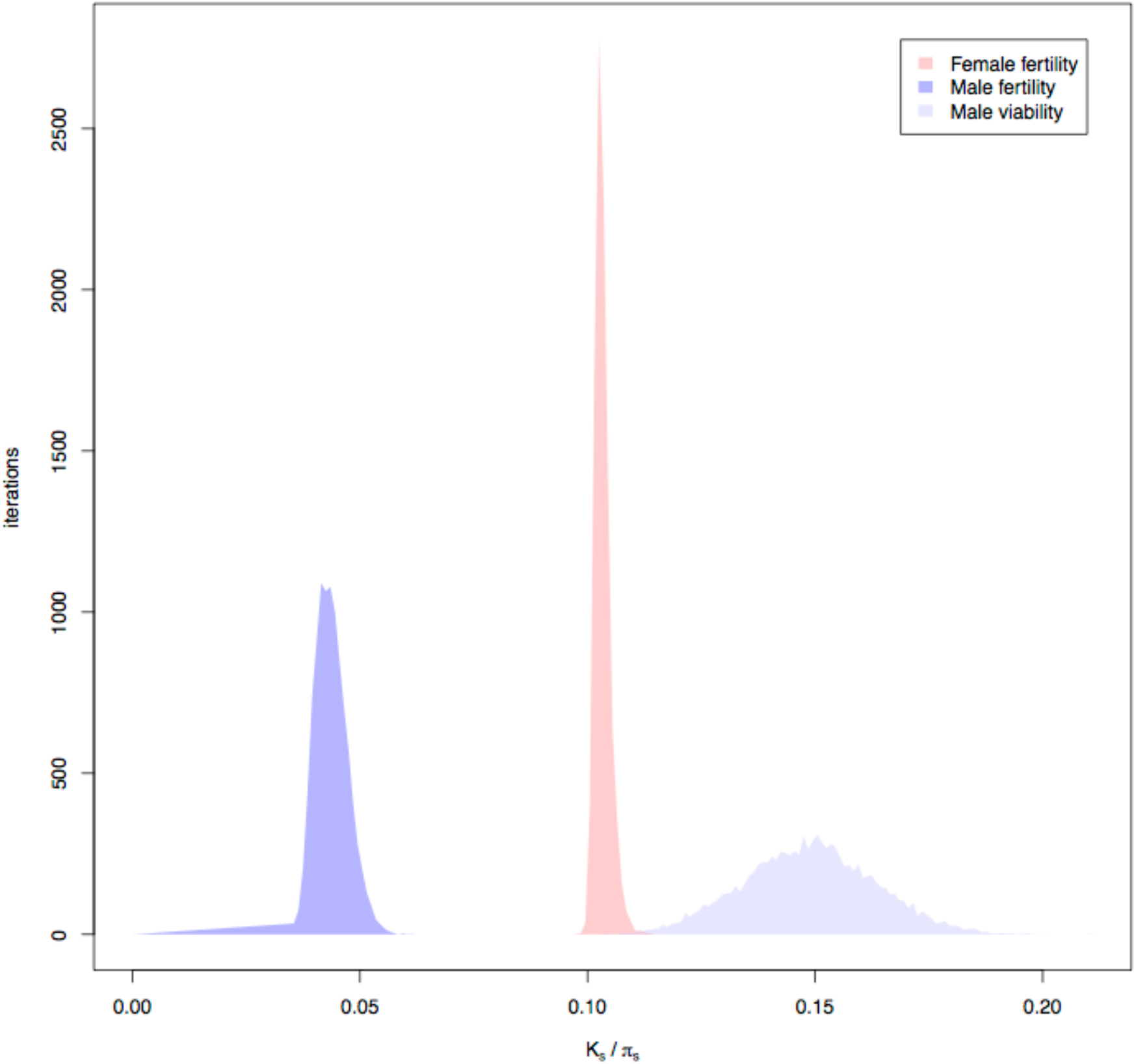
Female sterility, male sterility, and female sterility evolve at different rates in the *melanogaster* species subgroup. Threshold_Ks indicates the Ks value for which a given RI barrier achieves a value of 0.95 (similar to the analyses shown in Figure 2). To assess significance, we compared distributions of bootstrapped values of Threshold_Ks, which determines how quickly isolation approaches 1. Female sterility values are red, male sterility are purple, and male inviability are blue. Female viability did not reach (or approached) an asymptote in our study and for that reason there is no distribution of bootstrapped values for this RIM.

### Rate of evolution of different types of hybrid inviability

Hybrid inviability can manifest within three discrete developmental stages in holometabolan insects: the larvae, the pupae, or the adults. We assessed if hybrid inviability evolved faster at any of these three developmental stages. We quantified developmental stage specific rates of inviability by scoring the number of individuals that die at each of three crucial developmental transitions: embryo-to-L1 larva (embryonic lethality), L1 larvae-to-pupa (larval lethality), and pupa-to-adult (pupal lethality). As expected, the strength of all three types of postzygotic isolation increased with divergence time (Figure 5), and in general, the lowest viabilities were observed for the crosses between the most distantly related species (Table S5). We next compared the rate at which hybrid inviability increased with genetic distance by asking how quickly each type of hybrid inviability reaches 95%. Comparisons of Threshold_Ks rates at each developmental stage showed that embryonic lethality evolves more quickly than larval lethality, and larval lethality evolves faster than pupal lethality (Figure 5D).

**FIGURE 5.**
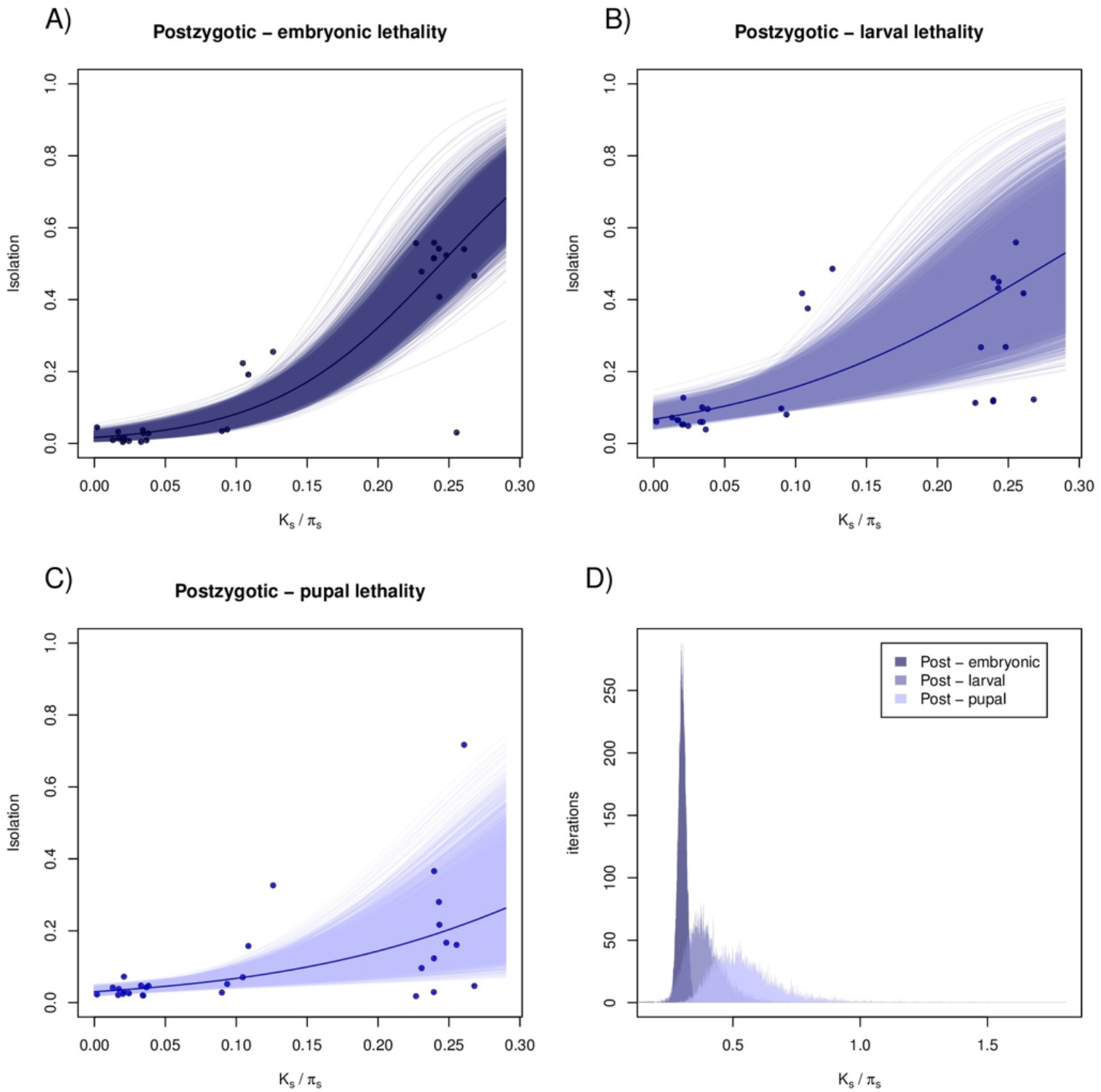
Rates that inviability increases with genetic distance for three developmental stages. Measures of inviability at each developmental stage were regressed against phylogenetic distance (K_s_ between species and π_s_ within species). The three types of postzygotic isolation (i.e., death at a particular developmental stage) increase with genetic distance. **A**. Embryonic lethality. **B**. Larval lethality. **C**. Pupal lethality. The thick lines represent fitted logistic regressions for each developmental stage. The thinner lines are the regressions for each of 10,000 bootstrap resamplings of the data. **D**. Threshold_ks differs among embryonic, larval, and pupal lethality. Distributions of bootstrapped values of Threshold_ks, a parameter that determines how quickly isolation approaches 1. Early inviabiliy (hybrid embryonic lethality) evolves faster than later inviability (hybrid pupal lethality).

### Detection of reinforcement using comparative analyses

We evaluated the possibility of different mechanisms of RI evolving through reinforcing selection. We found no difference between allopatric and sympatric lines in the magnitude of premating, NCGI, CSP, or postzygotic isolation (Figure S5). Similarly, we found no differences in embryonic, larval, or pupal inviability (in all cases Mann-Whitney U ~ 0, P > 0.5 in all cases). A parallel approach using linear models revealed a similar pattern; the geographic origin effect, whether a line was sympatric or not, did not significantly predict the strength of any RIM (Table 4). We thus find no support for the hypothesis of that reinforcing selection consistently acts on any one RIM. This does not mean that reinforcement has not played a role in the evolution of RI in some of these pairs (e.g., Matute 2010); rather, the influence of reinforcing selection is likely to be idiosyncratic and does not always influence the same RIM.

**TABLE 4.** Linear models show no difference at the strength of most types of RI between sympatric and allopatric pairs of lines. The only RIM that shows an origin effect is NCGI, whose significance is exclusively driven by the cross ♀*D. yakuba* × ♂*D. santomea* (Matute 2010). When this cross is excluded, the origin effect is not significant anymore (F_10,123_= 1.7319, P = 0.0808).

|  | Sympatric lines |  | Allopatric lines |  | Degrees of freedom<br>(numerator, denominator) | Origin effect |  |
| --- | --- | --- | --- | --- | --- | --- | --- |
|  | Mean | SD | Mean | SD |  | F-value | P-value |
| **Premating isolation** | 0.702 | 0.580 | 0.703 | 0.584 | 17,306 | 1.1462 | 0.3089 |
| **NCGI** | 0.8938 | 0.0751 | 0.8644 | 0.0966 | 11,141 | 6.9568 | $2.561 \times 10^{-9}$ |
| **CSP** | 0.1488 | 1.5807 | 0.1434 | 1.7568 | 18,88 | 0.4602 | 0.9679 |
| **Postzygotic isolation** | 0.5036 | 0.2687 | 0.5142 | 0.2824 | 12,203 | 2.2074 | 0.01262 |
| **Postzygotic isolation – embryonic lethality** | 0.0905 | 0.1909 | 0.0905 | 0.1900 | 12,203 | 1.6188 | 0.08846 |
| **Postzygotic isolation – larval lethality** | 0.2280 | 0.2231 | 0.2224 | 0.2388 | 12,203 | 1.3817 | 0.1767 |
| **Postzygotic isolation – pupal lethality** | 0.2048 | 0.1923 | 0.2282 | 0.2058 | 12,203 | 0.6689 | 0.7801 |

We also compared the magnitude of RI in two species triads characterized by an allopatric pair and a sympatric pair. The comparisons for each RIM for each triad are shown in Table S9. The majority of RIMs show no difference in magnitude between allopatric and sympatric pairs in any of the two triads. There are two notable exceptions. Behavioral isolation is stronger between *D. yakuba* and *D. teissieri* (a sympatric pair) than between *D. santomea* and *D. teissieri* (an allopatric pair). This observation has been reported before and is consistent with the action of reinforcement (Turissini et al. 2015). Second, and contrary to the expectations of reinforcing selection, *D. sechellia* and *D. melanogaster* (a mostly allopatric pair) show stronger hybrid inviability than *D. simulans* and *D. melanogaster* (a mostly sympatric pair). In particular, hybrid inviability is stronger in crosses where *D. melanogaster* is the female (*D. melanogaster* × *D. simulans*—mean = 0.593—vs. *D. melanogaster* × *D. sechellia*—mean = 0.833 —;Welch Two Sample t-test: t_7.8_ = 6.5561, P = 1.984 ×10^−4^). These results are opposite to the expectations if hybrid inviability evolved through reinforcement in this species pair. Regardless of how it was measured, our analyses indicate that reinforcement has indeed occurred in the *melanogaster* species subgroup but does not leave a consistent signature on any one RIM.

### Robustness of the pattern

We collected additional data for four more species pairs from other *Drosophila* subgroups different from *melanogaster* to address two potential issues. First, we needed to assess whether our measurements of the rate of evolution of different RIMs were robust to phylogenetic non-independence (i.e., multiple overlapping branches when only studying the *melanogaster* species subgroup). Second, when premating isolation is complete, the number of measurements of estimates of any type of postmating isolation (either PMPZ or postzygotic) is reduced. This will inflate the estimates of the rate of evolution of premating isolation (Wu 1992). To address these two potential issues, we identified species for which we could measure the magnitude of all the four types of reproductive isolation. We added these four species pairs to the data from our original study for three *melanogaster* species pairs that are phylogenetically independent (the most possible hybridizations without overlapping branches; Figure S6). In total, this gave us seven independent species pairs to compare. (Results did not change when measuring isolation between any random two overlapping branches in the *melanogaster* subgroup—accounting for the *D. simulans, D. sechellia, D. mauritiana* polytomy— for a total of seven species pairs, or when excluding this the species in this polytomy altogether— for a total of six species pairs.) Similar to what we observed in the *melanogaster* subgroup-only analysis, premating isolation is the fastest RIM to evolve, followed by conspecific sperm precedence, non-competitive gametic isolation, and finally postzygotic isolation (Figure 6). Also, as observed in the *melanogaster* species group, the four possible pairwise comparisons between the bootstrapped distributions of the rates of RIMs differed from each other (Table S10). These results indicate that the ranking of the rates of evolution of the four RIMs is not exclusive to the *melanogaster* species subgroup of *Drosophila*, and instead is a more general pattern that might pertain the whole *Drosophila* genus.

**FIGURE 6.**
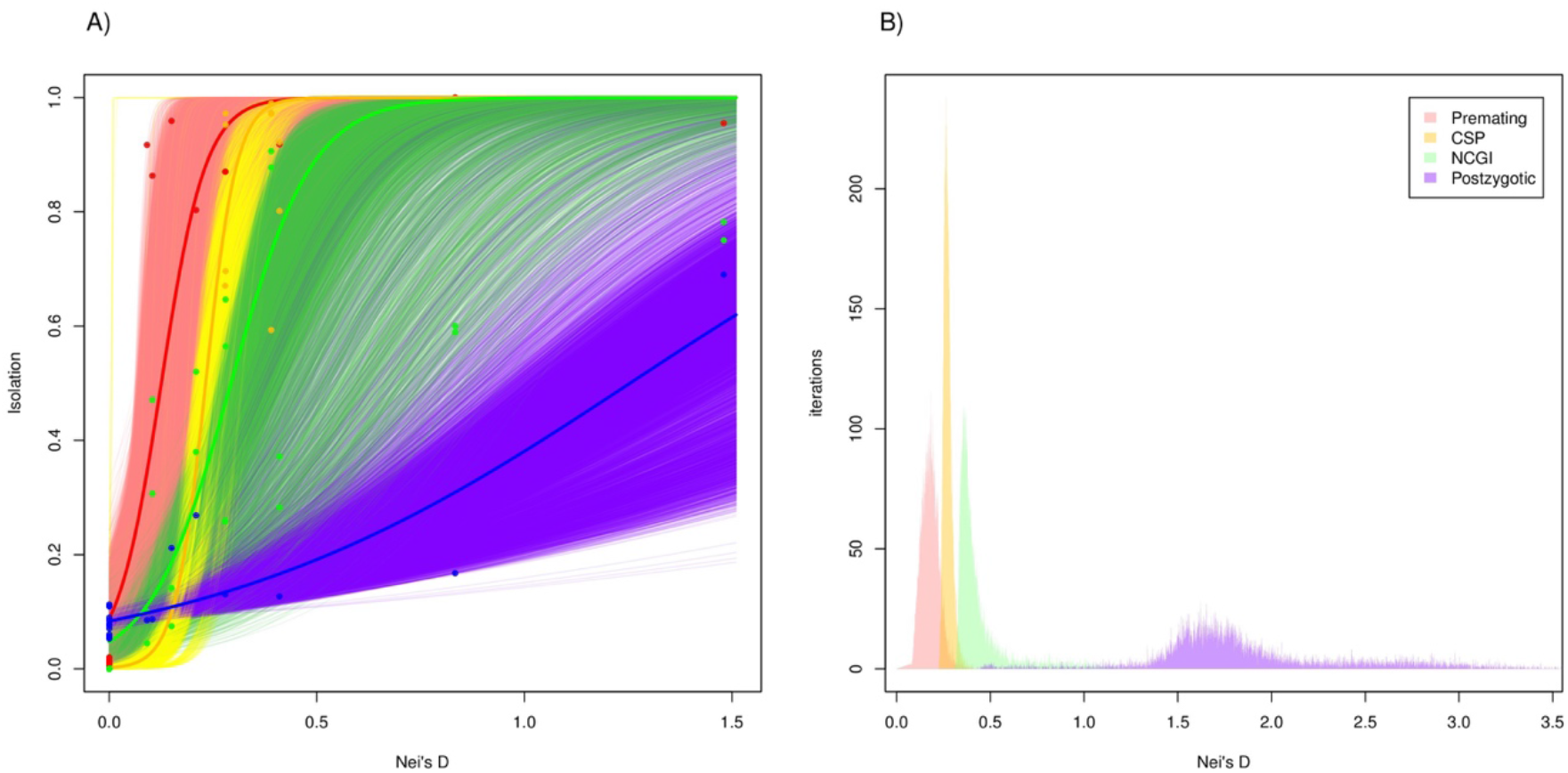
Premating, CSP, NCGI, and postzygotic isolation show a strong phylogenetic signal across the *Drosophila* genus. To account for the possibility of phylogenetic non-independence in the *melanogaster* species subgroup, we subsampled phylogenetically independent crosses and added a pair of species from other four *Drosophila* clades. Proxies of RI are similar to the ones shown in Figure 1. Premating isolation (red), conspecific sperm precedence (CSP, yellow), non-competitive gametic isolation (NCGI, green), and postzygotic isolation (blue) were regressed against phylogenetic distance (K_s_ between species and π_s_ within species). **A**. As observed in the *melanogaster* species subgroup, the four types of isolation increase with genetic distance, and premating isolation evolves faster than hybrid inviability. The thick red, yellow, green, and blue lines represent fitted logistic regressions for the premating and postzygotic data respectively. The thinner lines of each of the four colors are the regressions for each of 10,000 bootstrap resamplings of the data. **B**. Premating, NCGI, CSP and postzygotic isolation (hybrid inviability) evolve at different rates in the *Drosophila* genus in a set of phylogenetically independent species pairs. Threshold_Ks indicates the Ks value for which a given RI barrier achieves a value of 0.95.

### No signature of generalized positive selection in genes potentially involved in RI

Finally, we tested whether any GO term associated with any of six types of RI (premating, NCGI, CSP, embryonic development, larval development, and pupal development) showed a signature of accelerated molecular evolution compared to the rest of the genome. The median K_A_/ K_S_ for the genome was 0.06595. All GO terms showed a median K_A_/ K_S_ similar to the genome wide median (Tables S11 and S12). These results suggest that selection at the molecular level is not pervasive at any particular component (i.e., GO term) of RI.

## DISCUSSION

Little is known regarding the rate of evolution of postmating prezygotic (PMPZ) isolation in animals. Studies on plants have found that PMPZ and postzygotic isolation evolve at a similar rate in at least three plant genera. Unlike plant studies, most studies evaluating the rate of accumulation of reproductive isolation in animals have a common limitation: they have not looked at the rate of evolution of PMPZ barriers. We thus measured the rate of evolution of such barriers in *Drosophila* species pairs while assessing the magnitude of premating and postzygotic isolation in the same crosses. This makes our study the first to measure the magnitude of PMPZ isolation, compare it with other RIMs, and explicitly test its rates of evolution in animals. Our results have implications for our understanding of three large topics in speciation: *i*) the evolution of PMPZ barriers, *ii*) the role of PMPZ isolation on speciation via reinforcement, and iii) the evolution of postzygotic isolation. We discuss each of these topics as follows. We also present a series of caveats and general conclusions of our analyses.

### The evolution of PMPZ barriers

PMPZ RIMs, both non-competitive and CSP have been hypothesized to evolve through the influence of sexual selection (Birkhead and Pizzari 2002) and natural selection (Knowles et al. 2004). The female × male interactions that underlie NCGI can be interpreted as discrimination against heterospecific sperm by an inseminated female and are thus sexually selected (Price et al. 2001; Birkhead and Pizzari 2002; Fricke and Arnqvist 2004). Phenotypes involved in fertilization success and sperm morphology have both been found to show fast rates of change across species (Pitnick et al. 1999, Presgraves et al. 1999, Byrne et al. 2003, Manier et al. 2013). These phenotypes also evolve rapidly within and between species due to the constant influence of antagonistic sexual conflict (e.g., Knowles and Markow 2001, Comeault et al. 2016, reviewed in Pizzari and Snook 2003). Proteins involved in fertilization, gametic fusion, and stimulation of oviposition show signatures of positive selection across all taxa for which this signature has been systematically sought (e.g., Lee et al. 1995, Swanson et al. 2001, Galindo et al. 2003, Marshall et al. 2011, Harrison et al. 2015). This accelerated evolution at the phenotypic and molecular level are consistent with evolution of these interactions via either sexual or natural selection.

The second type of PMPZ barrier we examined, CSP, is also affected by sexual selection. CSP (and its plant analogous conspecific pollen precedence) is ubiquitous and has been uncovered in a variety of organisms (references in Yeates et al. 2013). CSP is the aggregate of the three possible interactions between female reproductive tract, conspecific sperm, and heterospecific sperm. These interactions create the grounds for reproductive incompatibilities due to sexual antagonism between sexes of different species, sexual antagonism between sexes of the same species, and sexual competition between sperm of different species (Howard 1999; Simmons 2005). Our results indicate that at least one of these interactions scales up with genetic divergence which leads to stronger CSP in divergent species.

Since premating and PMPZ traits usually co-occur with premating isolation in organisms with internal fertilization (such as *Drosophila*), the rapid evolution of PMPZ traits poses a conundrum: how can sexual selection or reinforcing selection influence the evolution of PMPZ barriers in the presence of premating isolation? First, premating isolation is often not complete, giving natural and sexual selection the opportunity to drive the evolution of PMPZ traits. Second, in organisms with internal fertilization, the evolution of PMPZ traits might be accelerated in instances where “the wallflower effect” applies (Kokko and Mappes 2005); females might adaptively lower their sexual preferences for males of high condition (conspecifics) if they are only exposed to males of low condition (heterospecifics). Similar models (Wilson and Hedrick 1982) and experimental measurements (Matute 2014) have shown that females might mate with heterospecifics if mates are rare. These instances where premating isolation is not an effective RIM might favor the accelerated evolution of PMPZ. The formal test of this hypothesis will require measuring PMPZ isolation in sister populations (or species) that differ in their strength of premating isolation.

### The role of PMPZ isolation on speciation via reinforcement

Our results are also important in the context of reinforcement. Two different approaches have been historically used to detect reinforcement: detecting reproductive character displacement in areas of secondary contact, and detecting the phylogenetic signal in sympatric species pairs. We used a modified version of the former approach. The comparison of allopatric and sympatric lines from the same species detects reinforcement at recent scales (after secondary contact). On the other hand, the phylogenetic comparison of the magnitude of RIMs detects reinforcement at deeper scales of divergence. We found no new evidence for cases of reinforcement besides the already reported influence of reinforcing selection in NCGI in the *D. yakuba/D. santomea* hybrid zone (Matute 2010), and the phylogenetic signature of reinforcement at behavioral isolation in the *D. teissieri/D. yakuba* species pair (Turissini et al. 2015). It is possible that our experiment has little power to detect differences because all RIMs are already strong and the influence of sympatry is minimal compared to the amount of divergence that has already occurred between species.

A surprising result comes from the comparisons between the pairs *D. simulans/D. melanogaster* and *D. sechellia/D. melanogaster*. The latter pair shows extremely high hybrid inviability compared to the former pair. Given that *D. melanogaster* and *D. sechellia* are largely allopatric while *D. melanogaster* and *D. simulans* are largely sympatric, this pattern goes against the expectation of evolution of RI by reinforcing selection. The reasons behind such stark difference in the magnitude of RI remain unknown but we can formulate two possibilities. The first one is that *D. sechellia* has accumulated more hybrid incompatibilities due to the extreme bottlenecks to which it has been subjected during its evolutionary history. The second one is that *D. melanogaster* and *D. simulans* have had more chance to interbreed in the distant past thus purging hybrid incompatibilities that still separate *D. sechellia* and *D. melanogaster*. More research on the demographic history of these species, as well as the effect of different demographic events on the accumulation of incompatibilities is needed before addressing the reasons for this difference.

The influence of PMPZ isolation on speciation by reinforcement remains largely unstudied. Reinforcement is traditionally viewed as the process of strengthening premating isolation driven by selection against unfit hybrids. However, PMPZ acts earlier in the reproductive cycle and has a faster rate of evolution compared to hybrid inviability and hybrid female sterility. Therefore, deleterious and costly PMPZ incompatibilities, such as reduced female fertility after heterospecific matings, might lead to the evolution of behavioral barriers in the same manner that postzygotic costs lead to premating isolation during conventional reinforcement (Harrison 1993; Servedio 2001). Even though we did not perform a formal comparison between the rates of evolution of PMPZ barriers and hybrid male sterility, the two types of RIMs seem to evolve at roughly the same rate. Thus, both PMPZ and hybrid male sterility might be equally important in inducing the evolution of premating isolation via reinforcement. Currently, the evidence that reproductive interference (excluding the production of unfit hybrids) might be costly is currently scattered and has been circumscribed to premating interactions (e.g., reproductive character displacement caused by noisy neighbors, Mullen and Andres 2005 but see Matute 2015).

Theoretical arguments have also suggested that CSP can hamper the evolution of premating isolation by reinforcement because if CSP is complete, and a female has the chance to mate with multiple males, then no hybrids are likely to be produced if one of those males is a conspecific (Lorch and Servedo 2007). A similar argument can also be made about non-competitive gametic isolation. If NCGI is strong, then no hybrids will be produced after heterospecific matings and if a female remates with a conspecific, then most of her progeny will be pure species and fit. In both these cases, there will be no cost to hybridization and no incentive to strengthen premating isolation. This hypothesis yields a clear prediction: reinforcement of premating isolation should be more rare in clades where PMPZ isolation is strong. In spite of its straightforwardness, it might be premature to test this hypothesis because bona fide cases of reinforcement and of gametic isolation are still rare.

Conversely, postzygotic isolation might also lead to the evolution of PMPZ traits in the same manner that it leads to the evolution of premating isolation. This is obvious in aquatic organisms that spawn in open waters but it has been more controversial in animals with internal fertilization. Overall, reinforcement should affect the evolution of any trait that minimizes maternal investment on an unfit hybrid (Coyne 1974; Servedio and Noor 2003). Two examples show that reinforcement can indeed lead to the evolution of PMPZ traits. In the case of *Drosophila yakuba*, females from the hybrid zone with *D. santomea* show stronger non competitive gametic isolation than females from areas where *D. yakuba* is not present (Matute 2010, Comeault et al. 2016). Similarly, CSP in *D. pseudoobscura* is stronger in areas of sympatry with *D. persimilis* (Castillo and Moyle 2016). Both patterns of reproductive character displacement are highly suggestive of reinforcement and indicate that reinforcement of PMPZ barriers might not be a rare instance even in animals with internal fertilization.

### The evolution of postzygotic isolation

Our measurements also allowed us to discern which developmental stage was most affected by hybrid defects. Of the three possible transitions (embryo to larva, larva to pupa, and pupa to adult). We found that embryonic lethality arises first and evolves faster than larval or pupal lethality. A possible explanation for this result is that more genes are involved in embryogenesis than in other developmental stages, which seems to be the case from multispecies analyses of gene expression (Graveley et al. 2011). Hybrid incompatibilities would, therefore, have more potential targets at the embryonic stage. The GO category ‘embryo development’ indeed contains more genes (740) than ‘larvae and prepupal development’ (177), which in turn contains more genes than ‘pupal development’ (17). The identification of alleles involved in hybrid inviability has yielded mixed support for this hypothesis. Mapping of *X*-linked dominant factors in three *Drosophila* interspecific hybrids revealed that the majority of *X*-linked alleles in *mel/san* hybrids cause embryonic inviability. In the other two hybrids (*mel/sim* and *mel/mau*), however, no embryonic lethality alleles were found (Matute and Gavin-Smyth 2014). Similarly, in *mel/sim* hybrid males the vast majority of alleles involved in hybrid inviability cause postembryonic and not embryonic lethality (Presgraves 2003). We found that genes associated to all RIMs (clustered by GO terms and including different developmental transitions) show average rates of molecular evolution comparable with the rest of the genome. This is important because a nontrivial fraction of genes found to cause hybrid inviability and hybrid sterility have signatures of positive selection (e.g., Presgraves et al. 2003, Tang and Presgraves 2009 reviewed in Coyne and Orr 2004 and Nosil and Schluter 2011). Yet, we find no evidence for a consistent signature of positive selection at a particular RIM. Even though these GO analyses have important caveats (reviewed in Rhee et al. 2008) and we cannot rule out that strong selection has occurred at *cis*-regulatory elements, our results indicate that broadly speaking, there is no support for the idea that genes involved in RIMs that act early in reproduction evolve faster than genes involved in RIMs that act later on.

Finally, we also saw complete hybrid male sterility for all interspecific crosses where males were viable (Ks higher than ~0.05), and hybrid females were consistently sterile when Ks exceeded ~0.10. It is worth noting how dramatically full hybrid sterility can arise when compared to the other forms of isolation we investigated. Since hybrid inviability is the slowest barrier to evolve, our results are consistent with the idea that hybrid sterility evolves faster than hybrid inviability in both sexes (Wu 1992), an idea that had remained formally untested because indexes of postzygotic isolation usually conflate sterility and inviability.

### Caveats

Our study is not devoid of caveats. The first one pertains to how much reinforcement can affect different types of reproductive barriers. Since reinforcement is thought to affect prezygotic isolation more commonly, then it is possible that reinforcing selection has led to an increase in the rate of evolution in premating, NCGI, an CSP isolation. This in turn would lead to an inflation of our estimated rate of evolution of these three RI barriers. Even though we compared allopatric and sympatric populations from eight species pairs, reinforcement might act at deeper levels of divergence that do not involve contemporary coexistence. An obvious research avenue is to test whether sympatric species evolve PMPZ mechanisms faster than allopatric pairs. This approach is not trivial as the range of species contracts and expands along their history making the distinction between allopatric and sympatric a gray area. Our dataset does not allow us to split between currently allopatric and currently sympatric pairs because species from the *melanogaster* subgroup are largely sympatric (Lachaise et al. 1988) and an expanded dataset will be necessary to address the importance of reinforcement.

A second bias is that when an early acting barrier is complete, we cannot measure the magnitude of later acting barriers. This introduces a bias that might inflate the rates of evolution of premating isolation because there will be more measurements of strong premating isolation than of postmating isolation. Since we tried all possible crosses, and for one analysis only included species for which we had measured the magnitude of the four types of RI barriers, this is not a concerning caveat.

### Conclusions

In general, we find that both PMPZ barriers evolve faster than postzygotic isolation, but slower than premating behavioral mechanisms. These results indicate that there is a qualitative difference between the rate of evolution of PMPZ barriers in *Drosophila* and plants; in the latter PMPZ and postzygotic barriers evolve at similar rates (Moyle et al. 2004, Jewell et al. 2012). A possible explanation for this dichotomy is that in *Drosophila*, mate choice is a primary source of intrinsic isolation, while in plants the main source of intrinsic isolation might occur as pollen reaches the stigma. More research in different plant and animal taxa are needed to establish whether this difference is real or whether it is the byproduct of sparse taxonomic sampling. Similarly, and even though there is evidence for the existence of PMPZ in fungi (Turner et al. 2010, 2011) the rate of evolution of premating and postmating isolation in this group and other eukaryotes remains largely unexplored (but see Gourbière and Mallet 2010; Giraud and Gourbiere 2012) and are sorely needed to understand what biological features drive the origin of new species.

Across metazoans, *Drosophila* has been one of the premier model systems for studying the evolution of reproductive isolation which in turn has provided support for several hypotheses such as the existence of reinforcement (Coyne and Orr 1989; Coyne and Orr 1997; Nosil 2013), the relative rate of evolution of RI (Yukilevich 2012; Rabosky and Matute 2013), and the role of ecology in speciation (Funk et al. 2006; Turelli et al. 2014). Overall, the fast accumulation of PMPZ isolation indicates that they are likely to be driven by selection, either sexual or natural. The integration of our results show that the earlier a barrier acted during the reproductive/developmental process, the faster its rate of accumulation over time. PMPZ isolation accumulates quickly in *Drosophila* thus indicating that this type or RI might be an important source of isolation in promoting the evolution of new species but also in keeping them apart.

## MATERIALS AND METHODS

### *Drosophila melanogaster* subgroup: Species and stocks

All wild-type stocks are described in Table S13. Briefly, for all genetic crosses we used synthetic stocks (i.e., outbred stocks derived from a combination of isofemale lines) with the exception of *Drosophila erecta*. Stocks from *D. santomea, D. yakuba, D. teisseiri, D. orena, D. sechellia, D. simulans* and *D. melanogaster* were collected by DRM (Table S13). Stocks from these species were kept in large numbers (>200 flies) since their creation. *Drosophila erecta* was purchased at the San Diego Stock Center (Stock number: 14021-0224.00). All lines were reared on standard cornmeal/Karo/agar medium at 24°C under a 12 h light/dark cycle in 100mL bottles. Adults were allowed to oviposit for one week and after that time they were cleared from the bottles. We added 1mL of propionic acid (0.5% V/V solution to the vials and provided a pupation substrate to the vial (Kimberly Clark, Kimwipes Delicate Task; Irving, TX). At least 10 bottles of each species were kept in parallel to guarantee the collection of large numbers of virgins.

To measure conspecific sperm precedence, we also used mutants from each of eight of the species (with the exception of *D. orena*, see below). All mutants were raised in identical conditions to the wild-type stocks.

### Virgin collection

Pure species males and females of each species were collected as virgins within 8 hours of eclosion under CO_2_ anesthesia and kept for three days in single-sex groups of 20 flies in 30mL, corn meal food-containing vials. Flies were kept at 24°C under a 12 h light/dark cycle. On day four, we assessed whether there were larvae in the media. If the inspection revealed any progeny, the vial was discarded.

### Premating isolation: Insemination rates

We measured premating isolation as the number of females that did not accept heterospecific males when housed together in no-choice experiments for 24 hours. Two hundred females (i.e., individuals pooled from 10 virgin vials) were housed with 200 males either from the same species or from a different species. Females and males were housed together for 24 hours. After that time, females were anesthetized with CO_2_ and males were discarded. We dissected all the females and extracted their reproductive tract (spermathecae, seminal receptacles, and uterus) and placed it in chilled (4°C) Ringer’s solution. We assessed whether the female carried any sperm, either dead or alive anywhere in their reproductive tract. We used the proportion of females inseminated in the *en masse* matings in each bottle (see ‘Insemination rates’) and calculated a proxy of the strength of premating isolation:

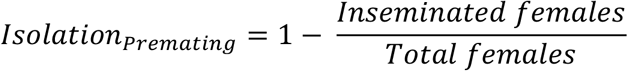

Five batches (bottles) per cross (each with 100 females) were counted.

We assessed whether there was heterogeneity in insemination rates among conspecific matings. We counted how many females were inseminated in 5 replicates. To detect heterogeneity, we fit a linear model in which the proportion of inseminated females in these conspecific crosses was the response and the cross (i.e., species) was the only factor.

Next, we studied whether there was heterogeneity in premating isolation among interspecific crosses. We fit two linear regressions to analyze the data. First, to assess if any particular combination of species was more prone to mating than others, we fit a linear regression in which *Isolation_Premating_* in each bottle was the response, and the identity of the cross was the only fixed effect. There were five replicates per species for a total of 360 bottles. Second, we analyzed whether any type of female (and male) were more prone to hybridize with other species. To do so, we used the same data set but fit a factorial model in which *Isolation_Premating_* in each bottle was the response and the identity of the female and that of the male were fixed effects. We also included an interaction term. All statistical analyses were carried out using the package “stats” in R (function: lm; R Core Team 2016).

### PMPZ isolation: non-competitive gametic isolation (NCGI)

We next measured gametic incompatibilities between females and males from different species in single matings, namely, the inability of a male to induce a female to lay eggs (Price et al. 2001; Matute 2010b; Marshall and DiRienzo 2012). We watched single heterospecific and conspecific pairs for 8 hours and kept the females that mated successfully for each of the 81 possible hybrid crosses (72 heterospecific + 9 conspecific). We repeated this approach until we collected at least five females from each of the heterospecific and conspecific crosses. We kept all females who mated (either to con- or heterospecific males) to measure gametic isolation. To prevent females from re-mating, males were removed from the vial by aspiration after mating. Each mated female was allowed to oviposit for 24 h in a vial. The female was then transferred to a fresh vial, and the total number of eggs were subsequently counted daily for 10 days. At least five females were scored for each cross.

I_g_, an index of PMPZ isolation which was calculated as:

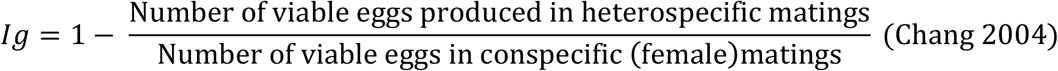

I_g_ values were compared across crosses using a linear model in which cross was the only fixed factor.

### PMPZ isolation: competitive gametic isolation

#### CSP indexes

We also scored how much hybrid progeny a female produces after mating with two males: a heterospecific, and a conspecific male. To do so, we used a combination of mutants to differentiate between hybrid and pure species progeny in crosses that involved more than one male. Traditionally, CSP is measured as P2, the proportion of progeny sired by the second male in double matings (Boorman and Parker 1976; Chang 2004). Nonetheless, this measurement conflates two important biological aspects. First, second males have an advantage over first males (regardless of their genotype) and in conspecific crosses, they invariably sire more progeny than first males. Second, conspecifics sperm might indeed have an advantage over heterospecific sperm (true conspecific sperm precedence). We propose to quantify CSP as the proportion of progeny sired by a heterospecific male by a doubly mated female respective to a conspecific mating that mated to two conspecific males. To account by the fact that the second male usually has an advantage over the first male, we propose to do normalizations taking into account the order of mating. Matings with two males can occur in two different orders. In a heterospecific/conspecific mating, we counted the number of hybrid progeny (HC_H_) and the number of conspecific progeny (HC_c_). In crosses where the heterospecific male was mated first followed by a conspecific male, the indexes took the form HC_H1_ and HC_C2_. In crosses were the conspecific male was first and was followed by a heterospecific male, the indexes took the form HC_C1_ (i.e., the progeny sired by the heterospecific male in females mated to a heterospecific male and then to a conspecific male) and HC_H2_ (i.e., the progeny sired by the heterospecific male in females mated to a conspecific male and then to a heterospecific male). In conspecific/conspecific matings, we counted the progeny sired by a wildtype and a mutant stock of the same species. This yielded two quantities: the progeny sired by the first male, CC_C1_ (i.e., the progeny sired by the first male in females mated to two conspecific males), and the progeny sired by the second male, CC_C2_ (i.e., the progeny sired by the second male in females mated to two conspecific males)

These quantities were then incorporated into two indexes of conspecific sperm precedence, one for crosses when heterospecific males were the first to mate, and one for crosses where conspecific males were first. The two indexes followed the following form:

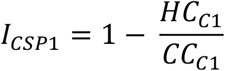

and

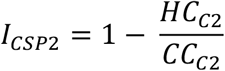

There was only true CSP if both *I_CSP1_* and *I_CSP2_* indexes were low.

*Mutant stocks:* In order to quantify CC_C1_and CC_C2_ we needed to obtain females that mated to conspecific males twice and be able to distinguish between the progeny sired by each father. To do so, we needed mutant stocks that could be visually recognized from the wild type. We described the mutants for eight species in the melanogaster species subgroup (no mutants were available for *D. orena)*.

i. *Drosophila melanogaster: D. melanogaster yellow* white (*mel^y1 w1^*) males and females were derived from a stock purchase at the Bloomington Stock Center (Stock number: 1495).
ii. *Drosophila mauritiana: D. mauritiana yellow* (*mau^y1 w1 f1^*) males and females were derived from a stock purchase at the San Diego Stock Center (Stock number: 14021–0241.55).
iii. *Drosophila yakuba: D. yakuba yellow* (*yak^y^*) males and females were derived from a stock originally collected in the Täi Forest (Liberia) in 1998.
iv. *Drosophila teissieri: D. teissieri yellow* (*tei^y^*)was isolated from a line collected in Bioko (2009). Both, *yak^y^* and *tei^y^*, are a fully recessive body color mutation identical to that on the *D. melanogaster X* chromosome (Llopart et al. 2002).
v. *Drosophila erecta: D. erecta yellow* (*ere^y^*) also has a body color mutation. Whether this yellow mutation complements *mel^y^* remains unknown.
vi. *Drosophila santomea: D. santomea white* (*san^w^*) males and females were derived from the STO.4 isofemale line, originally collected on São Tomé in 1998. All the white-eyed mutations described in this contribution are fully recessive eye color mutations orthologous (i.e., fail to complement) to white eyed mutations on the *D. yakuba* and/or *D. melanogaster X* chromosomes.
vii. *Drosophila sechellia: D. sechellia white* (*sech^w^*) was isolated from a recently collected line in Denis island.
viii. *Drosophila simulans: D. simulans white* (*sim^w^*) was donated by D.C. Presgraves.

#### Measuring CSP

First matings were watched as described above (Section ‘Premating isolation: Insemination rates’). Mated females were then separated from the males and housed in groups of 1 to 5 females. On the morning of day 4, females were individually transferred to a new vial with cornmeal food. The male to be mated was also transferred to the vial by aspiration. We observed up to 300 individual matings at the same time. Second matings were allowed to proceed for 16 hours. To identify true CSP, we attempted to measure the magnitude of sperm precedence in two directions of the cross. Crosses that involved a conspecific male first and an interspecific male second are challenging and in some cases estimates involved only a few measurements. The sample sizes of each mating are shown in Table S3. Once doubly mated females were obtained, we removed the male from the vial, kept the females and tended their progeny. Females were transferred to a new vial every seven days until they died. Vial tending and fly husbandry were done as described immediately below. I_CSP_ were compared using a linear model with the identity of the cross and the direction (i.e., what male was mated first) as the two fixed factors.

### Postzygotic isolation: Hybrid inviability

Finally, we measured viability in F1 hybrids. For each interspecific and conspecific species pair, we calculated overall F1 inviability and three components of inviability: embryonic lethality (death during the embryo-to-L1 transition), larval lethality (death during the L1 larvae-to-pupa transition), and pupal lethality (death during the pupa-to-adult transition). We collected virgin males and females as described above (See ‘Virgin collection’). On the morning of day four after collection, we placed forty males and twenty females together at room temperature (21°−23°C) to mate *en masse* on corn meal media. We set up 50 crosses per species pair for a total of 4,050 crosses (81 crosses × 50 replicates). Vials were inspected every five days to assess the presence of larvae and/or dead embryos. We transferred all the pure species adults to a new vial (without anesthesia) every ten days. This procedure was repeated until the cross stopped producing progeny. L1 larvae were allowed to feed on an apple-agar plate and were tended daily. Once L2 larvae were observed, we added a solution of 0.05% propionic acid and a KimWipe (Kimberly Clark, Kimwipes Delicate Task, Roswell, GA) to the vial. All hybrids were collected and counted using CO_2_ anesthesia. To measure the magnitude of hybrid inviability, we transferred the adults from the vials that produced progeny to an oviposition cage with apple juice media and yeast. The plates were inspected every 48 hours for the presence of viable eggs. We transferred all the pure species adults to a new vial (without anesthesia) every ten days. In order to maximize the lifespan of the parents, we kept all the vials lying on their sides. We repeated this procedure until we obtained five cages that produced hybrid progeny for the crosses for which we could obtain inseminated females.

We partitioned overall inviability into three components by comparing the number of individuals that entered a developmental stage (Total) to the number that survived it (Successes) using the equations shown in Table 1. The proportions were then transformed to a logistic index following the form:

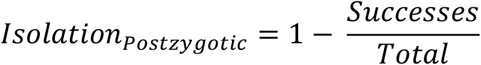

Overall inviability was based on the number of individuals that died during development (from embryo to adulthood). For a global estimate of inviability, we counted the total number of viable eggs and the number that developed into adults. To quantify embryonic inviability, we counted the total number of embryos defined as the total number of egg cases (successes) plus the number of dead embryos (brown eggs). This procedure was applied to 69 interspecific crosses (the exception being crosses between females from the *simulans* clade *and D. melanogaster* males; See below). To quantify larval lethality, we counted the number of egg cases (Total) and pupae (Successes) in each vial. If larvae pupated on the food media, the vial was discarded. Finally, to quantify pupal viability we counted the number of pupae (Total) and adults (Successes). We quantified lethality for at least five replicates per cross and summed the results. The number of replicates per cross is listed in Table S14.

For embryonic lethality we assessed the robustness of our two proxies: egg cases for viable eggs, and brown eggs for dead embryos. First, we studied how robust was the proxy of empty cases for the number of live embryos. As larvae feed, they churn the food and egg cases can disappear from the surface. As a result, counts of the number of egg cases + brown eggs (dead embryos) were sometimes less than the initial egg count. To account for this missing data, we inferred the number of missing dead embryos from the consolidated data from the replicates using the formula: missing dead embryos = (dead embryos / total eggs) × missing embryos. This estimate was rounded to the nearest integer and added to the number of dead embryos. The number of egg cases was likewise adjusted by adding the difference between missing and missing dead embryos. Accounting for the missing data does not affect our results (Figure S7).

Second, we adjusted the estimates of dead embryos. Three crosses have shown extensive embryonic mortality before the zygote stage is achieved: ♀ *D. simulans* × ♂*D. melanogaster*, ♀ *D. sechellia* × ♂*D. melanogaster*, and ♀ *D. mauritiana* × ♂*D. melanogaster* (Sawamura et al. 1993). We focused on these three crosses because no other cross in the *melanogaster* species group exhibit the same phenomenon. In these three interspecific crosses, brown eggs are rare, as the female diploid embryos that do not develop do not achieve the status of zygote. To measure the magnitude of female embryonic lethality in these three crosses, we collected one-hundred 72-hour embryos from each cross. From each of these embryos, we extracted DNA using the QIAamp DNA Micro Kit (Qiagen, Chatsworth, CA, USA) following the manufacturer's instructions and amplified the *yellow* locus of *D. melanogaster* by PCR using the primers mel_y_F: 5’ CGGCTCCCTTGGCCACTTTA3' and mel_y_R: 5' CGGGCATTCACATAAGTTTTTAACC 3’. Both primers include sites private to *D. melanogaster* in positions 20 and 25, respectively, and do not amplify in *D. simulans*. The primers amplify a 412 bp fragment (Tm = 59.4°C). The presence of this locus meant an embryo had been fertilized, and its absence meant the embryo was unfertilized. We used the primers control_y_F:: 5' CTGACTTGGATTATTCAGATACTAATTTC3' and control_y_R:: 5' CTACATTGCCTGAATTGGCG3' as a positive amplification control. These primers amplify a PCR product of 267 bp (Tm = 56.0 °C). PCR conditions were identical to those described elsewhere (Matute and Ayroles 2014). Amplicons were run in a 2% agarose gel and visualized using ethidium bromide and UV. None of the newly described hybrids produces only adult males, so there were no additional cases of female early embryonic lethality and thus no need to correct these estimates.

The proportion of dead embryos in an oviposition cage was calculated by multiplying the total number of eggs in the oviposition cage with the average proportion of embryos that were diploid (i.e., carried the *D. melanogaster yellow* locus) and did not hatch. To detect heterogeneity among crosses, we used the ‘lm’ function in the ‘stats’ package in R to fit linear model where the strength of hybrid inviability was the response and the identity of the cross was the only fixed effect. Since there were three different types of hybrid inviability (one for each developmental transition) and a life-long estimate of inviability, there were four linear models.

Correlations between viability at different development transitions were calculated using a Pearson's product-moment correlation (R package ‘Stats’: function ‘cor.test’; ρ). Critical P-value for significance was 0.01 to account for multiple comparisons (three comparisons).

To measure sex-specific hybrid viability we took advantage of the existence of *yellow* mutant stocks mutant stocks for which we could differentiate between males and females early in development. In crosses between a *yellow*-null carrying mother and a wild-type father, female progeny is heterozygote (*y*^+^*y*^−^) and larvae have black mouthparts. Male progeny will be hemizygous for the *y*-null and their mouthparts will be brown. Four of five *yellow*-used in this experiment stocks are described in above (i,ii,iii,iv in section ‘PMPZ isolation: competitive gametic isolation, Mutant stocks’). For these experiments, we also used an additional stock, *Drosophila simulans yellow*^−^, which was donated by J.A. Coyne. This *yellow* mutation does not complement *mel^y^*.

### Postzygotic isolation: Fertility assessments

For all pure-species and interspecific crosses, we assessed whether their progeny were fertile or sterile. The protocol was similar for both sexes: we extracted and dissected their gonads to look for the production of gametes. In the case of female hybrids, we looked at the presence of ovarioles in the ovaries; females with ovarioles were classified as fertile. We used this binary scale to avoid the significant effects that environment has on ovariole number (Wayne and Mackay 1998, Wayne et al. 2006). In the case of male hybrids, testes were dissected, mounted in Ringer’s solution and squashed to assess for the presence of motile sperm. We scored 100 individuals per cross per sex. We measured isolation separately for each sex as:

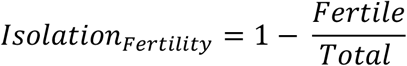

Hybrids that were then thought to be sterile were then housed with pure species individuals from the opposite sex (both parentals) to make sure our assessment of fertility was qualitatively adequate. Obviously, individuals that had been dissected and scored for fertility could not be mated; from a single cross we dissected half of the progeny and kept at least 50 of them to do en masse matings. Hybrid females were housed with males of the two species following (Turissini et al. 2015); hybrid males were housed with virgin females from both species. For both sexes, we assessed whether the crosses produced progeny until hybrid individuals were dead. Male sterility, and to a lesser extent hybrid female inviability were binomial traits that showed separation (i.e., all crosses above a particular Ks produce only sterile progeny) and for that reason these two traits were not directly compared with premating, NCGI, CSP or hybrid inviability which showed a continuous trait distribution.

The bootstrapped distributions of hybrid male sterility and hybrid male inviability were compared using a two-tailed Wilcoxon sign test (R package ‘stats’, function ‘wilcox.test’). Hybrid female sterility and hybrid female inviability were compared using one-sample t-test (R package ‘stats’, function, ‘t.test’) where the distribution of bootstrapped Ks values where female hybrid sterility achieved 95% was compared to the earliest fixed value of Ks at which hybrid female inviability reached the same level: Ks=0.25; See results).

### Genome sequencing

#### DNA extraction

DNA was extracted from single female flies using the QIAamp DNA Micro Kit (Qiagen, Chatsworth, CA, USA). We followed the manufacturer’s instruction using cut pipette tips to avoid shearing the DNA. This protocol can yield up to ~50ng of DNA per fly per extraction.

#### Library construction

For short read sequencing, we constructed libraries following two options. 54 libraries were built using the Kappa protocol for TrueSeq at the sequencing facility of the University of North Carolina, Chapel Hill. For these libraries, DNA was sheared by sonication. Briefly ~10 ug of DNA were sonicated with a Covaris S220 to 160 bp mean size (120–200 bp range) with the program: 10% duty cycle; intensity 5; 100 cycles per burst; 6 cycles of 60 seconds in frequency sweeping mode The second type of libraries were Nextera libraries which were built at the sequencing facility of the University of Illinois, Urbana-Champaign. For this type of library, DNA was segmented using Nextera kits which uses proprietary transposases to fragment DNA. Libraries were built following standard protocols.

#### Sequencing

Lines were sequenced in a HiSeq 2000 machine and were a mixture between single end and paired end sequencing. Table S15 indicates the sequencing type and coverage for each line. Libraries were pooled and 6 individuals were sequenced per lane. The HiSeq 2000 machine was run with chemistry v3.0 and using the 2 × 100 bp paired-end read mode and original chemistry from Illumina following the manufacturer's instructions. To assess the quality of the individual reads, the initial data analysis was analyzed using the HiSeq Control Software 2.0.5 in combination with RTA 1.17.20.0 (real time analysis) performed the initial image analysis and base calling. CASAVA-1.8.2 generated and reported run statistics of each of the final FASTQ files. Resulting reads ranged from 100bp or 150bp and the target average coverage for each line was 30X. The actual coverage for each line is shown in Table S15.

#### Public data

We accessed and used two additional sources of genomic data. We downloaded available raw reads (FASTQ files) from NCBI and mapped them to the corresponding reference genome (see below). Additionally, we downloaded *D. melanogaster* sequences from the nexus sequencing project (Lack et al. 2015).

#### Read mapping and variant calling

Reads were mapped using bwa version 0.7.12 (Li and Durbin 2010). *Drosophila yakuba, D. teissieri*, and *D. santomea* reads were mapped to the *D. yakuba* genome version 1.04 (Drosophila 12 Genomes Consortium et al. 2007), *D. simulans, D. sechellia*, and *D. mauritiana* reads were mapped to the *D. simulans w*^501^ genome (Hu et al. 2013), and *D. orena* reads were mapped to the *D. erecta* genome (Drosophila 12 Genomes Consortium et al. 2007). Bam files were merged using Samtools version 0.1.19 (Li et al. 2009). Indels were identified and reads were locally remapped in the merged bam files using the GATK version 3.2–2 RealignerTargetCreator and IndelRealigner functions (McKenna et al. 2010; DePristo et al. 2011). SNP genotyping was done independently for the *D. yakuba* clade, *D. simulans* clade, and *D. orena* using GATK UnifiedGenotyper with the parameter het = 0.01. The following filters were applied to the resulting vcf file: QD = 2.0, FS_filter = 60.0, MQ_filter = 30.0, MQ_Rank_Sum_filter = −12.5, and Read_Pos_Rank_Sum_filter = −8.0. Sequences were created for individual lines with perl scripts using the GATK genotype calls and coverage information obtained from pileup files generated using the samtools mpileup function. Ambiguous nucleotide characters were used to identify the two alleles at heterozygous sites. Sites were replaced with an ‘N’ if the coverage was less than 5 or greater than the 99^th^ quantile of the genomic coverage distribution for the given line or if the SNP failed to pass one of the GATK filters.

#### Genomic Alignments

Alignments were made based on the dmel6.01 annotation downloaded from Flybase: <u>ftp.flybase.net/genomes/Drosophila_melanogaster/dmel_r6.01_FB2014_04/gff/</u> dmel-all-r6.01.gff.gz (Santos et al. 2015). The *D. yakuba, D. simulans*, and *D. erecta* reference genomes were separately aligned to the *D. melanogaster* genome using nucmer version 3.23 with parameters –r and –q. A custom perl script then combined the nucmer genomic alignment coordinates and individual line sequences to create genomic sequences for each line that were syntenic to the *D. melanogaster* reference genome. A perl script then called a consensus sequence for each species using these *D. melanogaster* syntenic genome sequences.

### Between species genetic distance

The number of synonymous substitutions in coding genes (K_s_) was used as a measure of genetic distance between species pairs in the *melanogaster* species subgroup. A perl script generated a CDS alignment for each gene using the consensus *D. melanogaster* syntenic genomic sequences for *D. simulans, D. sechellia, D. mauritiana, D. yakuba, D. santomea, D. teissieri*, and *D. orena;* the reference genome sequences for *D. melanogaster;* and the *melanogaster* syntenic genome for *D. erecta*. For genes with multiple annotated transcripts in dmel6.01, we used the longest transcript. We excluded codons that had an N in any of the aligned species. We also excluded genes with either a premature stop codon in any species or whose length was less than 100 bases. We ran PAML version 4.8 (Yang 1997; Yang 2007) to calculate Ks individually for 8,923 genes using the basic model (model=0). PAML was also run with additional models: free ratios (model=1), 3 ratios (model=2, tree = ((*mel, (sim, sech, mau*)^2^)^1^, (((*yak, san), tei), (ore, ere))*^3^)), and 2 ratios (model=2, tree = ((*mel, (sim, sech, mau*))^1^, (((*yak, san), tei), (ore, ere))*^2^)). The super-indices indicate the branches that were allowed to vary in their K_A_/ K_S_. Pairwise K_s_ divergences were obtained by taking the average over all genes.

This genome wide dataset was also used to test whether genes involved in a particular RIM showed evidence for positive selection. We selected genes annotated for eleven GO terms that were related to the nature of each RIM included in this study and calculated their average K_A_/ K_S_. The list of relevant GO terms is shown in Table S11. We assumed a constant K_A_/ K_S_ across the tree (model=0, described immediately above). We could not compare the mean K_A_/ K_S_ value for each GO term with that of the rest of the genome because the sample sizes for each GO term were rather small (mean=7.5 genes per GO term), and there was extensive overlap across GO terms. In lieu of the comparisons, we present all the raw data for each GO term in Table S12.

### Within species neutral variation

We also calculated the level of genetic variation within species as a proxy of genetic distance between individuals of the same species. πs was used as a measure of the average genetic distance between individuals of the same species. We calculated πs and the number of synonymous sites in each gene for each species using Polymorphorama (Andolfatto 2007; Haddrill et al. 2008). A perl script generated a CDS alignment for each species for each gene using the *D. melanogaster* syntenic genome sequences and the dmel 6.01 gene annotations. Since the nexus *D. melanogaster* sequences were mapped to dmel5, we used the dmel5.10 gene annotations for that species. As was the case for interspecific alignments, we only used the longest transcript for genes with multiple transcripts. We only used sequences that were less than 5% Ns and required that at least 5 individuals met this criterion. We also excluded genes if a premature stop codon was encountered in any individual. A measure of within species variation for each species was obtained by averaging πs over all genes weighted by the number of synonymous sites.

### Reproductive isolation vs. genetic distance

Finally, we used a logistic regression with complete taxon sampling to analyze whether the distance between potentially hybridizing species influence the magnitude of reproductive isolation. Each index of reproductive isolation was independently regressed against the divergence between the parental species using the glm function with a logit link function with binomial errors in R ('stats' package, R Core Team 2016). K_s_ was used as a measure of neutral species divergence, and π_s_ was used as a proxy of neutral within species variation. Since we did not have population data for both *D. erecta* and *D. orena*, we used the average πs for the 7 other species: 0.0208.

The logistic function is given by:

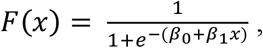

where the parameter β_1_ determines how quickly the function approaches 1 with larger values producing steeper slopes. The fit of each function was determined with a McFadden’s pseudo-r^2^ calculated with the R package ‘pscl’ (function ‘pR2’, Jackman et al. 2007).

To compare the rates of evolution of the four types of RI, we calculated the level of genetic divergence where the magnitude of RI crossed a 0.95 threshold. For the *melanogaster* subgroup analysis, genetic divergence was measured as Ks (Threshold_Ks). For the analyses in the *Drosophila* genus, genetic distance was measured as Nei’s D (Threshold_D; See below ‘Generality of the pattern: Other clades’). To compare the relative evolutionary rates of premating, NCGI, CSP, and hybrid inviability, we bootstrapped all four datasets 10,000 times and compared the bootstrap distributions of the Ks (or Nei’s D, depending on the analysis) where the threshold was crossed using a Mann-Whitney U test ('stats' package, R Core Team 2016). The same statistical methods were applied for regressions comparing evolutionary rates of embryo, larval, and pupal stage viability.

### Detection of reinforcement using comparative analyses

The magnitude on RI can be affected by the influence of reinforcement, a type of natural selection that strengthens prezygotic isolation as a byproduct to reduce maladaptive hybridization (Servedio and Noor 2003, but see Coyne 1974 for an argument of reinforcement of postzygotic isolation). To detect whether reinforcement has played a widespread role on the evolution of any particular of RI, we followed two approaches. First, we compared the magnitude of all the above mentioned types of RI in eight pairs of species for which we had sympatric and allopatric species (N = 8 species, 1 sympatric line pair and 1 allopatric line pair per species pair). In both sympatric and allopatric crosses, we measured reproductive isolation as described above. We compared the magnitude of each RIM (three types of prezygotic isolation, hybrid inviability as a whole, and the three developmental components of inviability) using two methods. First we quantified the amount of genetic divergence required to achieve 95% of the maximum value of RI in sympatric and allopatric populations independently. These values were then compared by generating 1,000 bootstrap replicates and a two-tailed Wilcoxon sign test (as described above). The expectation of this comparison is that if a RIM is evolving through reinforcement, sympatric lines should show a stronger RI than allopatric lines from the same species for that RIM. Second, we assessed whether the effect of geographic overlap (i.e., whether lines were sympatric or allopatric) influenced the magnitude of RI (while controlling by cross) by fitting linear models where each type of RI (7 linear models excluding female and male sterility) was the response and depended on the identity of the cross, the geographic origin (whether a line was sympatric or allopatric), and the interaction between these two main effects.

The second approach aimed to detect the phylogenetic signature of reinforcement (Noor 1997). We compared the magnitude of all RIMs in two species triads: (*D. teissieri*, (*D. yakuba, D. santomea*)) and (*D. melanogaster*, (*D. sechellia, D. simulans*)). Notably these triads include a pair of species that is sympatric (*D. teissieri, D. yakuba;* and *D. melanogaster, D. simulans*) and one that is allopatric (*D. teissieri, D. santomea;* and *D. melanogaster, D. sechellia*). If reinforcing selection has acted, then the magnitude of RI should be stronger in the sympatric pairs than in the allopatric pairs. We pooled the two directions of the cross for each pair and compared the mean strength of each RIM using permutation tests (function ‘oneway_test’ with and 9,999 Monte Carlo iterations; R package ‘coin’).

### Robustness of calculations of the rate of evolution of RI

Our measurements of the rate of evolution of RI on the *melanogaster* subgroup have an important caveat: since all the species are closely related and our design involved measuring all possible pairwise interactions, we could not apply phylogenetic corrections. This is an important limitation because if one species is more likely than others to be reproductively isolated and the branch leading to that species is used more than once, it might inflate the rate of evolution of a particular RI. Several approaches have been proposed to correct non-independent measurements of RI (i.e., those that include a species or a branch more than once). Nonetheless, reconstructing levels of ancestral RI at a node might be problematic as reproductive isolation does not follow the regular assumptions of quantitative traits (e.g., Moyle et al. 2004). We opted for a more conservative approach in which we performed regression using only strictly independent species pairs (i.e., non-overlapping branches; Figure S6). We thus evaluated whether the relative ranking of the rates of evolution of the four types of RIMs obtained in the *melanogaster* comparisons also held when we did a similar analysis with phylogenetically independent points. We evaluated our hypothesis in a phylogenetically independent subset of species from the *melanogaster* subgroup. In this case, three species pairs is the maximum number of independent species pairs (Figure S6). We also included measurements for hybridizations of the *willinstoni (D. paulistorum* Centroamerica, *D. paulistorum* Interior), *pseudoobscura (D. pseudoobscura, D. persimilis, D. bogotana), virilis (D. virilis, D. lummei, D. americana, D. novamexicana)*, and *mojavensis (movavensis baja, mojavensis* sonora) group. Nei’s D distance between these species was obtained from Yukilevich (2012). The choice of groups and species was dictated by the existence of phenotypic mutants and limited by the ability to measure sperm precedence in conspecific crosses; we needed mutant stocks to be able to quantify CC_C1_ and CC_C2_, two required components of the I_CSP_ indexes (see above). To this end, only four species satisfied the requirement: *D. paulistorum* Centroamerica *white* (14030–0771.04), *D. virilis eGFP* (15010–1051.108), *D. pseudoobscura GFP* (10411–0121.201), and *D. mojavensis w* (15081–1352.05). The performed crosses and sample sizes are shown in Table S16. We next estimated the rate of evolution of premating, non-competitive gametic isolation, competitive gametic isolation, and hybrid inviability in crosses for each group as described above.

## ACKNOWLEDGEMENTS

We would like to thank A.A. Comeault, V. Courtier-Orgogozo, Y. Brandvain, R. Marquez, K. L. Gordon and the members of the Matute lab for helpful scientific discussions and comments. We would also like to thank the Bioko Biodiversity Protection Program, and the Ministry of Environment, Republic of São Tomé and Príncipe for permission to collect and export specimens for study. The authors have no conflicts of interest.

## SUPPLEMENTARY INFORMATION

### SUPPLEMENTARY FIGURES

**FIGURE S1.**
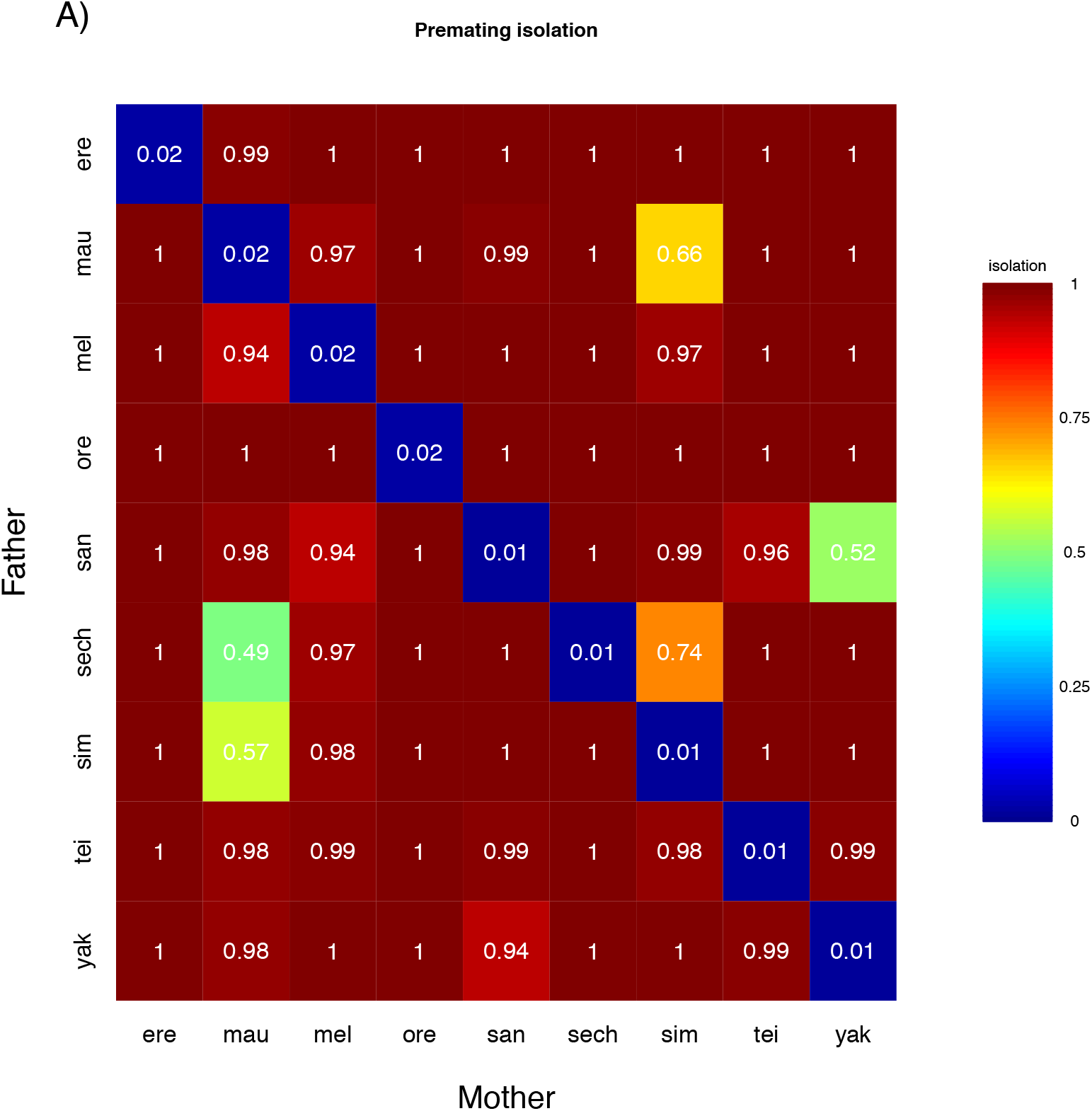
Levels of premating isolation in the *melanogaster* species subgroup. Average prezygotic isolation per cross.

**FIGURE S2.**
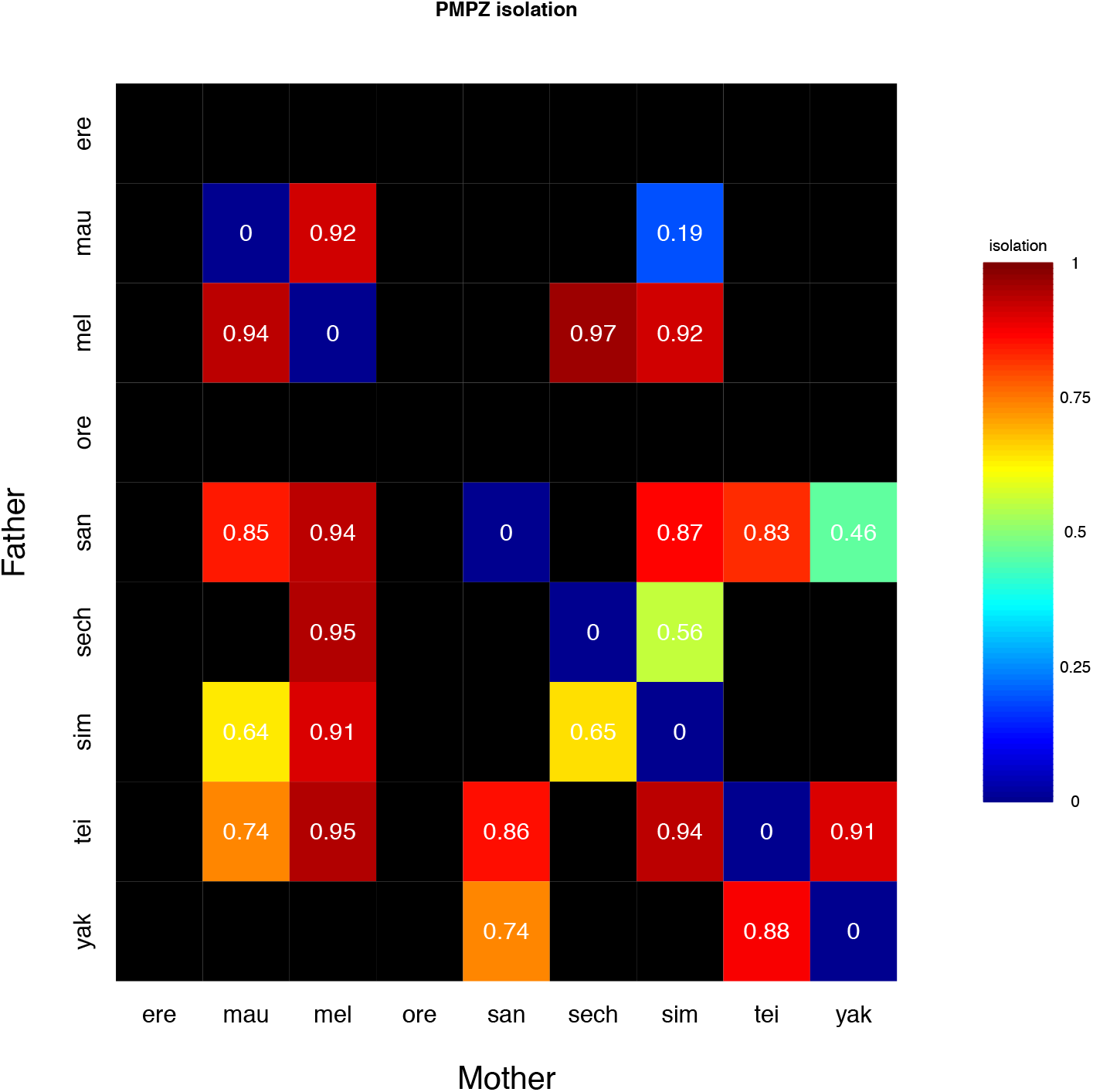
Levels of NCGI in the *melanogaster* species subgroup. Average PMPZ isolation per cross. Black rectangles represent crosses that did not produce embryos (either dead or alive).

**FIGURE S3.**
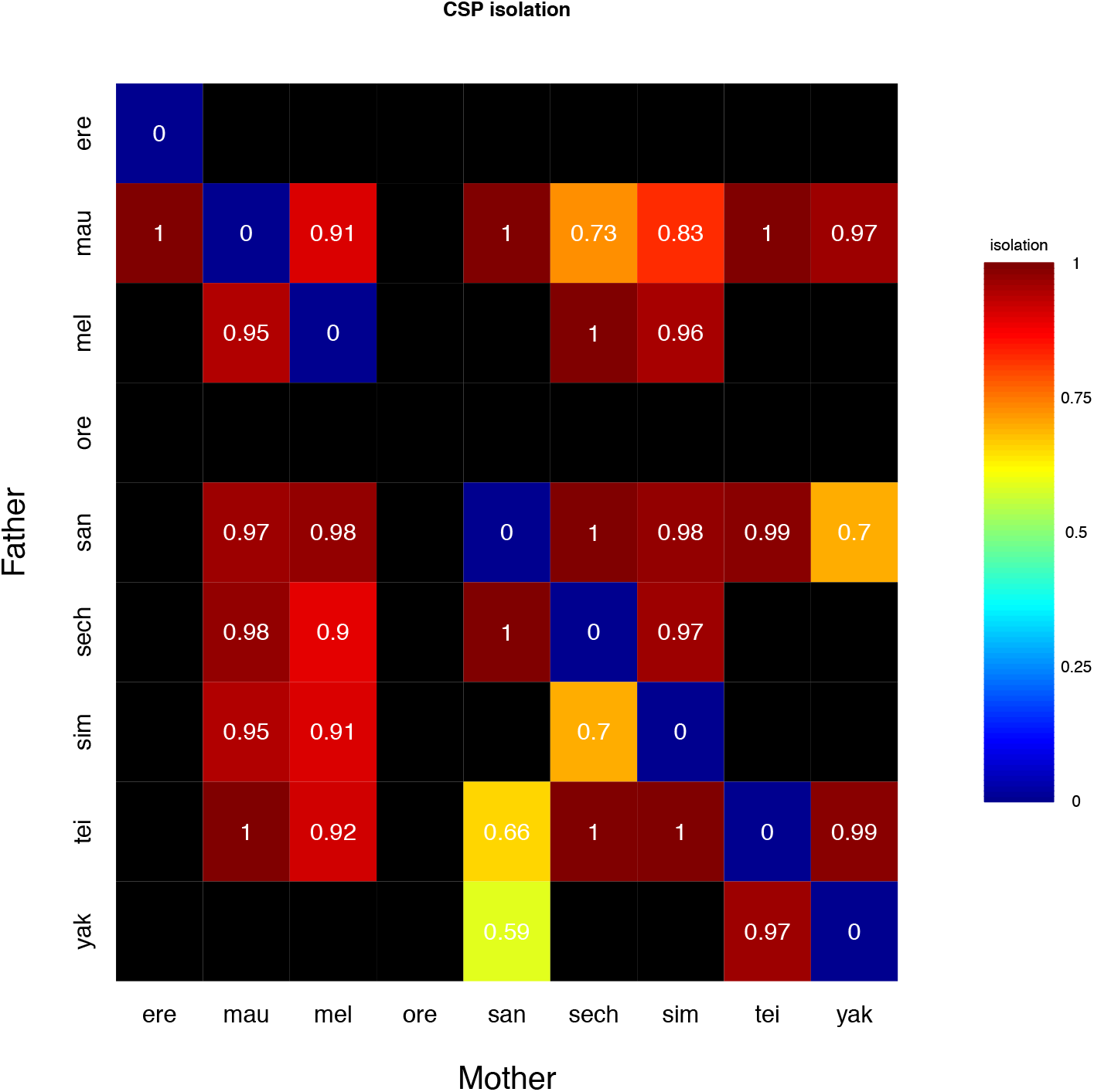
Levels of conspecific sperm precedence in the *melanogaster* species subgroup. Average conspecific sperm precedence isolation per cross. Black rectangles represent crosses that did not produce embryos (either dead or alive).

**FIGURE S4.**
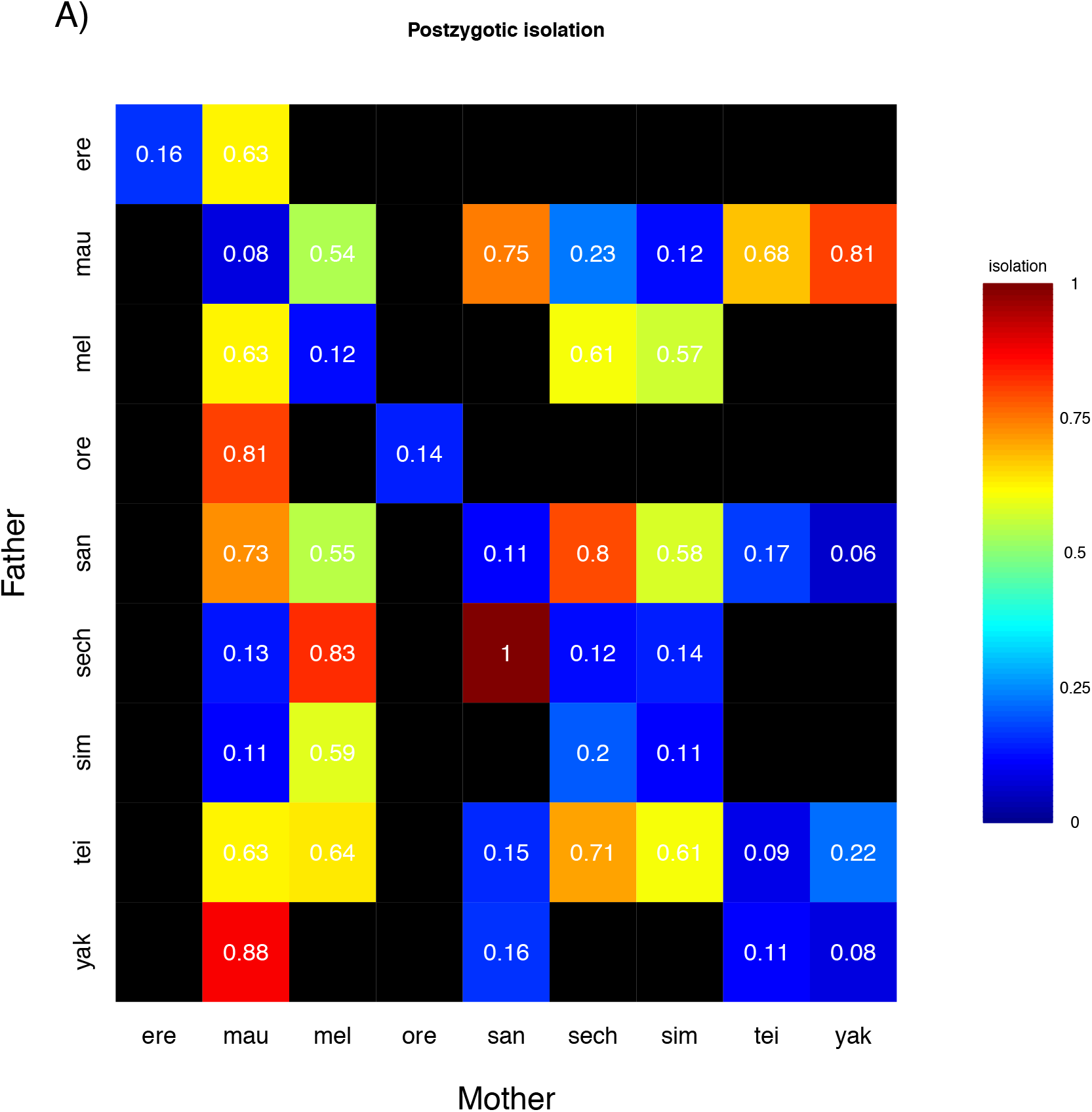
Levels of postzygotic isolation in the *melanogaster* species subgroup. Black rectangles represent crosses that did not produce viable eggs. (A) Average postzygotic isolation per cross.

**FIGURE S5.**
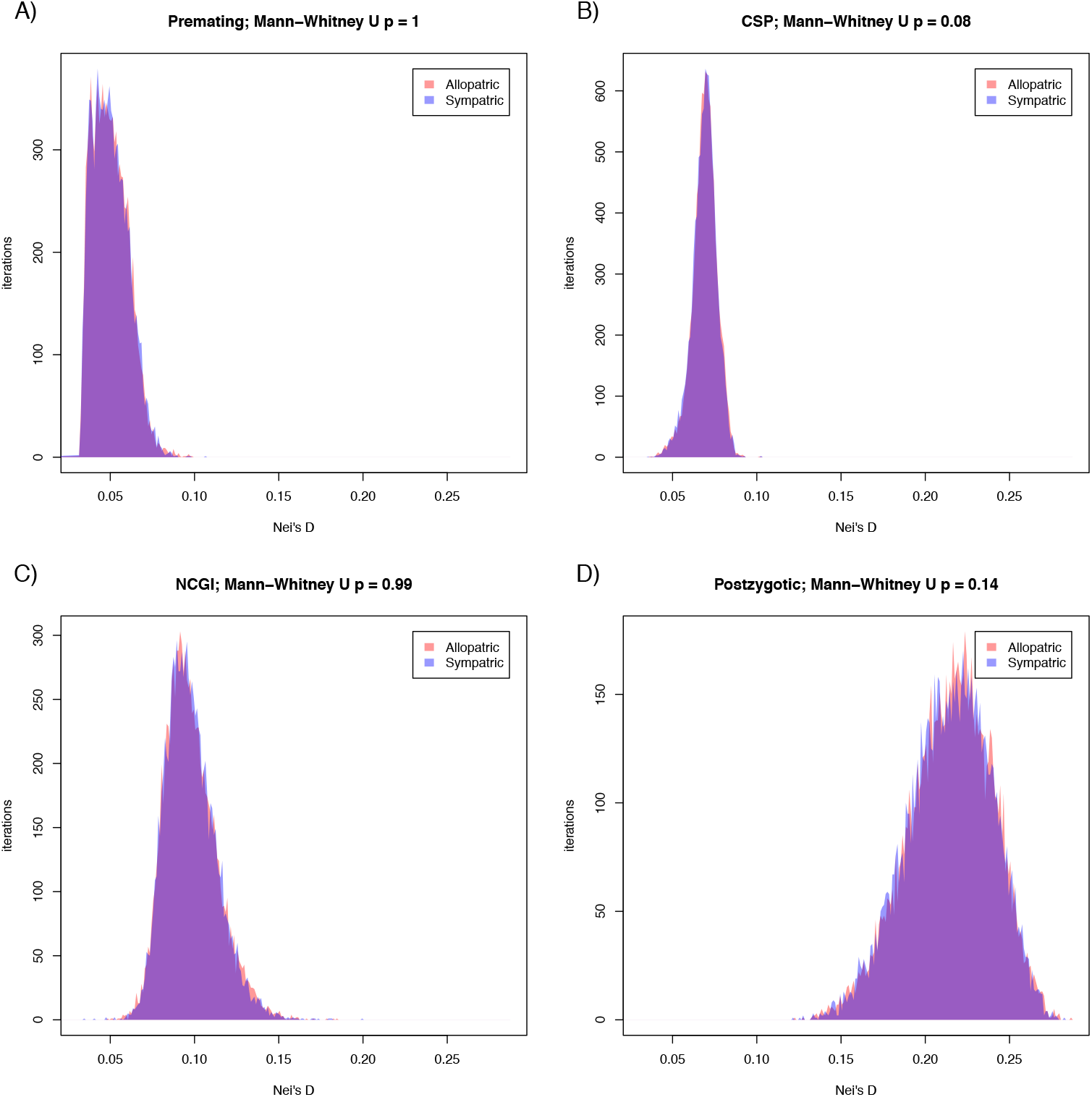
No evidence for generalized reinforcement at premating, conspecific sperm precedence, noncompetitive gametic isolation, or postzygotic isolation in the *melanogaster* species subgroup. **To detect the signature of reinforcing selection, we compared the** magnitude of the four RIMs between sympatric and allopatric lines. Comparisons were done using a Mann-Whitney U test on bootstrapped values of Threshold_Ks as described in the main text (e.g., Figures 2, 4, and 6).

**FIGURE S6.**
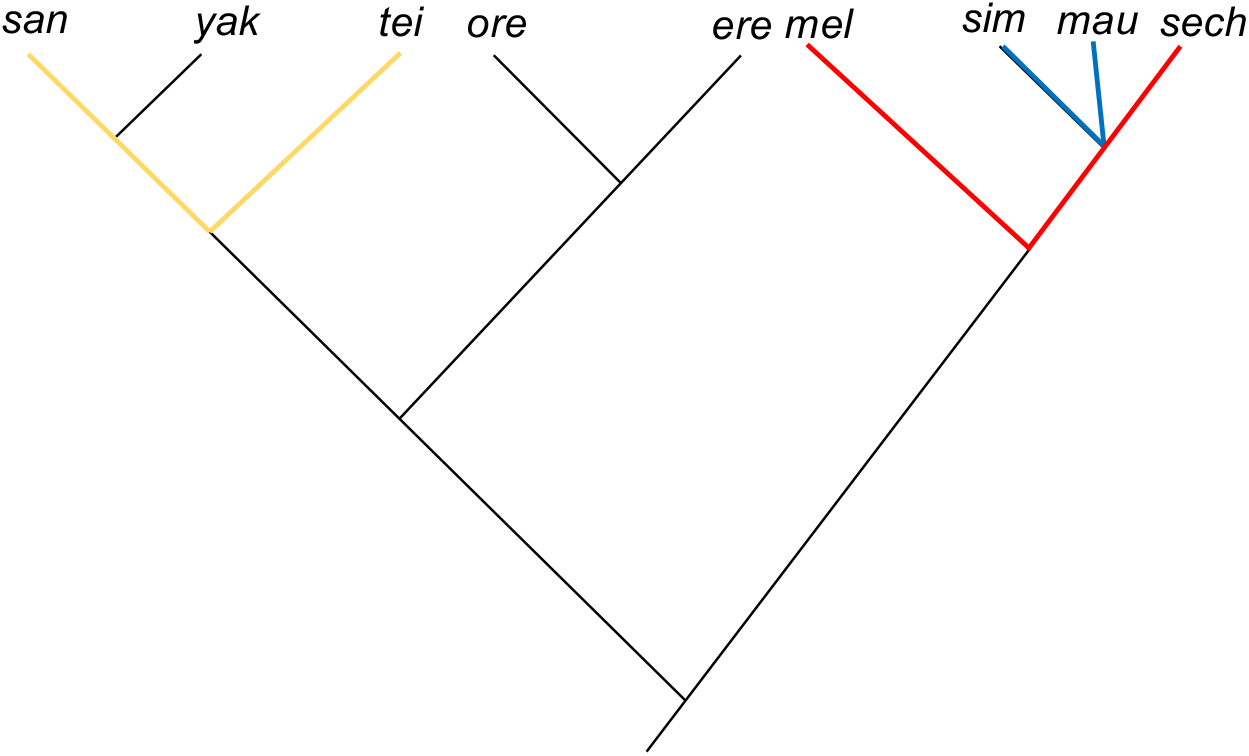
Phylogenetic tree depicting our approach to perform phylogenetic corrections. The tree shows the *melanogaster* species group and the three possible non-overlapping species pairs that are phylogenetically independent. Blue: *D. simulans-D. mauritiana*; Red: *D. melanogaster-D. sechellia*; Yellow: *D. santomea-D. teissieri*. The other four species pairs belong to different species subgroups and they are phylogenetically independent.

**FIGURE S7.**
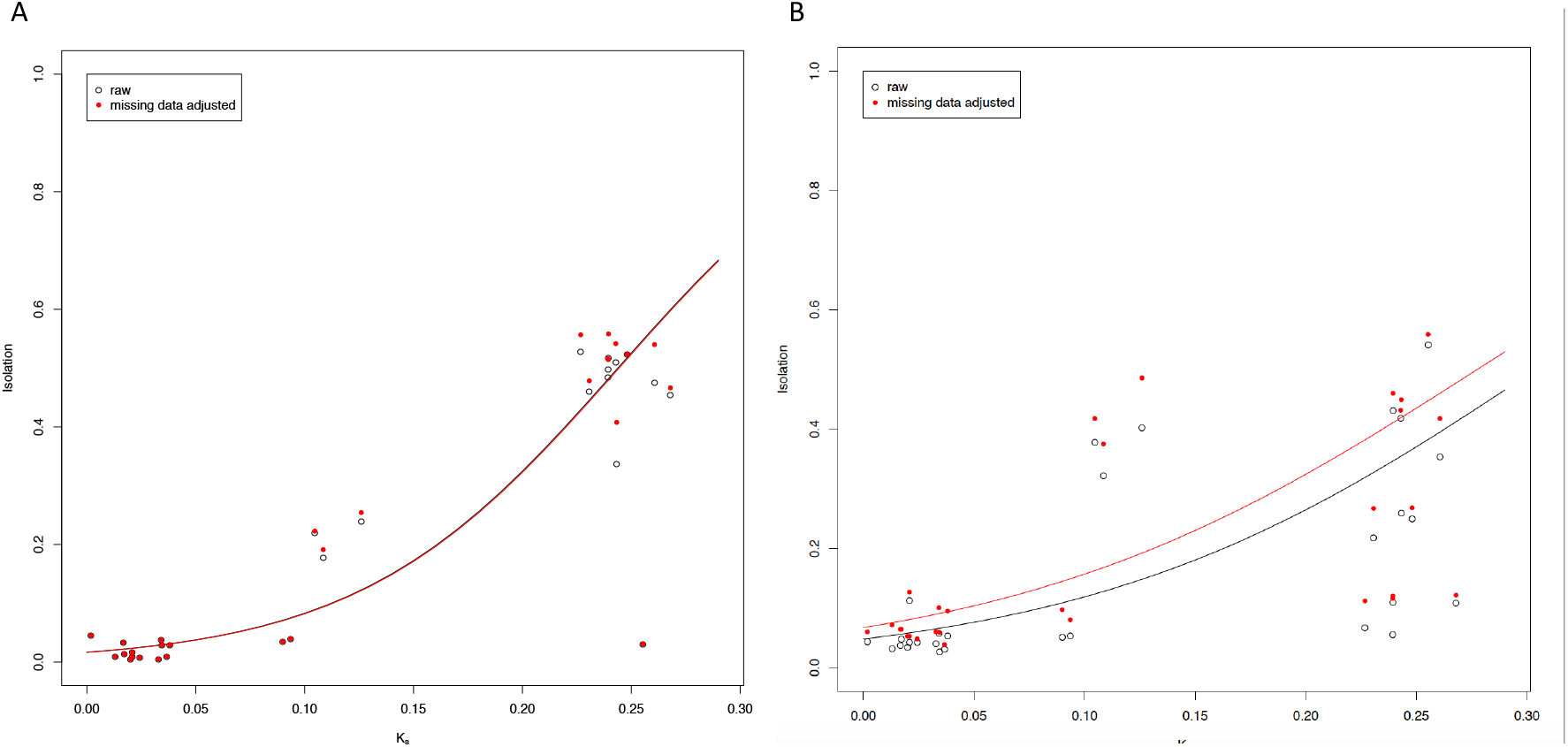
Measurements of egg and larval viability are robust to missing data. Feeding larvae can bury egg cases and dead embryos resulting in inaccurate counts. We adjusted counts of egg cases and dead embryos to account for missing data, and our results were unaffected. Estimates of isolation based on raw data are shown as black circles and adjusted estimates appear as red dots. The black line represents the logistic fit to the raw data, and the red line is the logistic fit to the adjusted data. Phylogenetic distance was measured as K_s_ between species and π_s_ within species. A. Postzygotic embryonic lethality estimations. B. Postzygotic larval lethality estimations.

### SUPPLEMENTARY TABLES

**TABLE 1.**
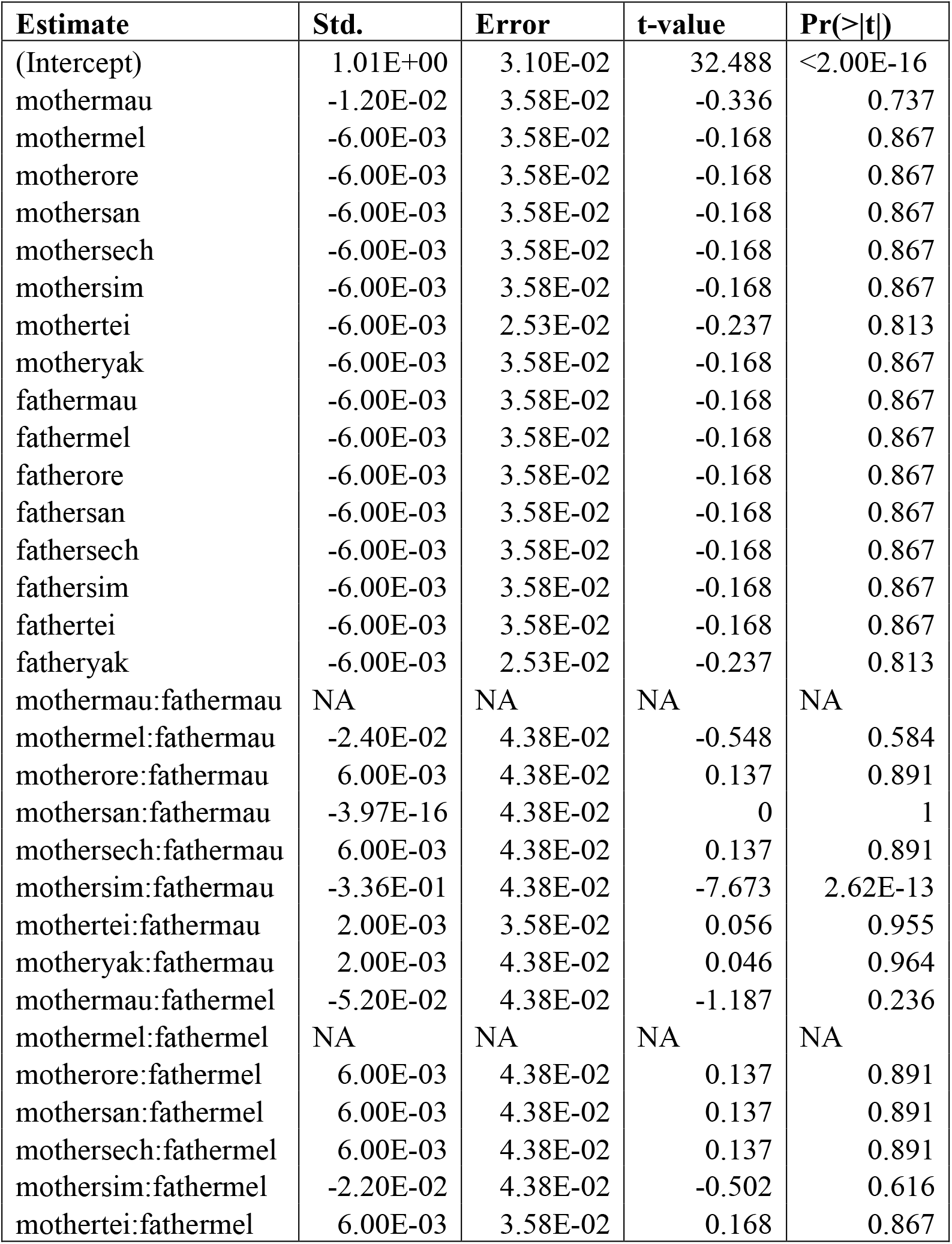
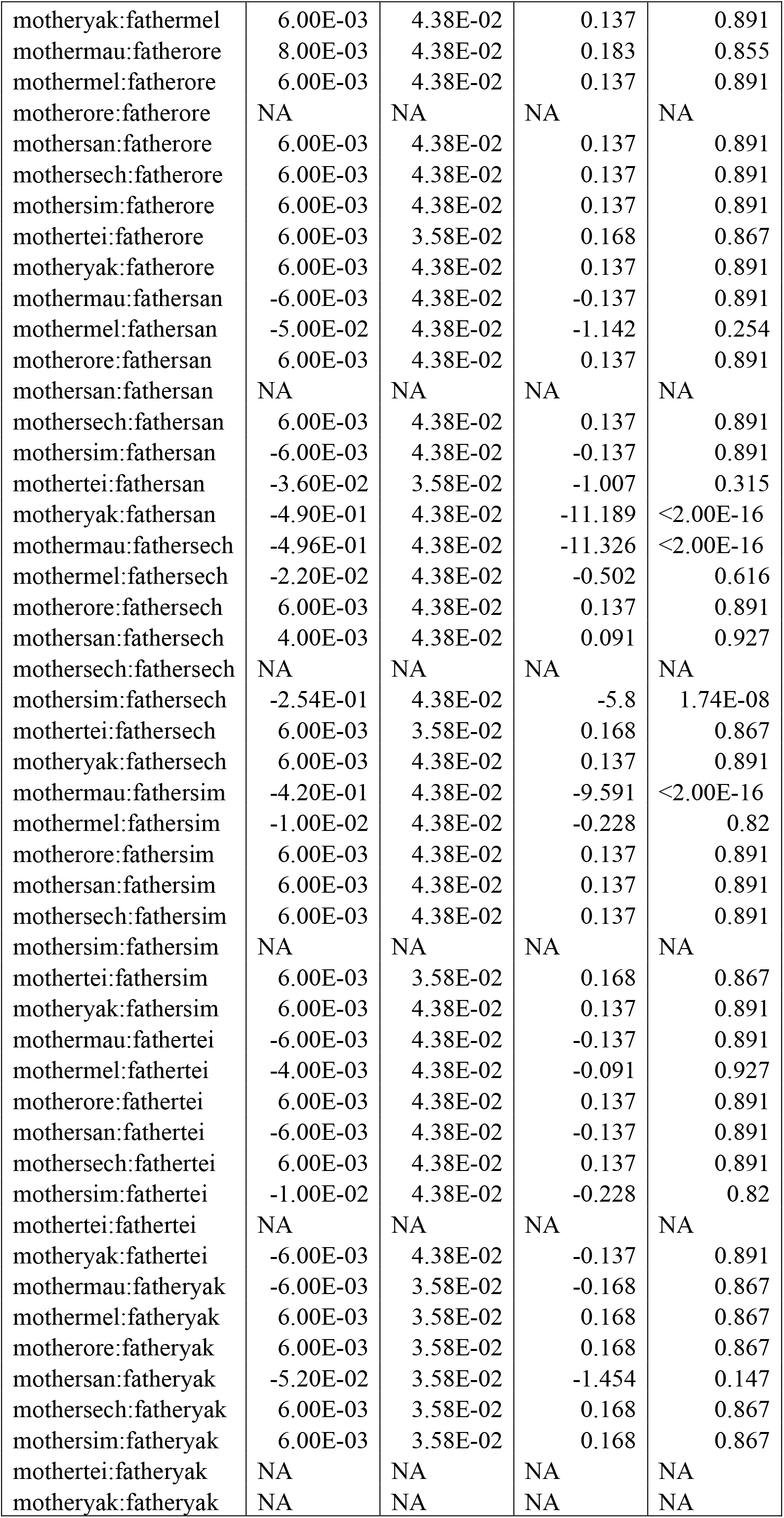
Linear contrasts for a full-factorial model analysis of the magnitude of premating isolation. Each genotype is summarized by the sex followed by the first three letters of the species. All linear contrasts were done using the number of degrees of freedom from the residuals of the linear model. df =288.

**TABLE S2.**
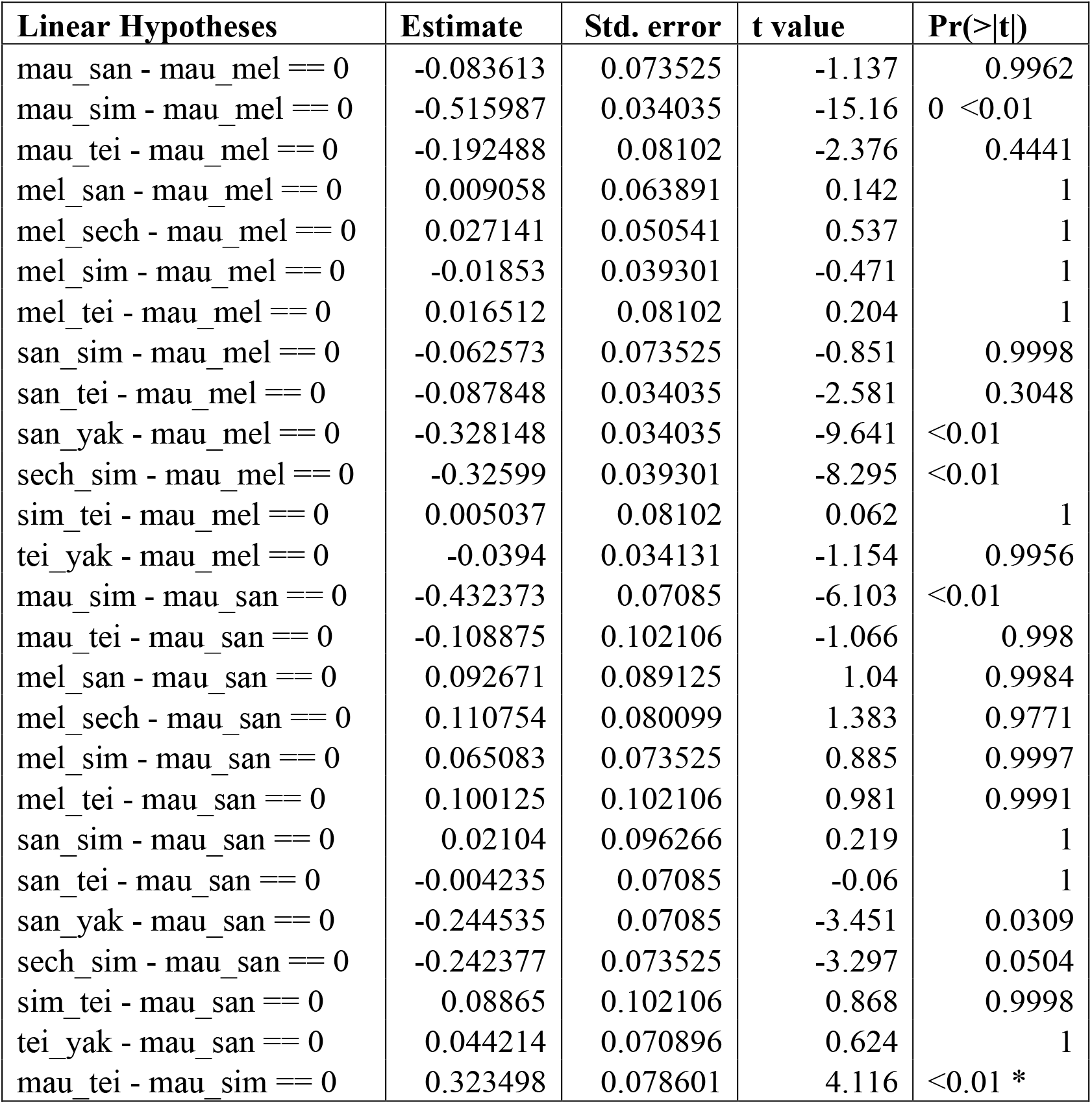
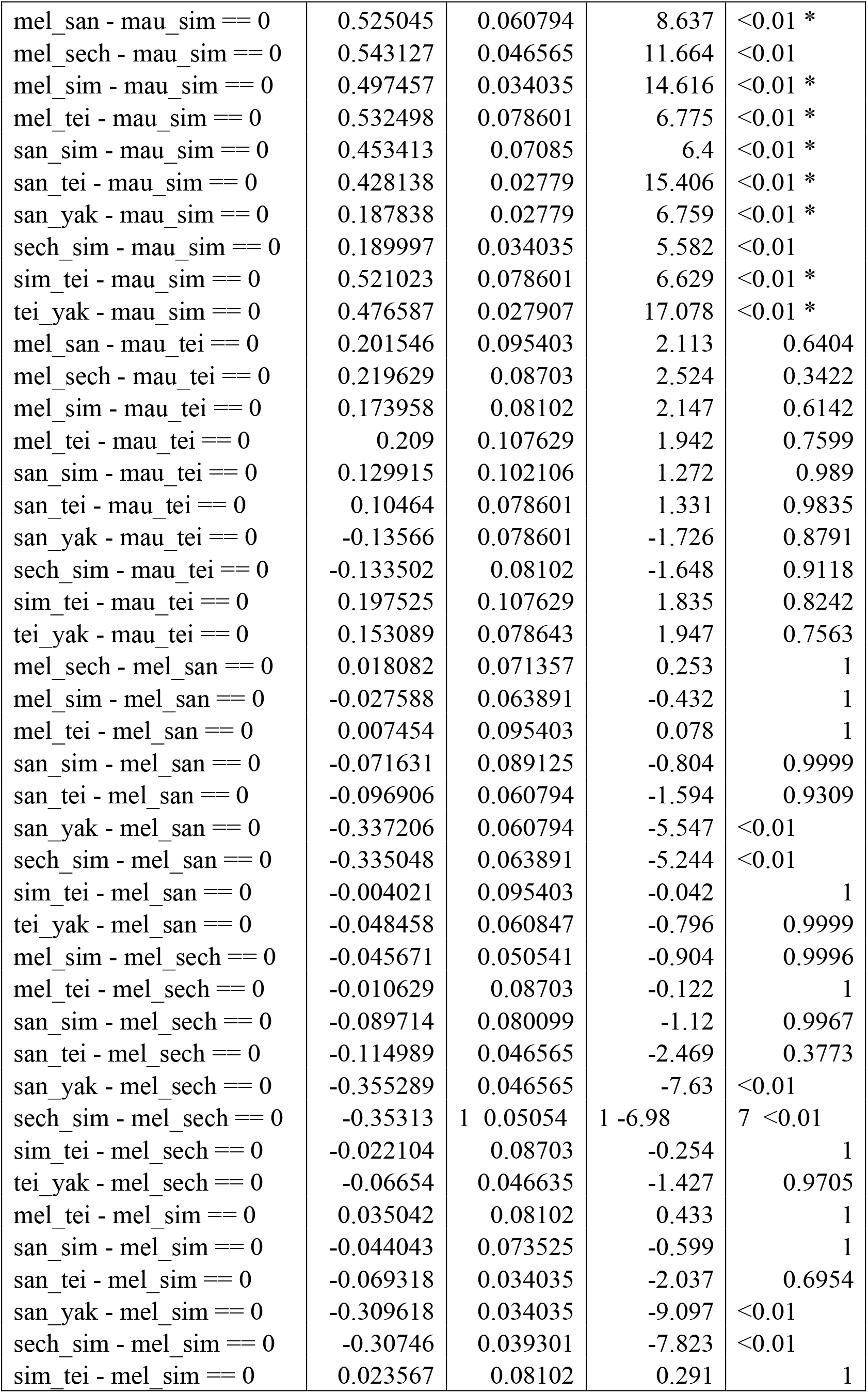
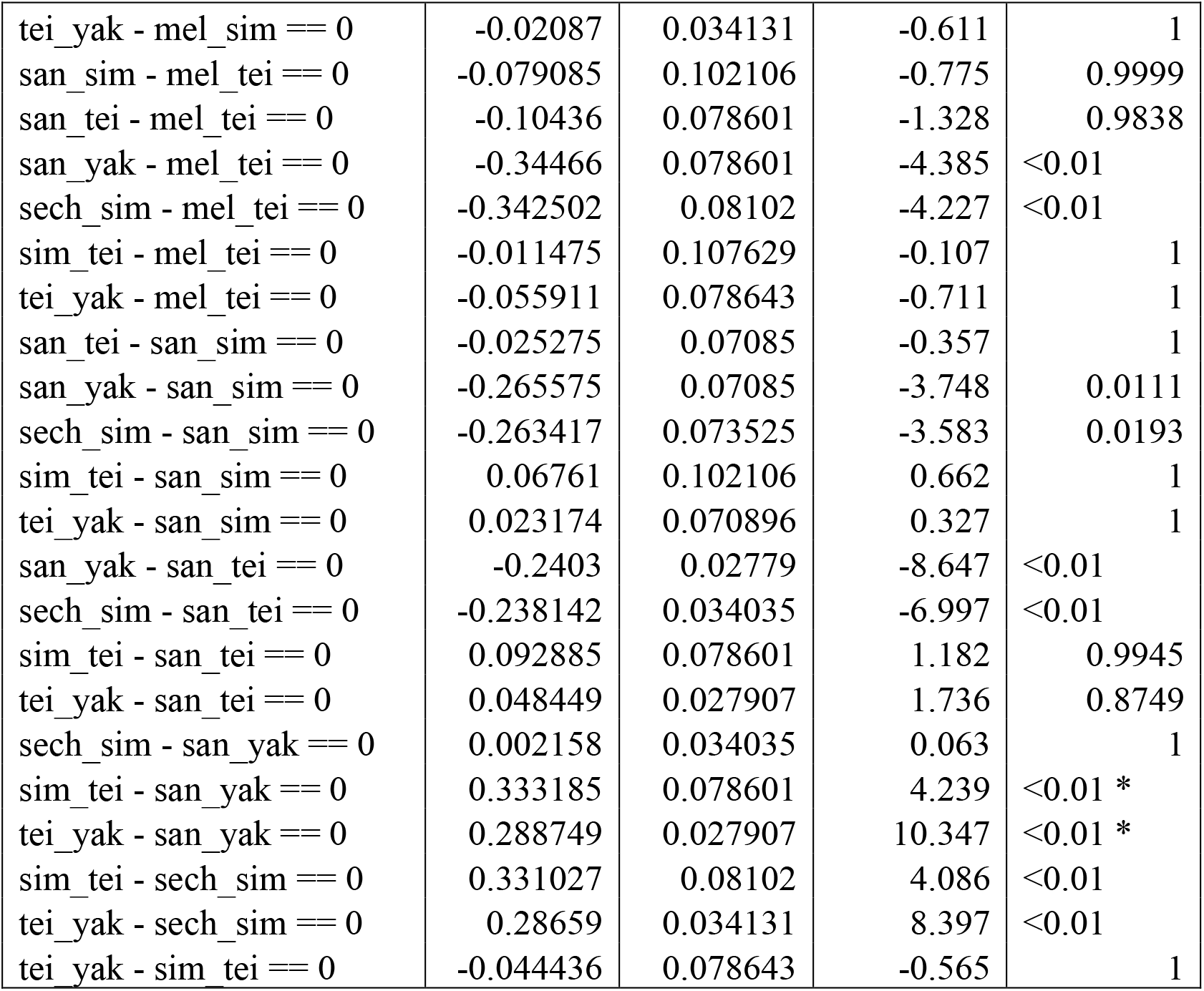
Pairwise comparisons between the magnitude of NCGI in different crosses between species of the *melanogaster* subgroup. Each hybridization is summarized by the first three letters of the genotype of the female, an underscore, and the first three letters of the genotype of the male. The number of degrees of freedom for each pairwise comparison equaled the total number of observations minus the number of means; df =137.

**TABLE S3.** Sample sizes of conspecific sperm precedence experiments in the *melanogaster* species subgroup.

| **Female** | **male_1** | **male_2** | **reps** |
| --- | --- | --- | --- |
| *D. erecta* | *D. erecta* | *D. erecta* | 26 |
| *D. erecta* | *D. mauritiana* | *D. erecta* | 2 |
| *D. mauritiana* | *D. mauritiana* | *D. mauritiana* | 14 |
| *D. mauritiana* | *D. mauritiana* | *D. melanogaster* | 3 |
| *D. mauritiana* | *D. melanogaster* | *D. mauritiana* | 4 |
| *D. melanogaster* | *D. mauritiana* | *D. melanogaster* | 11 |
| *D. mauritiana* | *D. santomea* | *D. mauritiana* | 3 |
| *D. santomea* | *D. mauritiana* | *D. santomea* | 1 |
| *D. mauritiana* | *D. mauritiana* | *D. sechellia* | 3 |
| *D. mauritiana* | *D. sechellia* | *D. mauritiana* | 10 |
| *D. sechellia* | *D. mauritiana* | *D. sechellia* | 9 |
| *D. mauritiana* | *D. mauritiana* | *D. simulans* | 4 |
| *D. mauritiana* | *D. simulans* | *D. mauritiana* | 18 |
| *D. simulans* | *D. mauritiana* | *D. simulans* | 15 |
| *D. simulans* | *D. simulans* | *D. mauritiana* | 6 |
| *D. mauritiana* | *D. teissieri* | *D. mauritiana* | 2 |
| *D. teissieri* | *D. mauritiana* | *D. teissieri* | 4 |
| *D. teissieri* | *D. teissieri* | *D. mauritiana* | 1 |
| *D. yakuba* | *D. mauritiana* | *D. yakuba* | 6 |
| *D. yakuba* | *D. yakuba* | *D. mauritiana* | 1 |
| *D. melanogaster* | *D. melanogaster* | *D. melanogaster* | 32 |
| *D. melanogaster* | *D. santomea* | *D. melanogaster* | 7 |
| *D. melanogaster* | *D. sechellia* | *D. melanogaster* | 9 |
| *D. sechellia* | *D. melanogaster* | *D. sechellia* | 5 |
| *D. melanogaster* | *D. simulans* | *D. melanogaster* | 10 |
| *D. simulans* | *D. melanogaster* | *D. simulans* | 9 |
| *D. simulans* | *D. simulans* | *D. melanogaster* | 2 |
| *D. melanogaster* | *D. teissieri* | *D. melanogaster* | 4 |
| *D. santomea* | *D. santomea* | *D. santomea* | 11 |
| *D. santomea* | *D. sechellia* | *D. santomea* | 1 |
| *D. sechellia* | *D. santomea* | *D. sechellia* | 2 |
| *D. simulans* | *D. santomea* | *D. simulans* | 3 |
| *D. simulans* | *D. simulans* | *D. santomea* | 1 |
| *D. santomea* | *D. santomea* | *D. teissieri* | 2 |
| *D. santomea* | *D. teissieri* | *D. santomea* | 6 |
| *D. teissieri* | *D. santomea* | *D. teissieri* | 6 |
| *D. teissieri* | *D. teissieri* | *D. santomea* | 3 |
| *D. santomea* | *D. santomea* | *D. yakuba* | 3 |
| *D. santomea* | *D. yakuba* | *D. santomea* | 8 |
| *D. yakuba* | *D. santomea* | *D. yakuba* | 16 |
| *D. yakuba* | *D. yakuba* | *D. santomea* | 12 |
| *D. sechellia* | *D. sechellia* | *D. sechellia* | 24 |
| *D. sechellia* | *D. simulans* | *D. sechellia* | 9 |
| *D. simulans* | *D. sechellia* | *D. simulans* | 15 |
| *D. simulans* | *D. simulans* | *D. sechellia* | 2 |
| *D. sechellia* | *D. teissieri* | *D. sechellia* | 1 |
| *D. simulans* | *D. simulans* | *D. simulans* | 16 |
| *D. simulans* | *D. teissieri* | *D. simulans* | 2 |
| *D. teissieri* | *D. teissieri* | *D. teissieri* | 14 |
| *D. teissieri* | *D. teissieri* | *D. yakuba* | 3 |
| *D. teissieri* | *D. yakuba* | *D. teissieri* | 4 |
| *D. yakuba* | *D. teissieri* | *D. yakuba* | 4 |
| *D. yakuba* | *D. yakuba* | *D. teissieri* | 2 |
| *D. yakuba* | *D. yakuba* | *D. yakuba* | 13 |

**TABLE S4.**
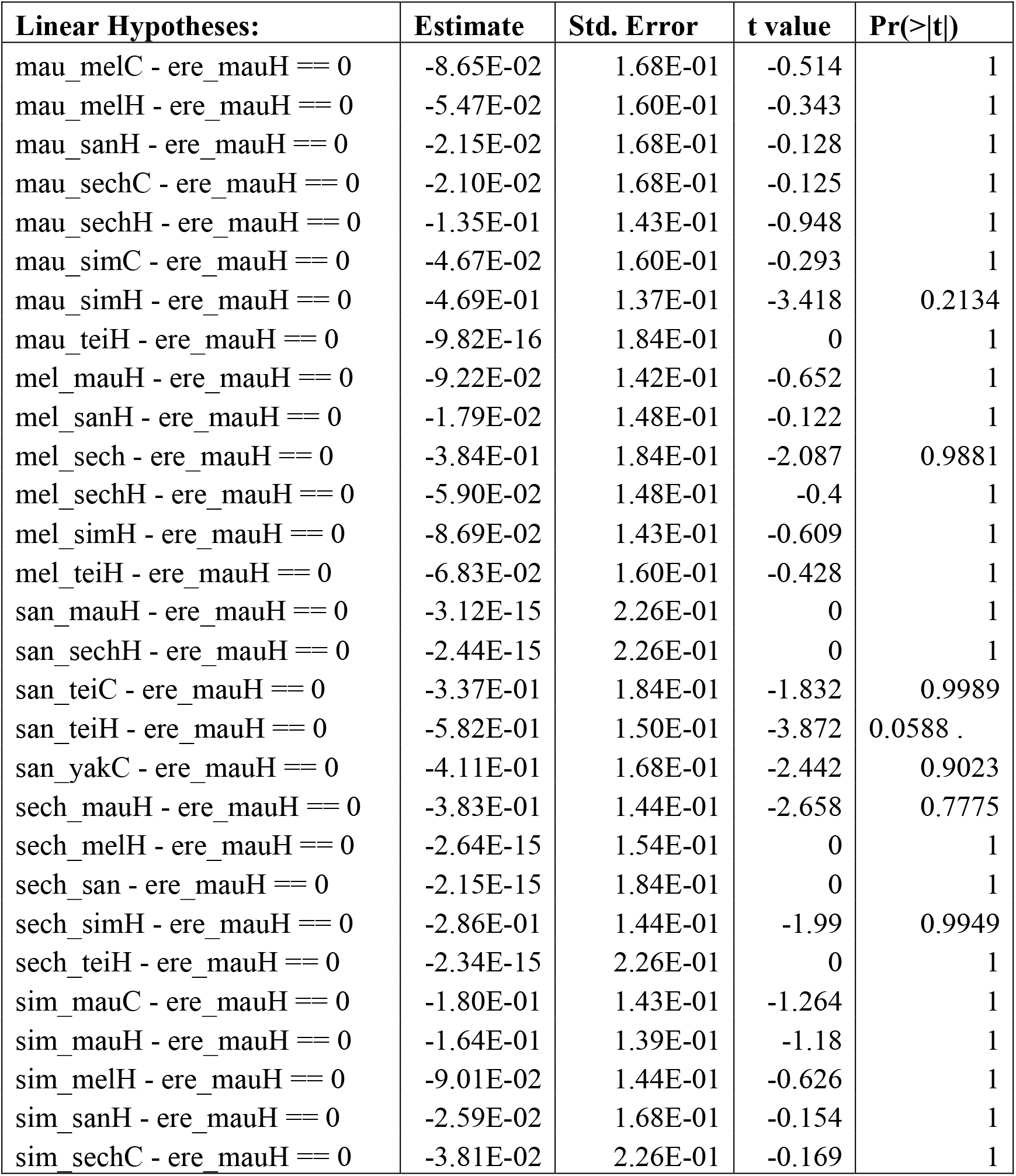
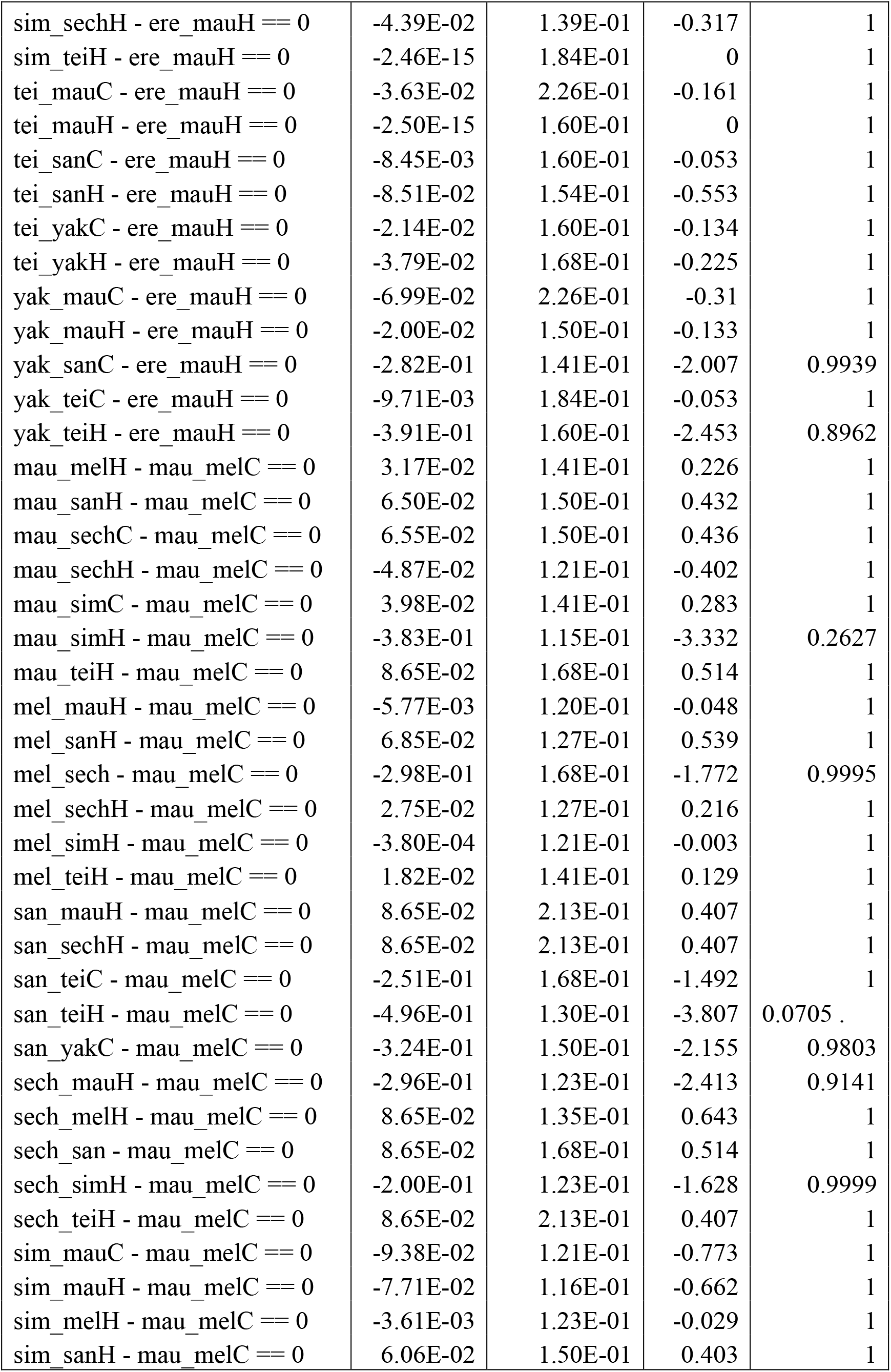
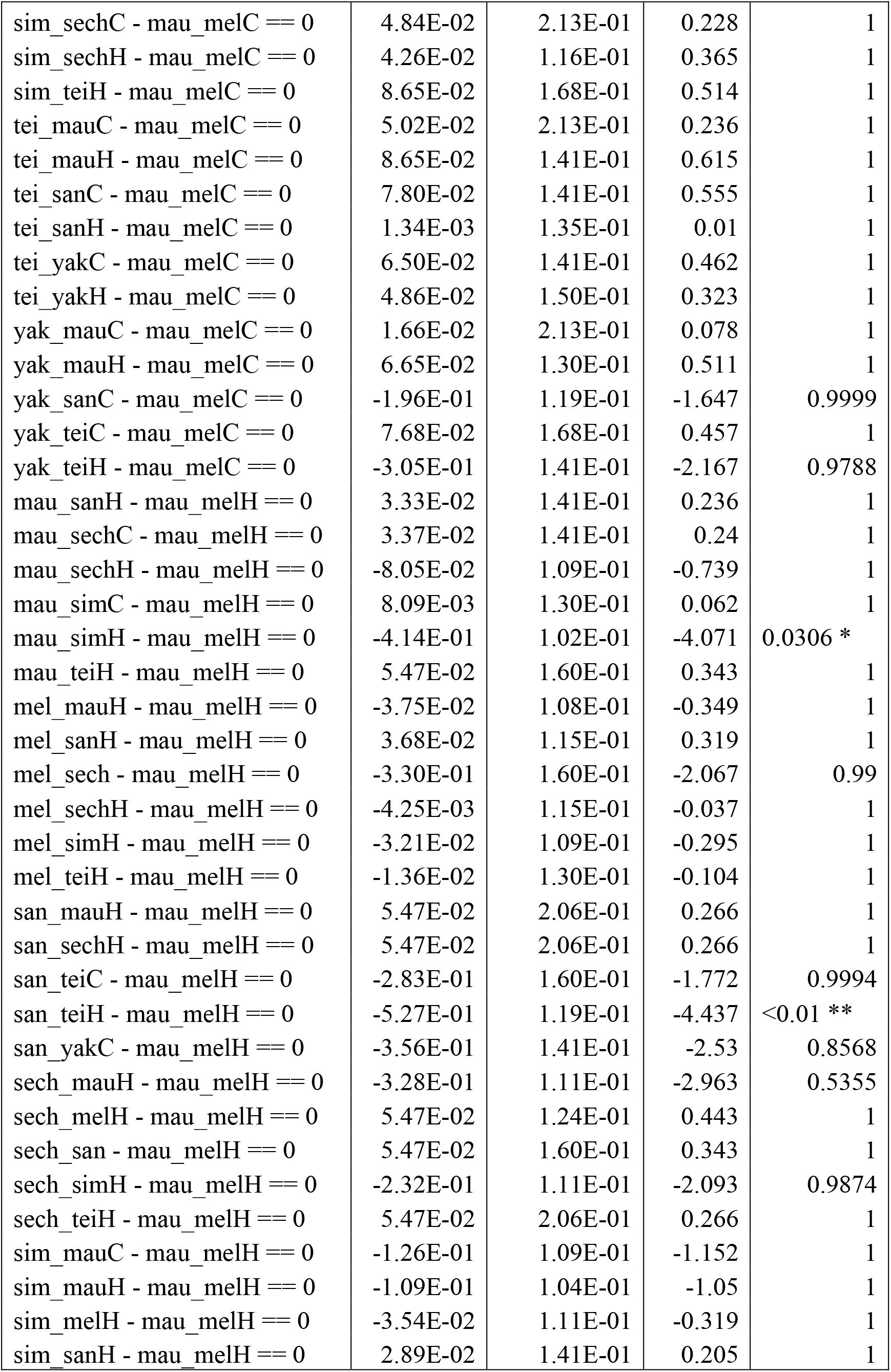
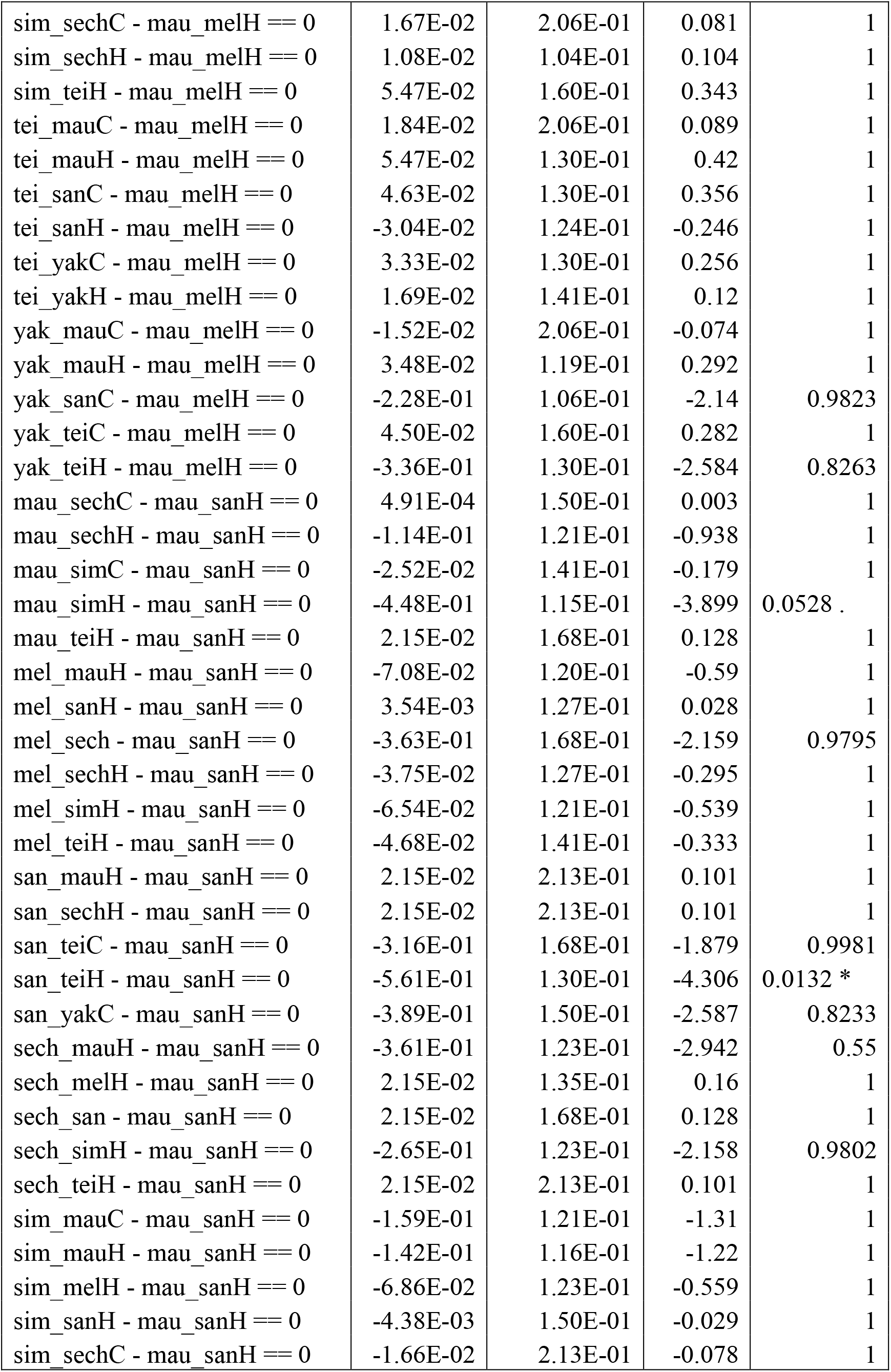
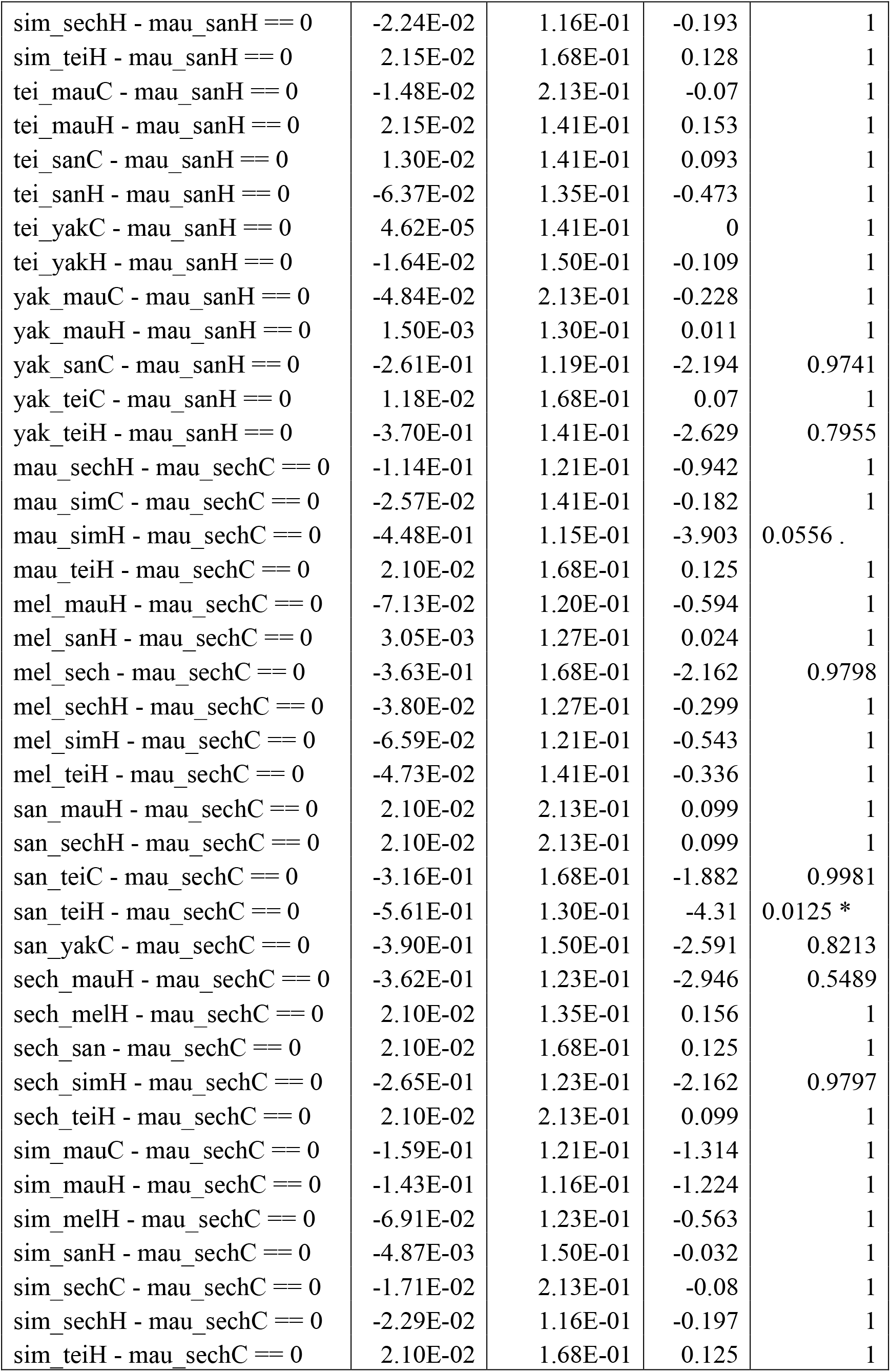
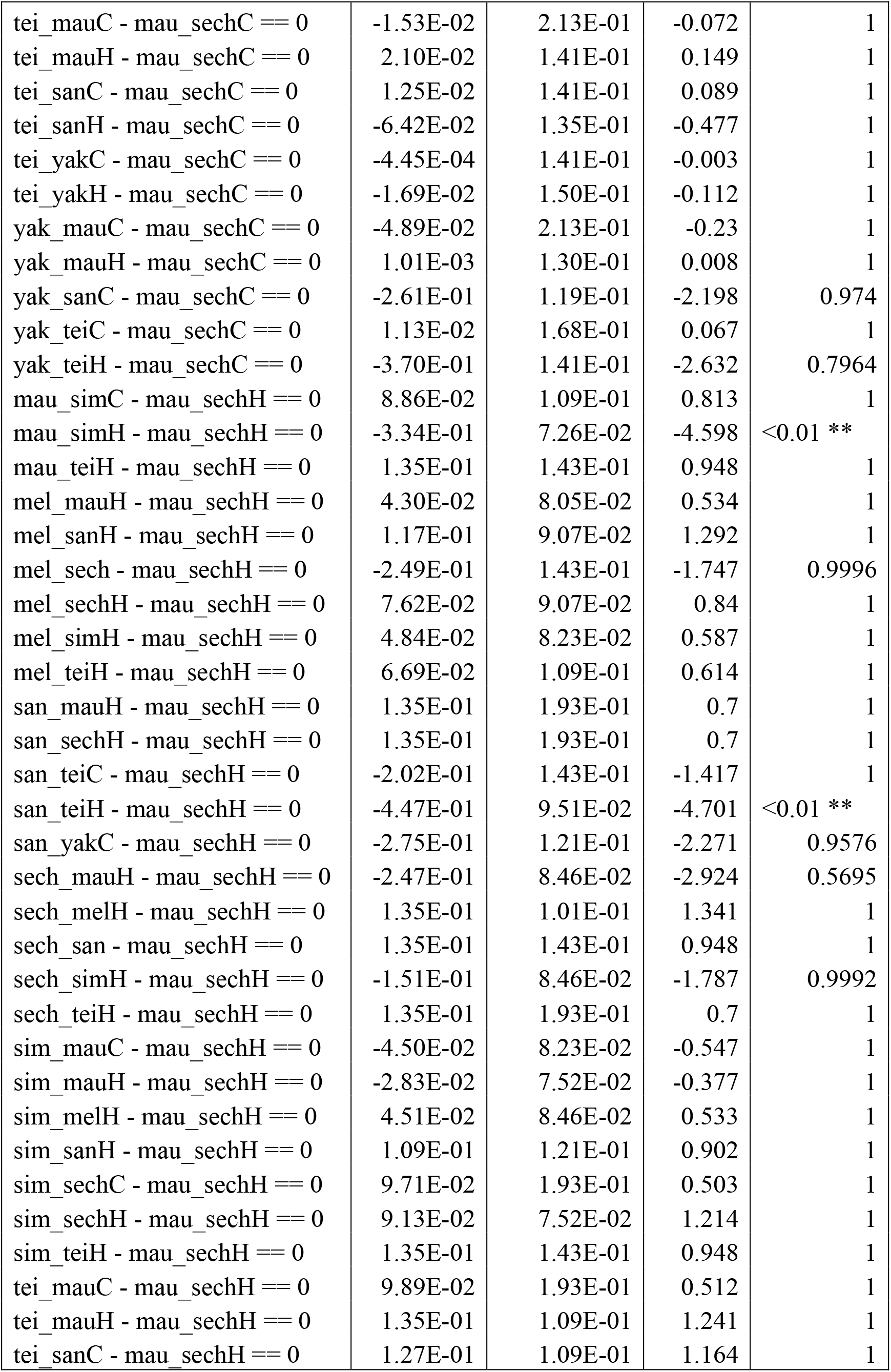
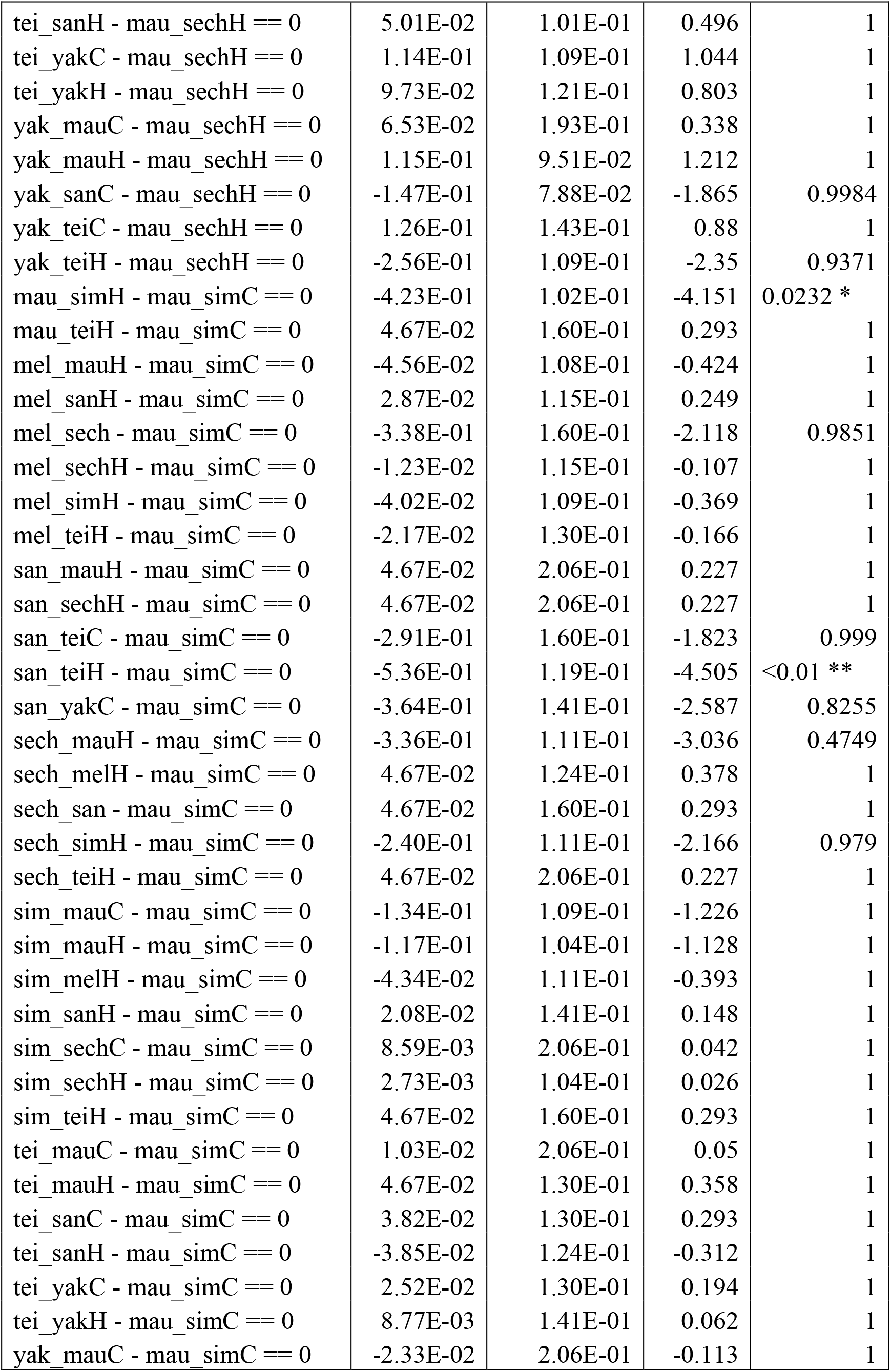
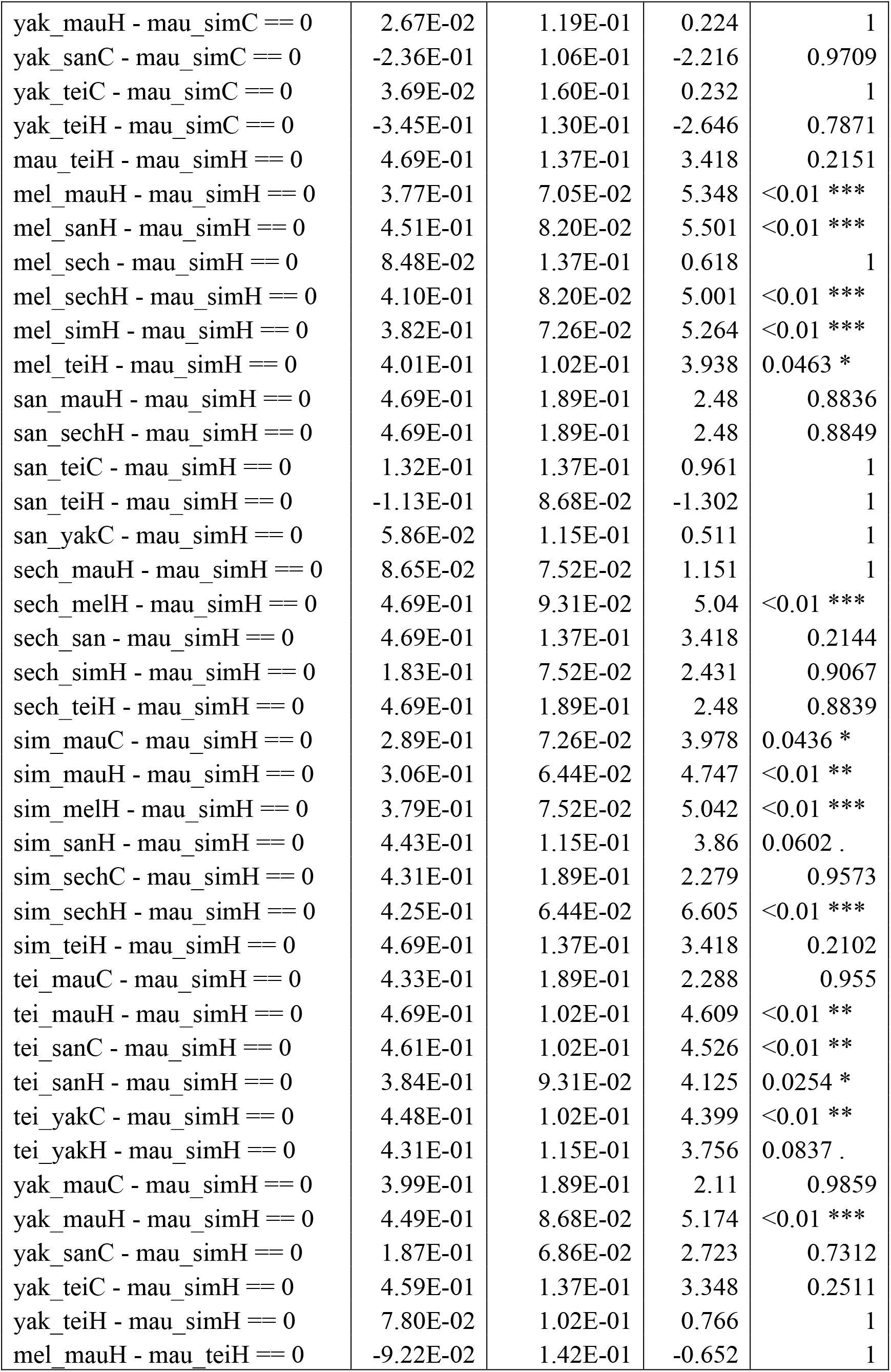
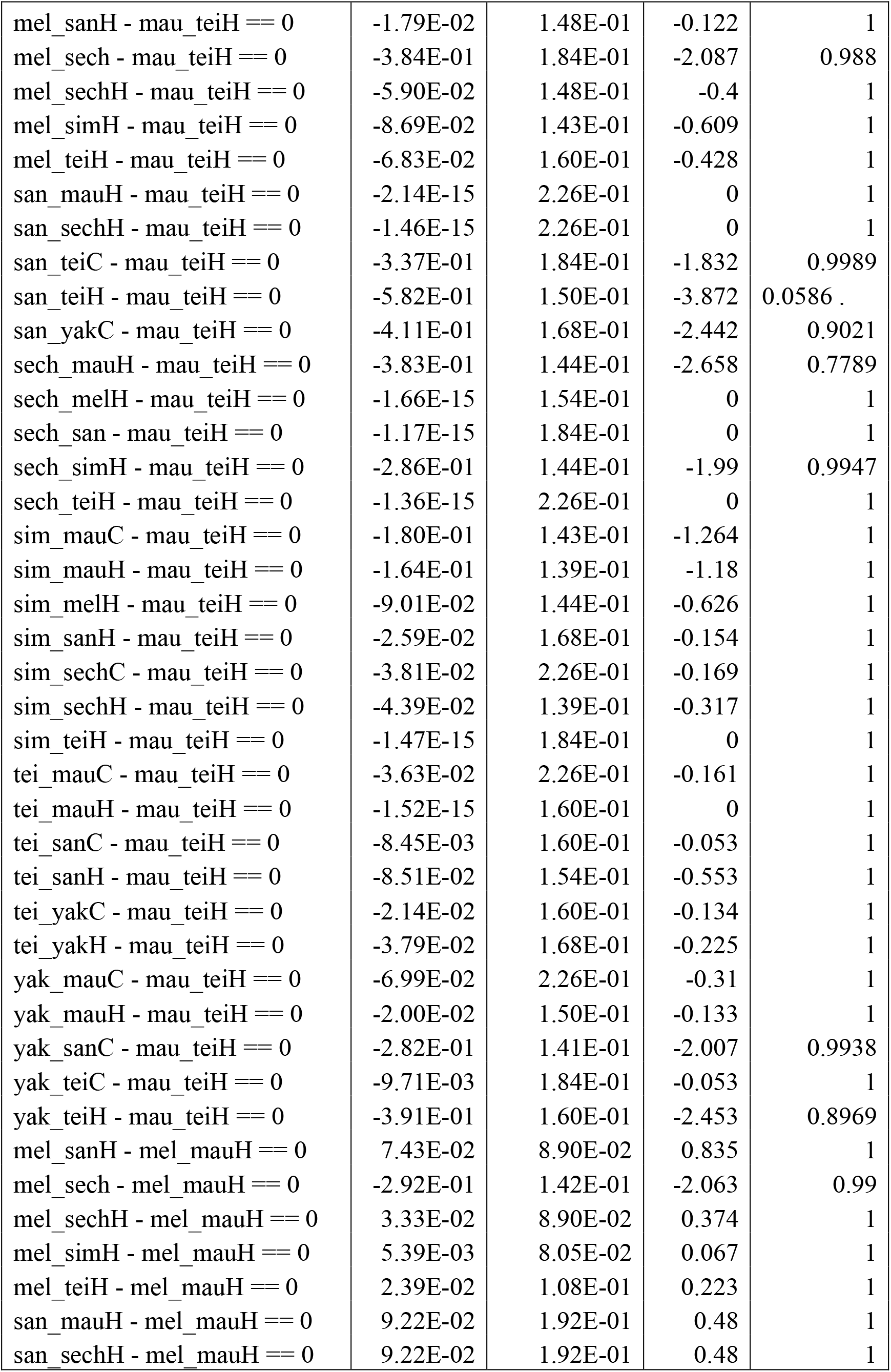
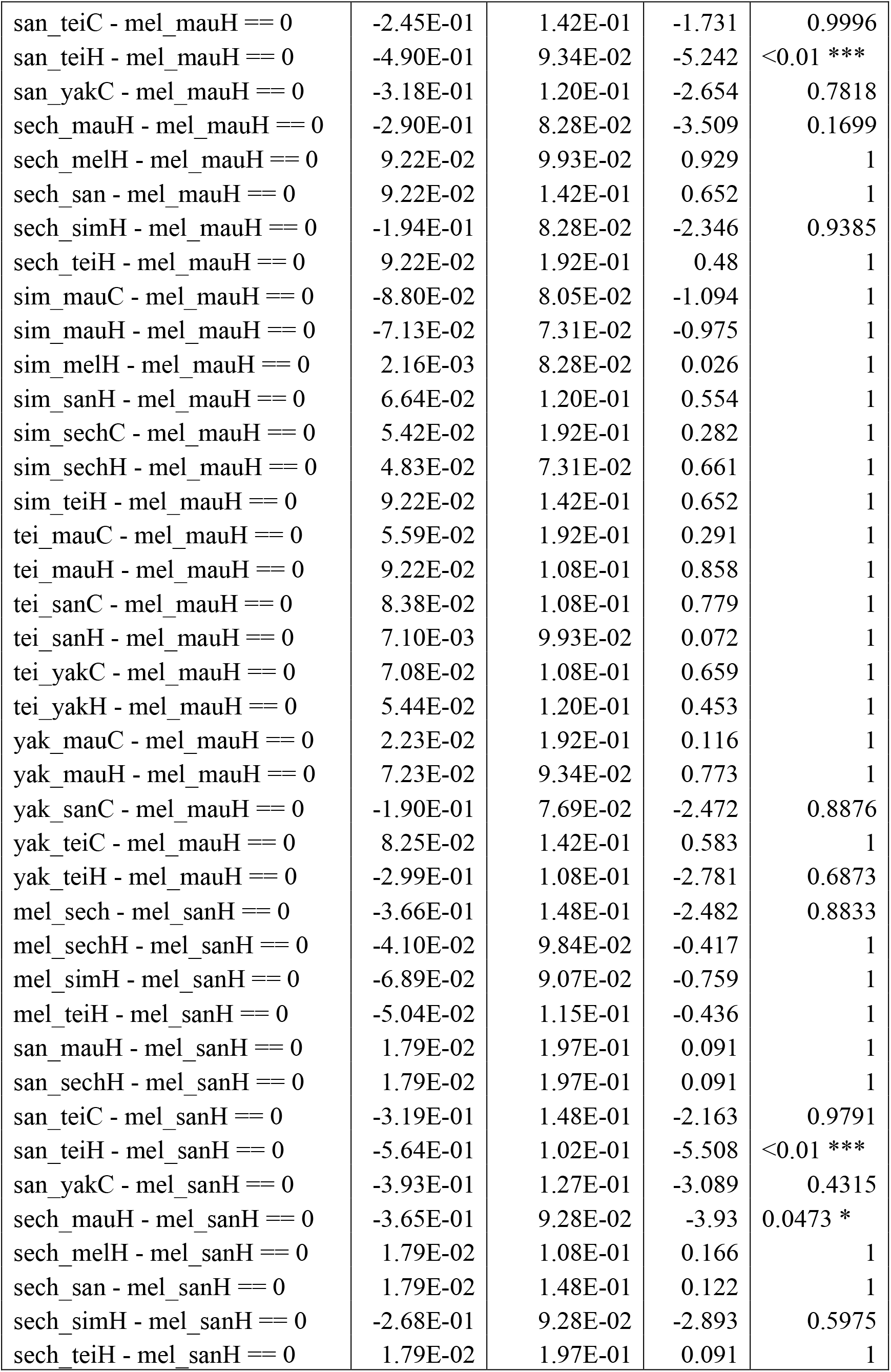
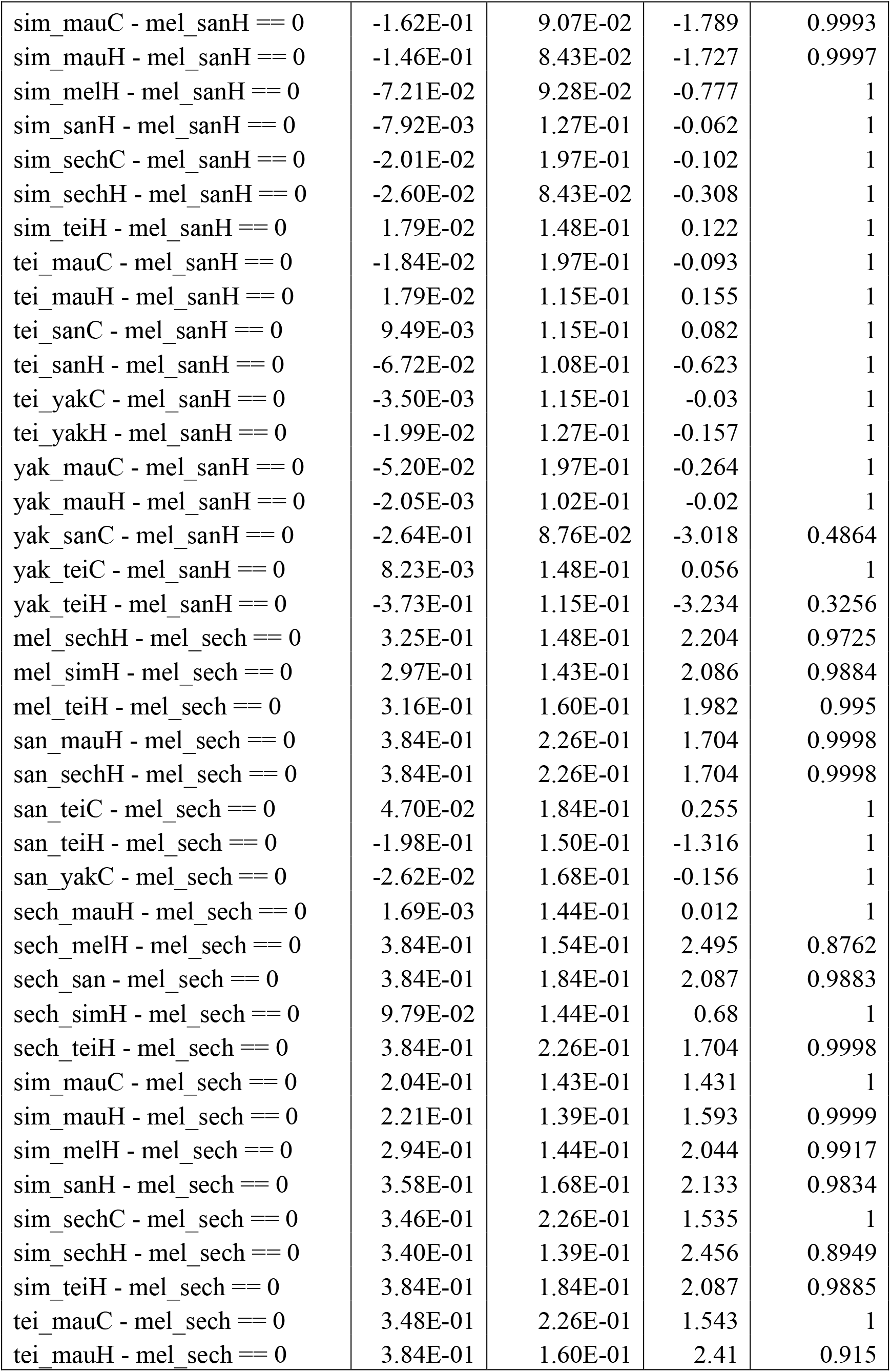
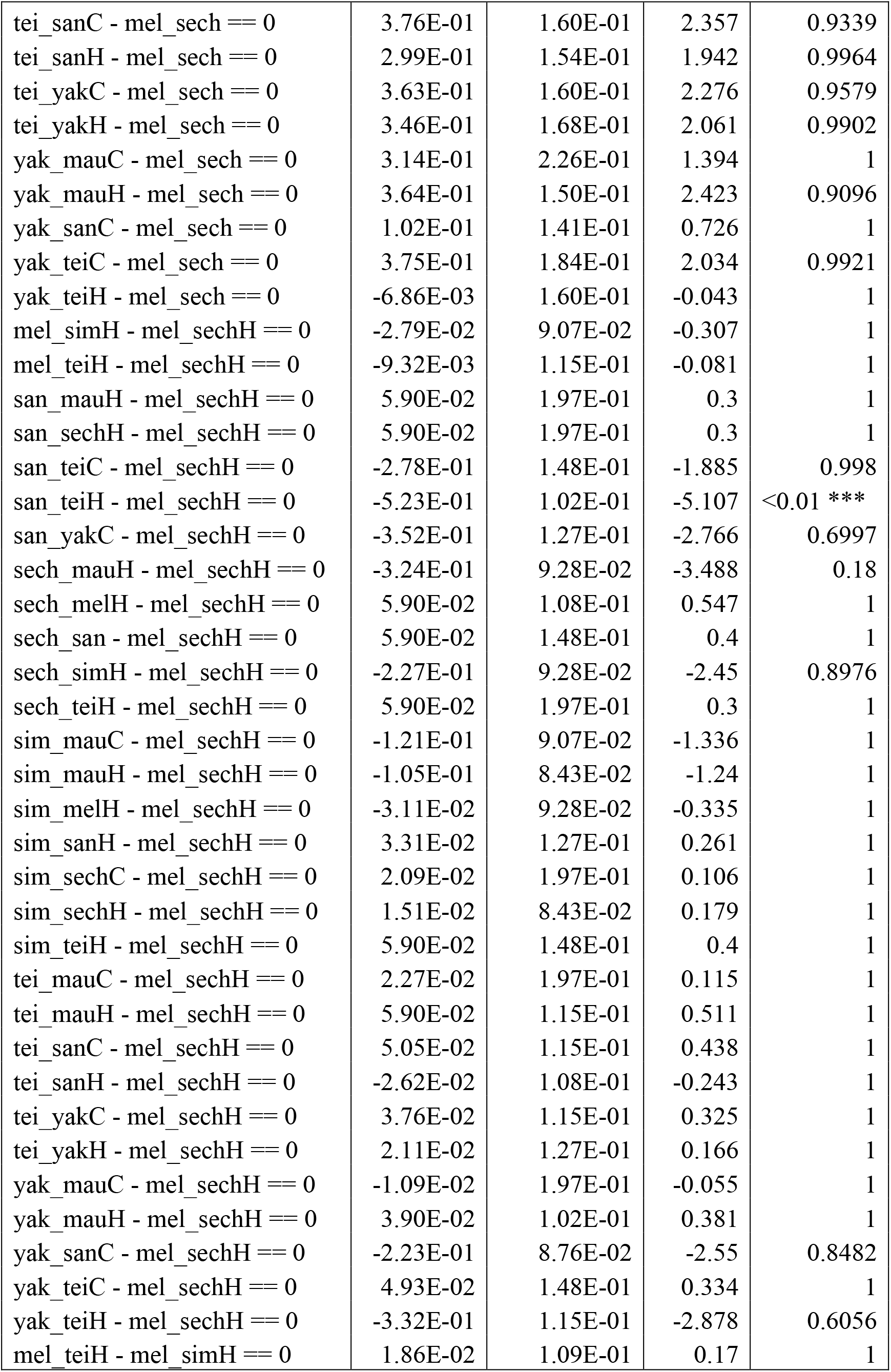
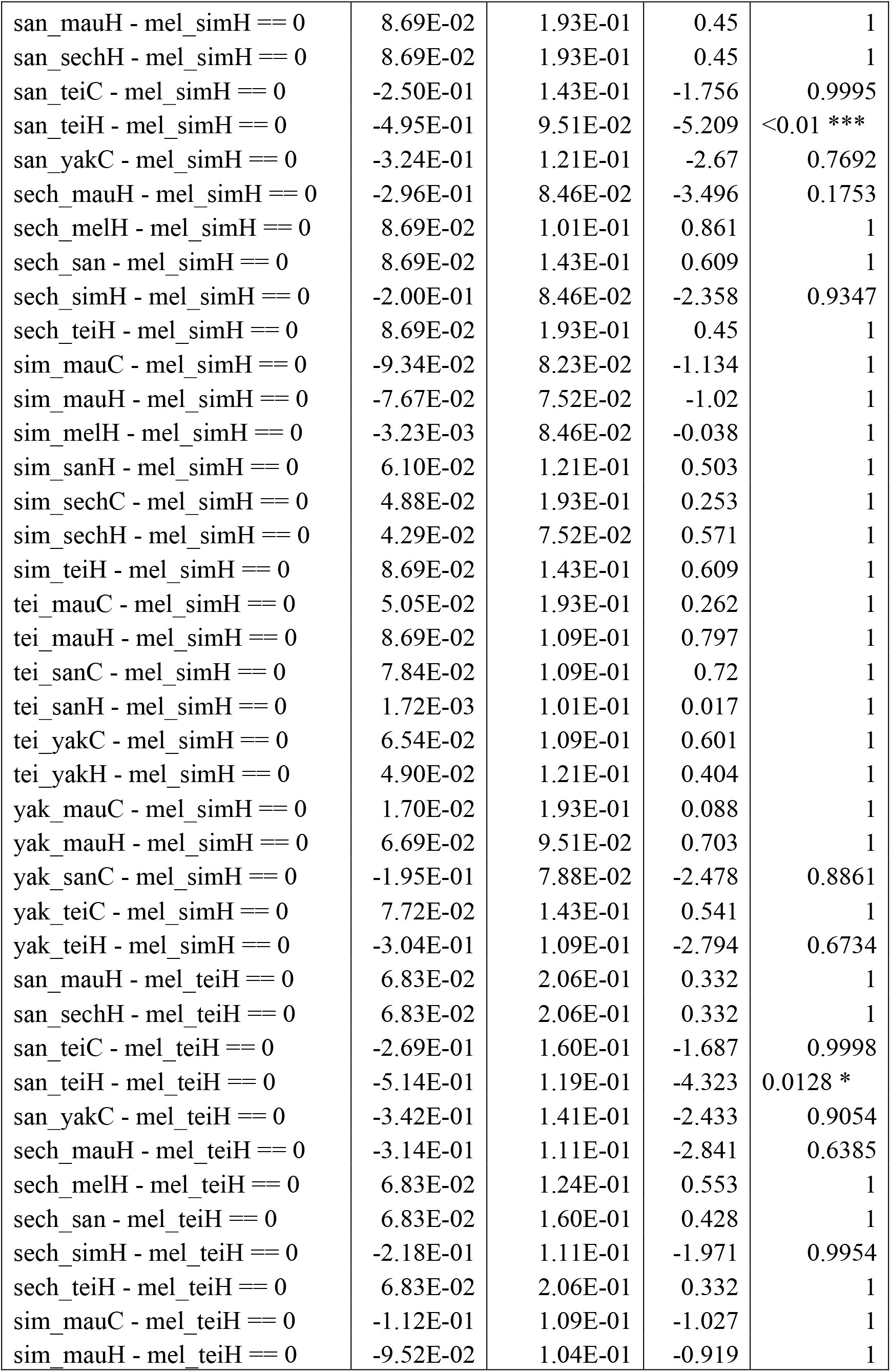
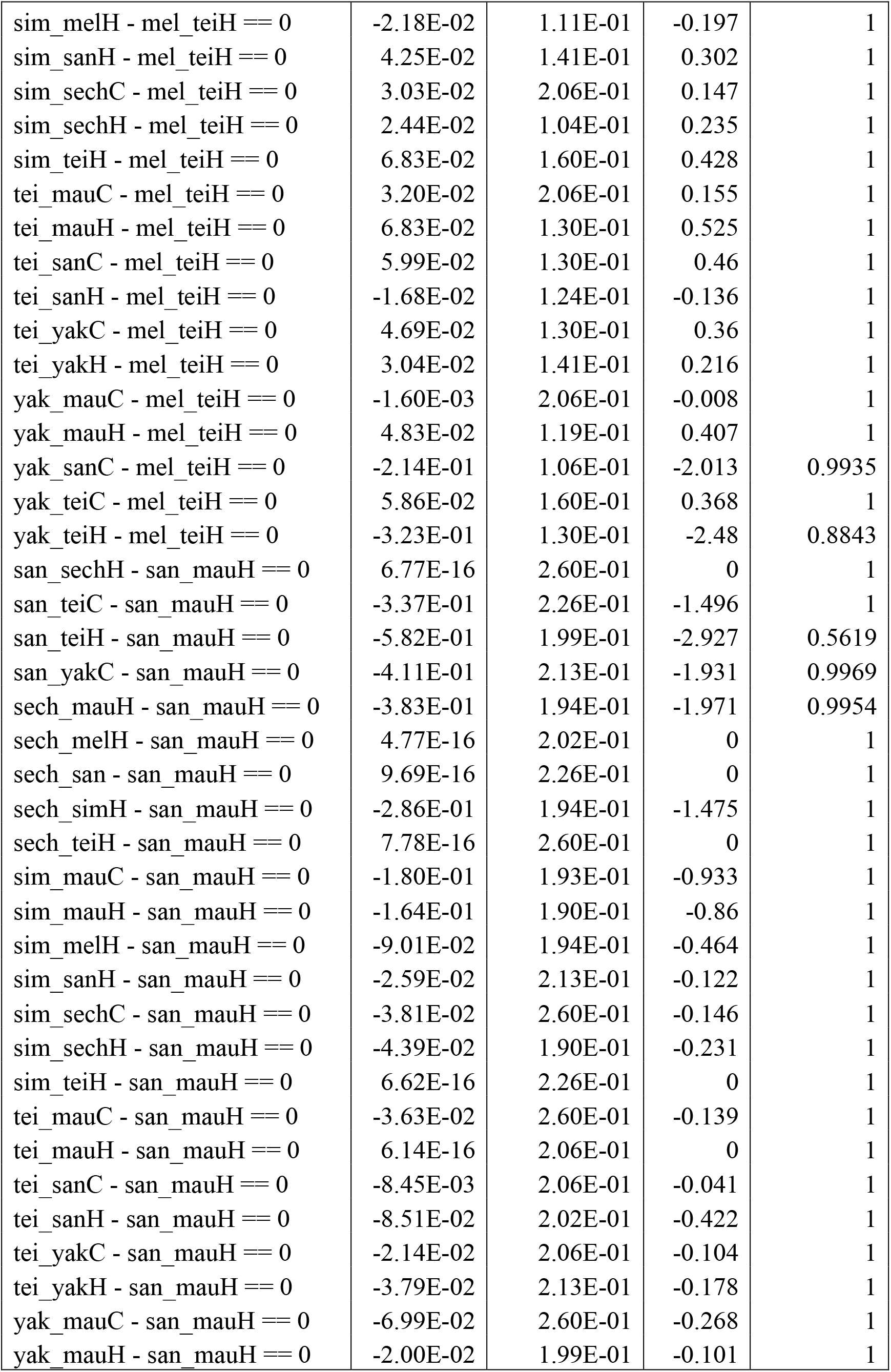
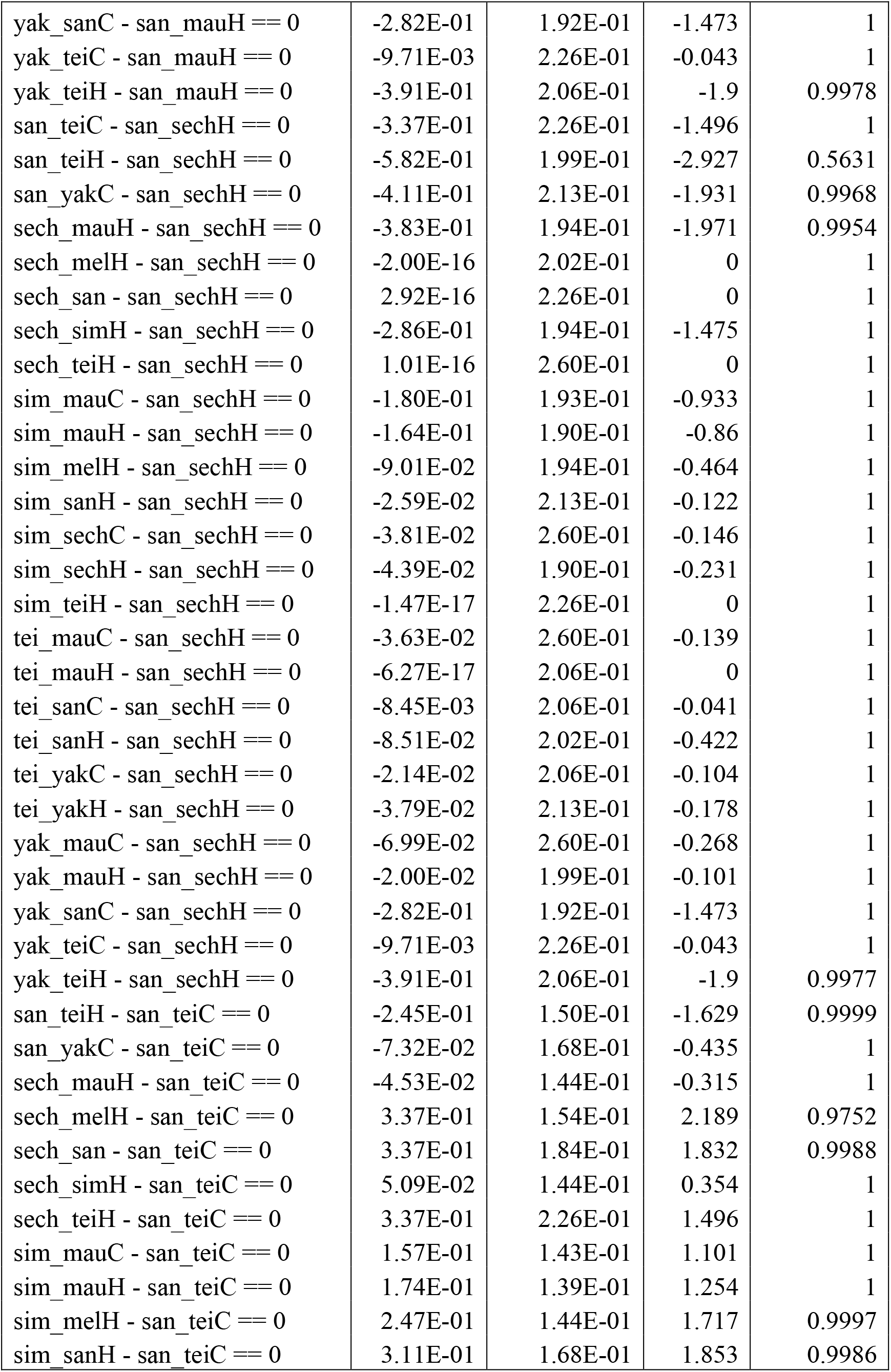
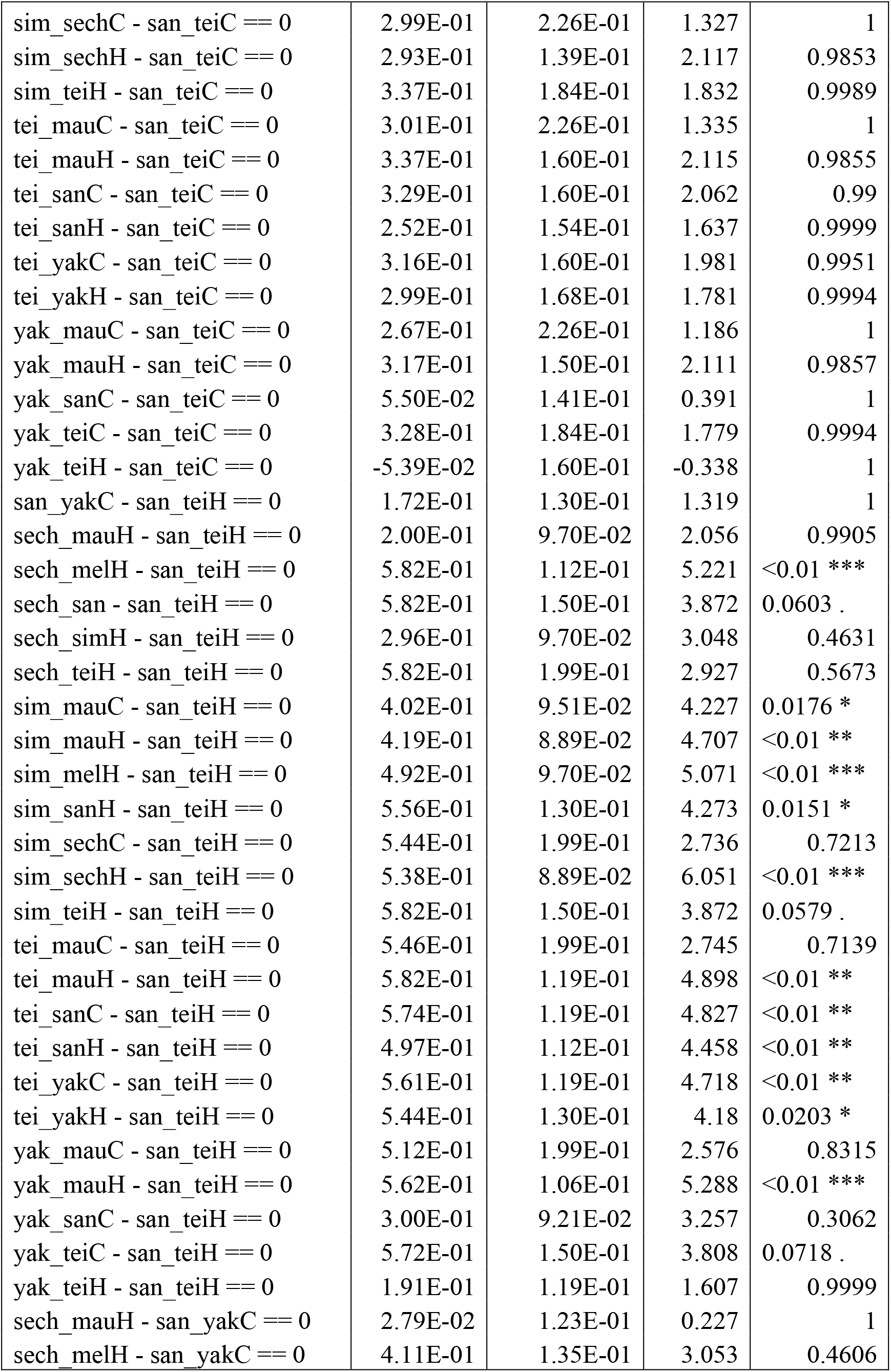
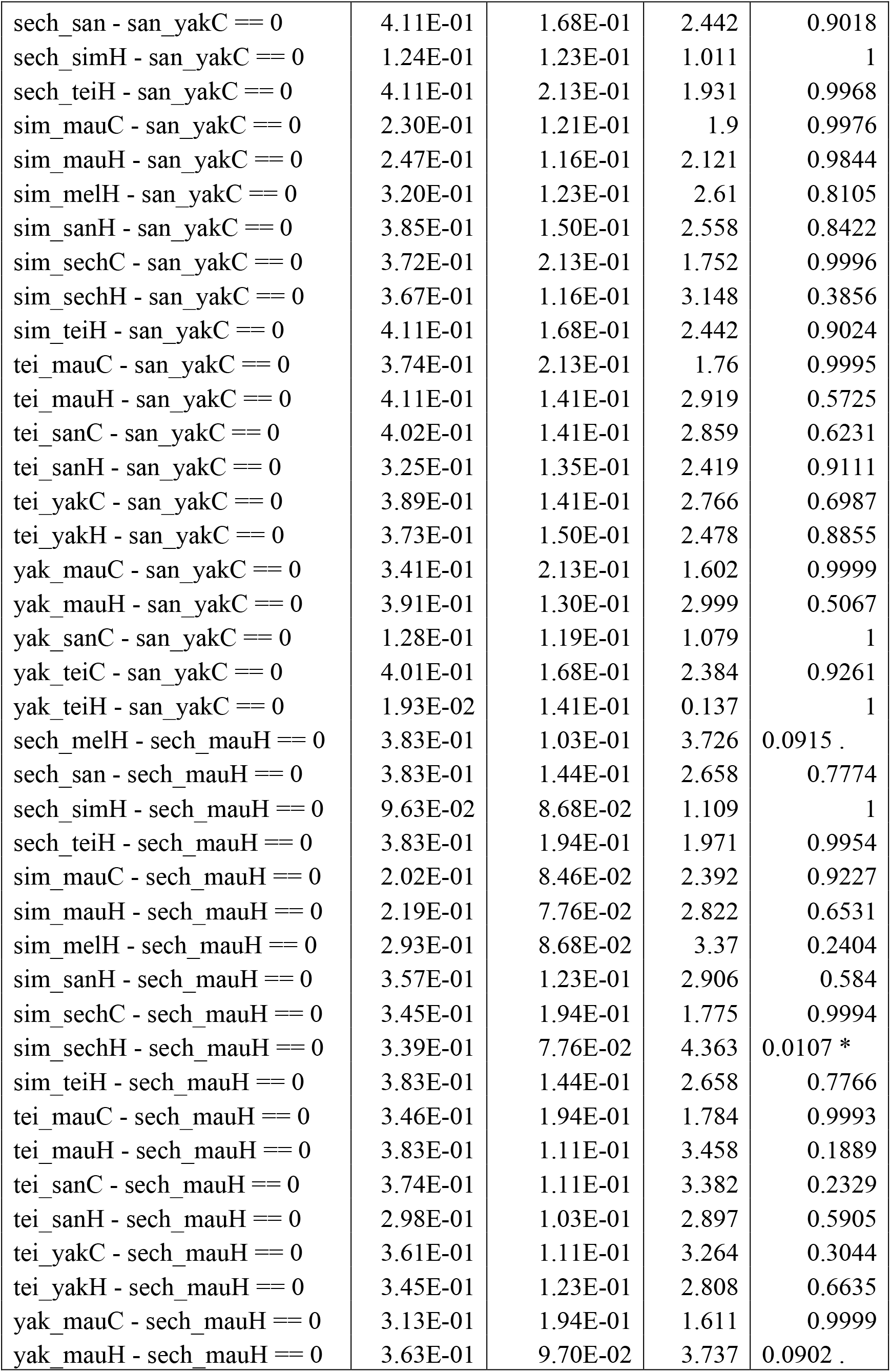
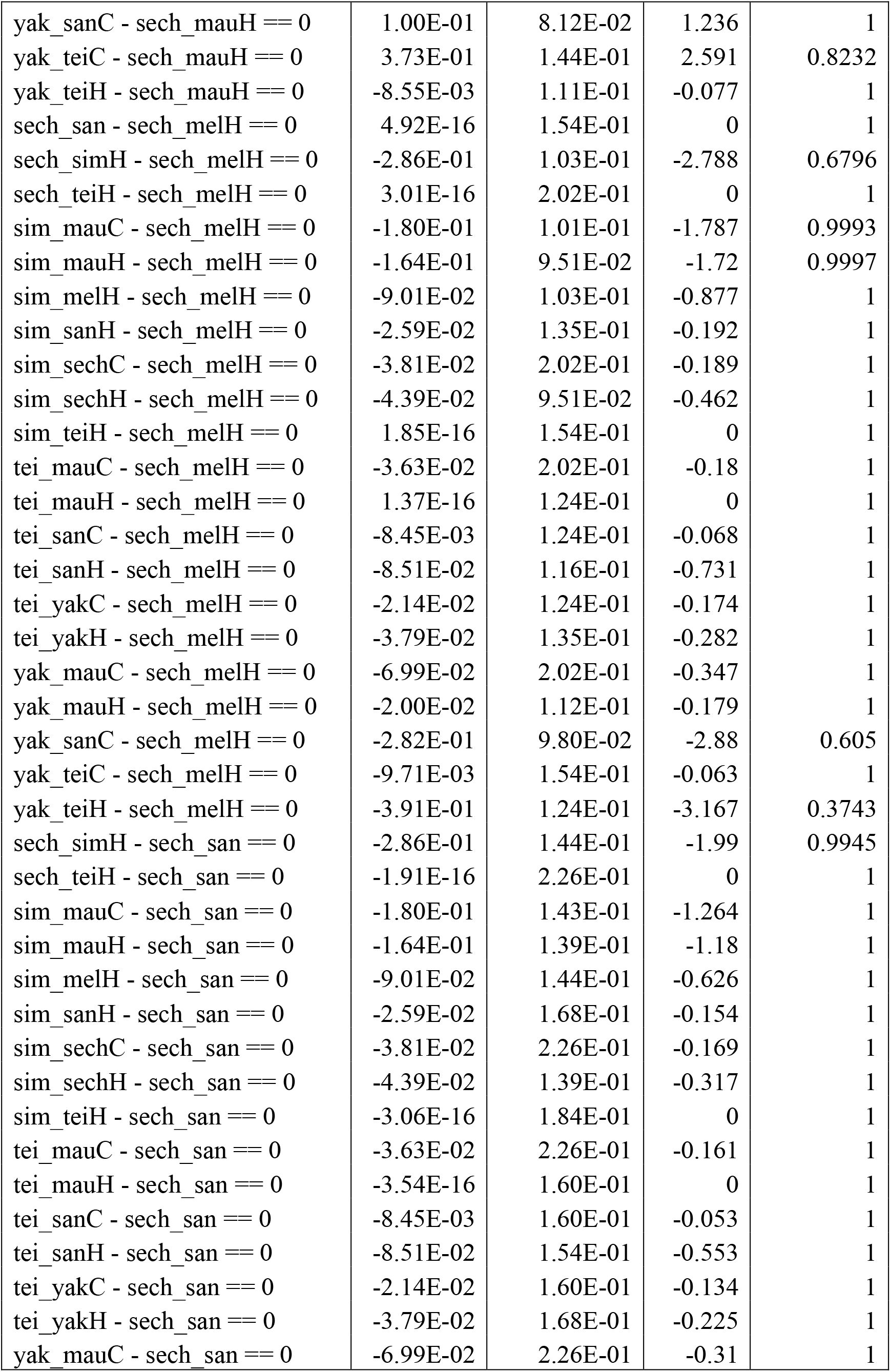
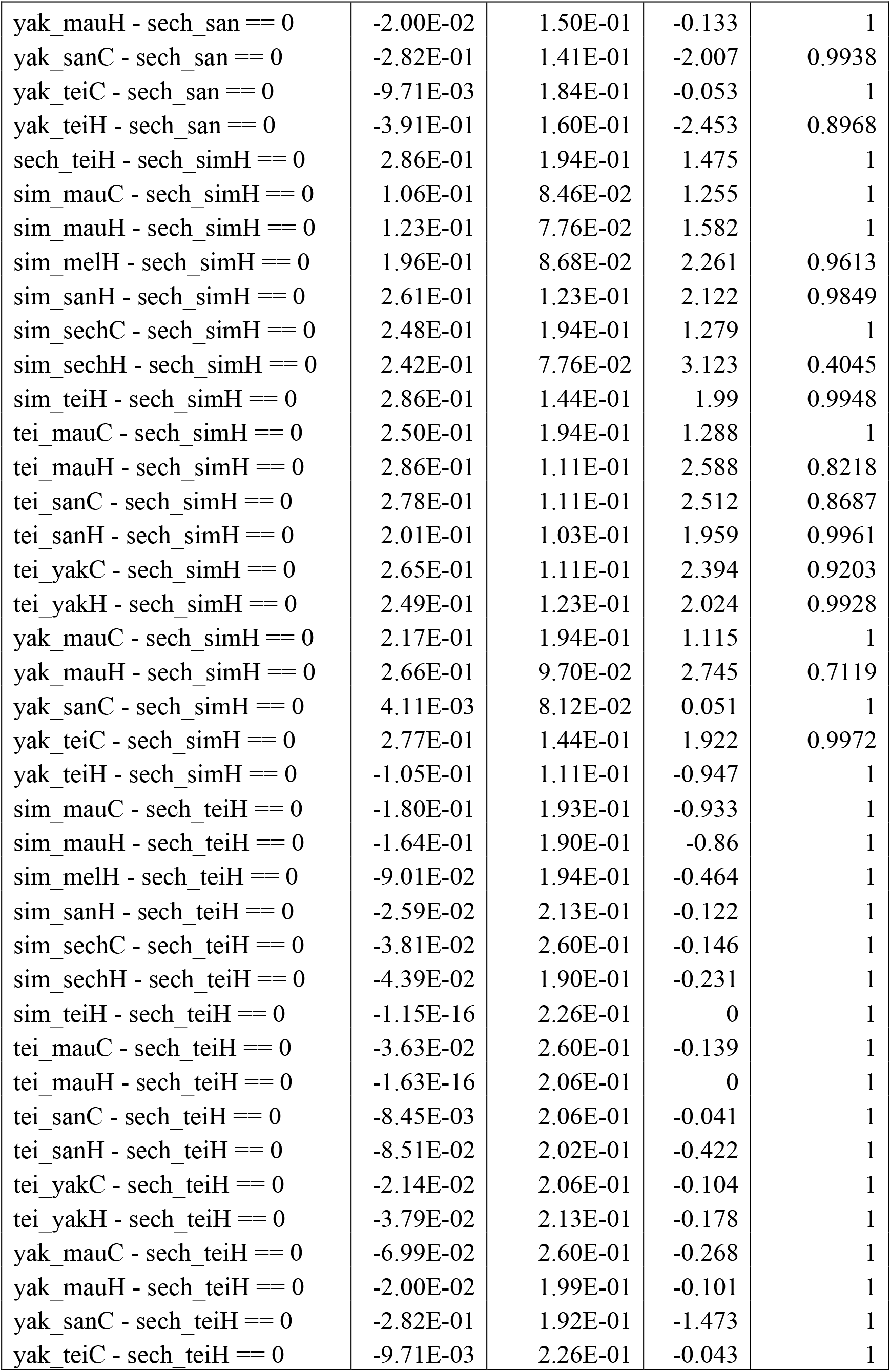
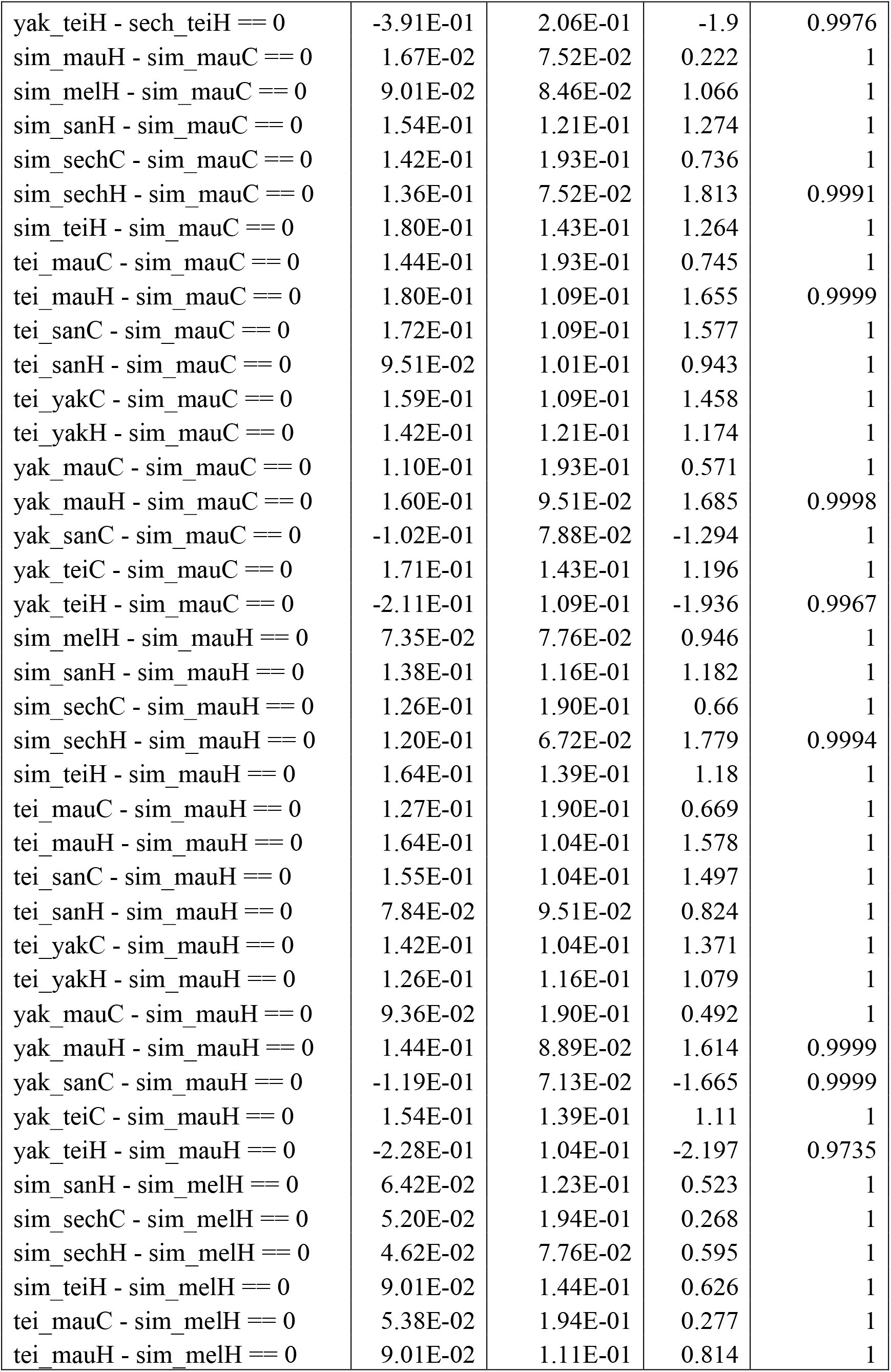
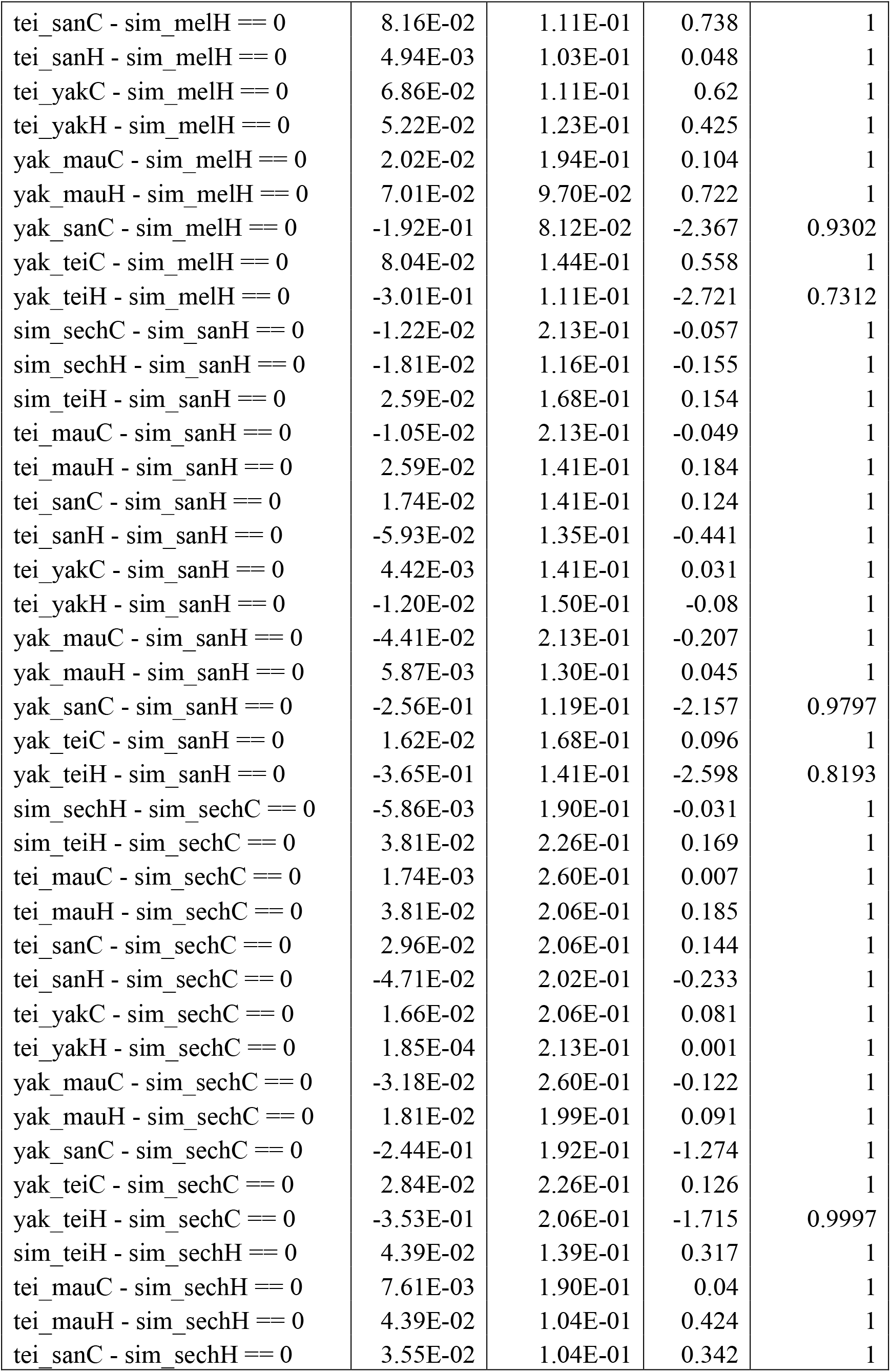
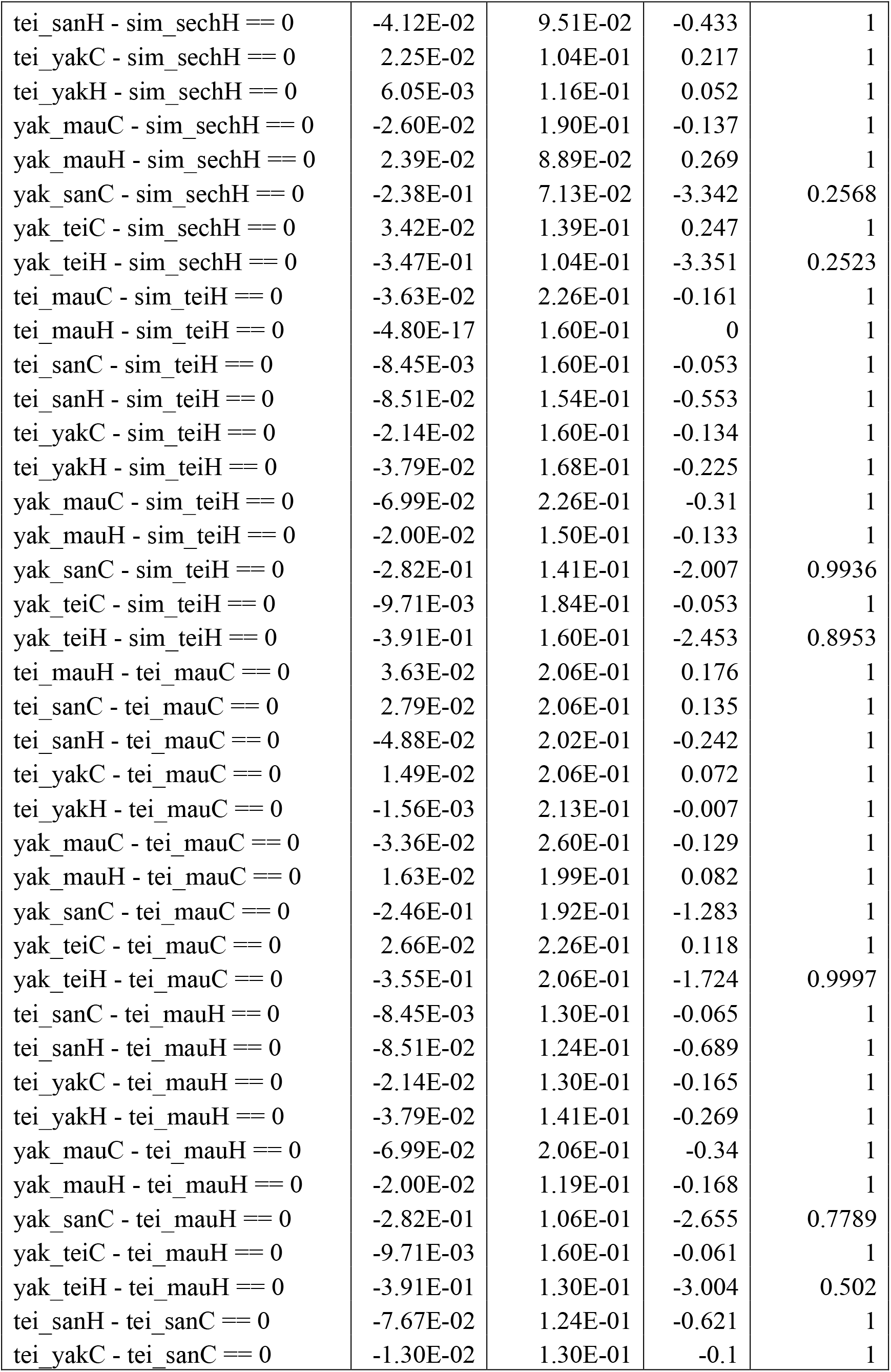
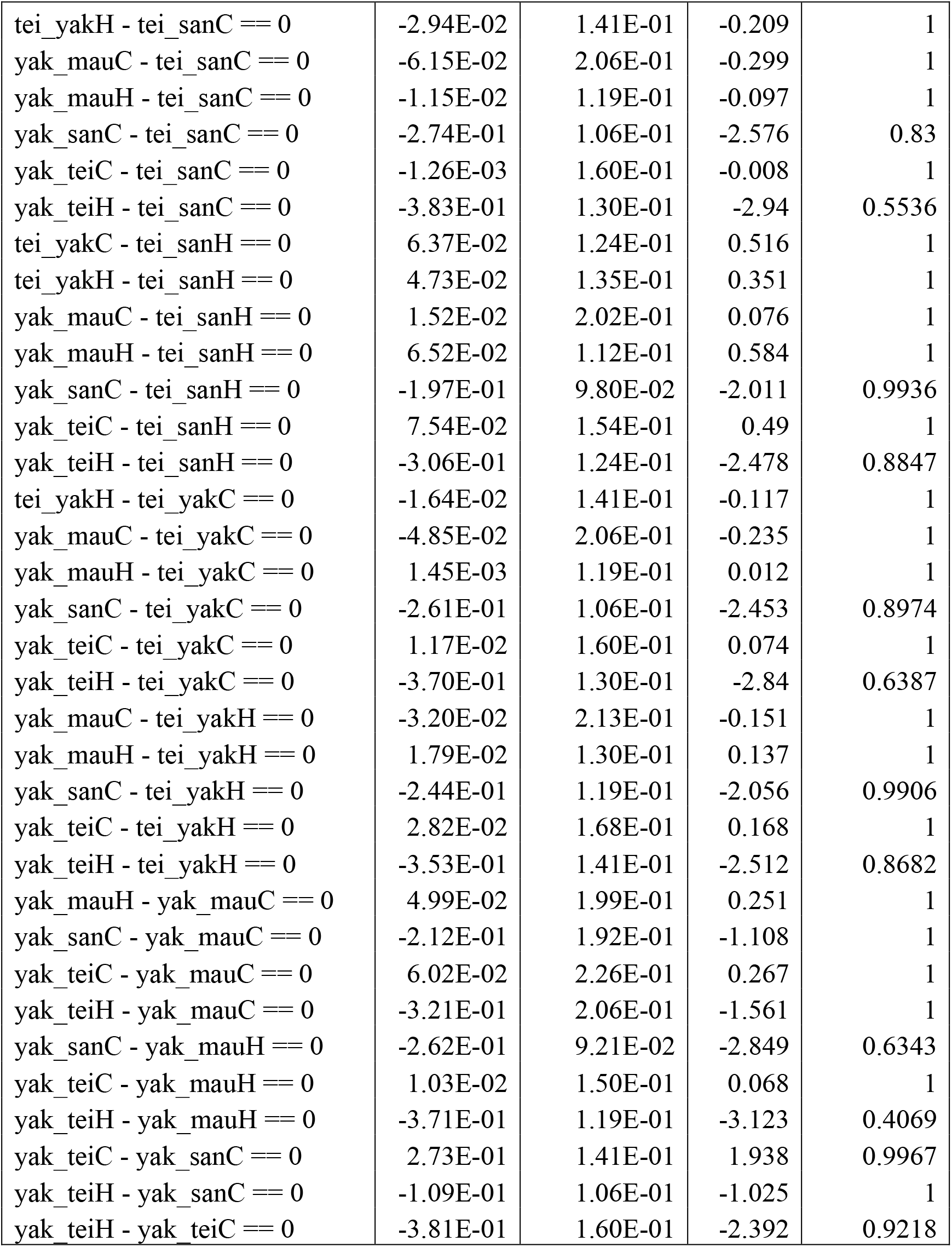
Pairwise comparisons between the magnitude of CSP in different crosses between species of the *melanogaster* subgroup. Each double mating is summarized by the first three letters of the genotype of the female, an underscore, and the first three letters of the genotype of the male. The C or H at the end of each term represents whether the first male was conspecific © or heterospecific (H). The number of degrees of freedom for each pairwise comparison equaled the total number of observations minus the number of means; df =187.

**TABLE S5.**
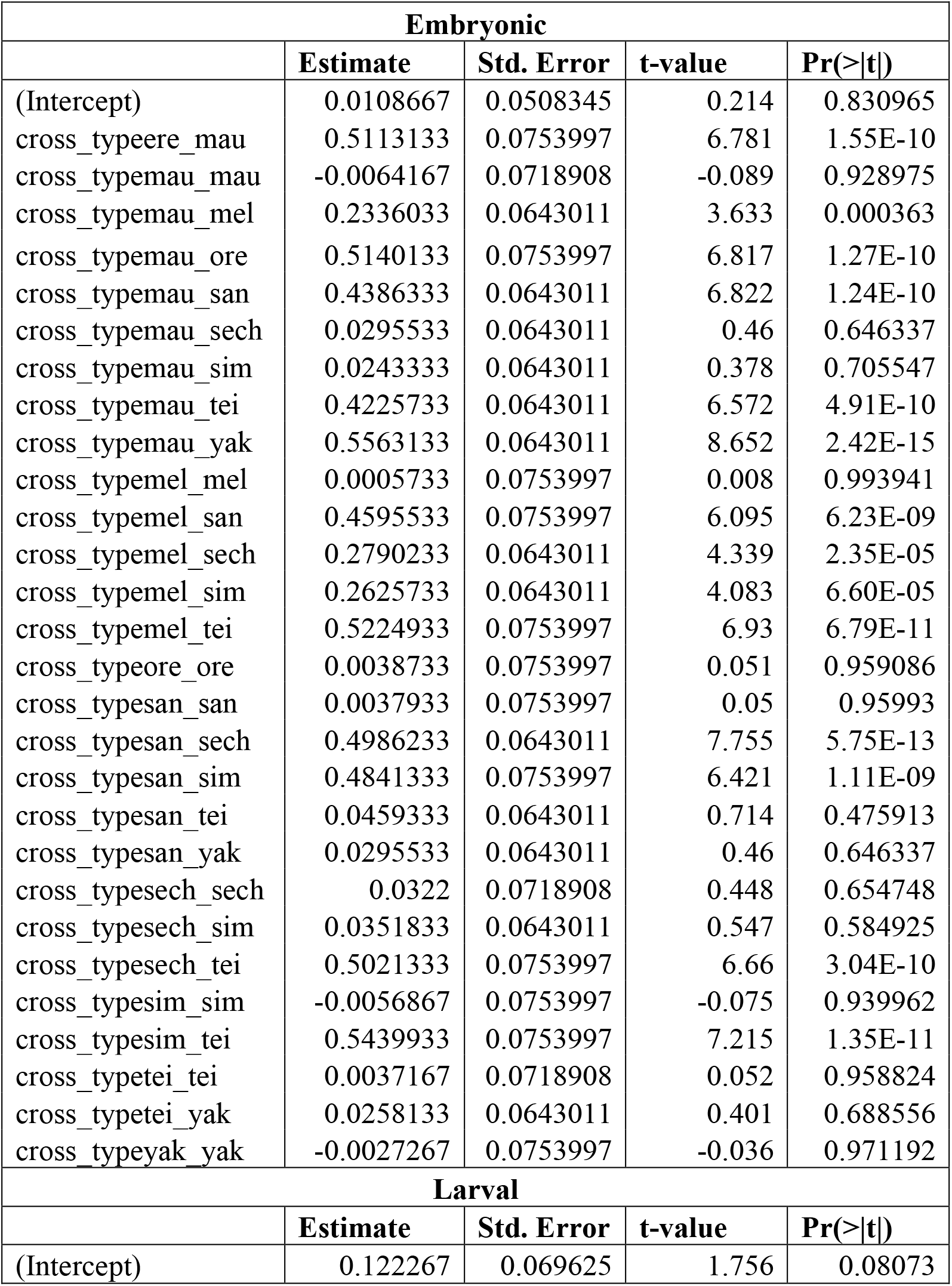
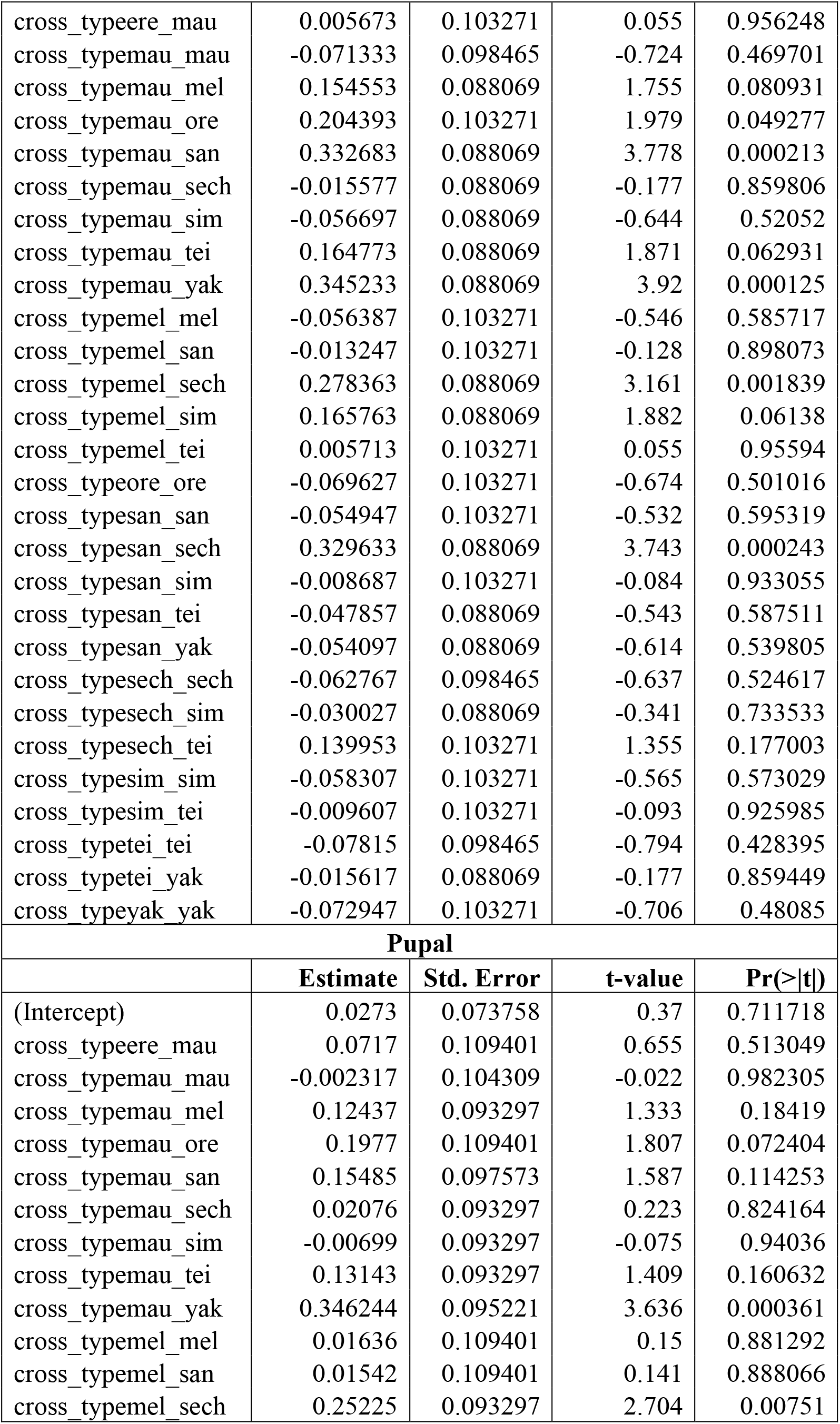
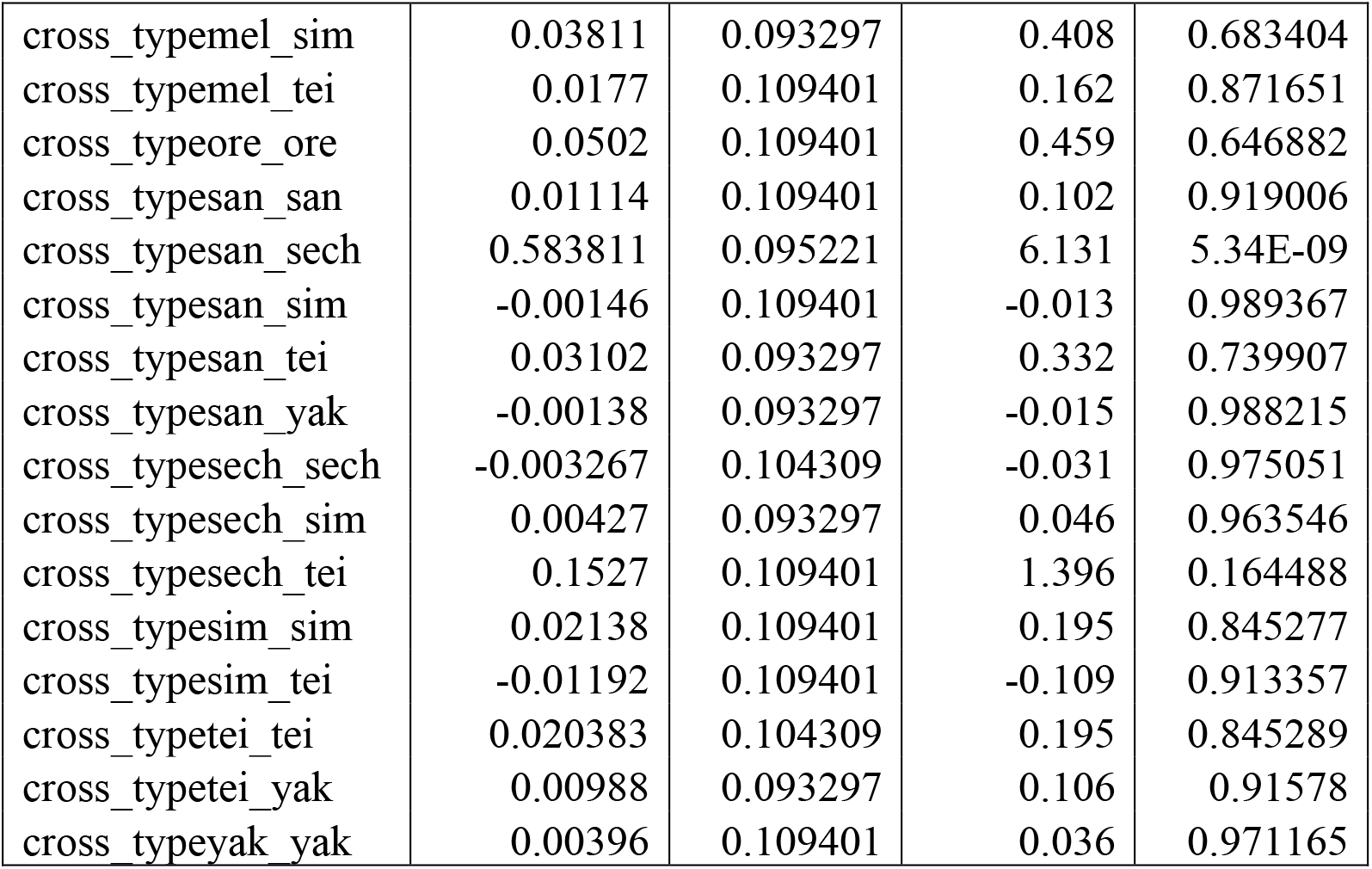
Linear contrasts for the magnitude of hybrid inviability at each of the three developmental stages (embryo, larvae, and pupae). The genotype of each cross is summarized by the first three letters of the genotype of the female, an underscore, and the first three letters of the genotype of the male. All linear contrasts were done using the number of degrees of freedom from the residuals of the linear model. df =132.

**TABLE S6.** K_s_, the number of per site synonymous substitutions between a pair of species was used as the average genetic distance between individuals of different species. Each species pair is summarized by showing the first three letters of one species, a dash, and the first three letters of the second species.

| **Species pair** | **$K_s$** |
| --- | --- |
| *ere - mau* | 0.2394 |
| *ere - mel* | 0.2641 |
| *ere - ore* | 0.0991 |
| *ere - san* | 0.2026 |
| *ere - sech* | 0.2568 |
| *ere - sim* | 0.2355 |
| *ere - tei* | 0.1901 |
| *ere - yak* | 0.1989 |
| *mau - mel* | 0.1086 |
| *mau - ore* | 0.2429 |
| *mau - san* | 0.2432 |
| *mau - sech* | 0.0381 |
| *mau - sim* | 0.0167 |
| *mau - tei* | 0.2307 |
| *mau - yak* | 0.2395 |
| *mel - ore* | 0.2676 |
| *mel - san* | 0.2679 |
| *mel - sech* | 0.126 |
| *mel - sim* | 0.1046 |
| *mel - tei* | 0.2553 |
| *mel - yak* | 0.2642 |
| *ore - san* | 0.2061 |
| *ore - sech* | 0.2603 |
| *ore - sim* | 0.239 |
| *ore - tei* | 0.1936 |
| *ore - yak* | 0.2024 |
| *san - sech* | 0.2607 |
| *san - sim* | 0.2393 |
| *san - tei* | 0.0936 |
| *san - yak* | 0.0344 |
| *sech - sim* | 0.0341 |
| *sech - tei* | 0.2481 |
| *sech - yak* | 0.257 |
| *sim - tei* | 0.2267 |
| *sim - yak* | 0.2356 |
| *tei - yak* | 0.0899 |

**TABLE S7.** McFadden pseudo r^2^ coefficients show the fit of logistic regressions for each RIM. We also show the regression coefficient of a linear regression for each case. In all cases we used logistic regressions instead of linear—even if the fit was slightly higher (e.g., CSP, Postzygotic isolation – embryonic viability) —because the logistic regressions make more biological sense (i.e., reproductive isolation cannot be higher than 1).

| Isolation | $r^2$ (Linear) | McFadden $r^2$ |
| --- | --- | --- |
| Premating | 0.6205 | 0.7704 |
| CSP | 0.3998 | 0.3667 |
| PMPZ | 0.4598 | 0.5372 |
| Postzygotic | 0.8264 | 0.7176 |
| Post - embryonic | 0.7994 | 0.7899 |
| Post - larval | 0.4058 | 0.4252 |
| Post - pupal | 0.2921 | 0.3525 |

**TABLE S8.** Pairwise comparison between four RI barriers in the *melanogaster* subgroup of species. The magnitude of variability within a RIM was calculated by subsampling 10,000 estimates and comparing their β1 value using a Mann-Whitney U test. Upper diagonal shows the Mann-Whitney test result. The lower diagonal is the empirical P-value of the observed value. The Wilcoxon result is significant in all six comparisons.

|  | Premating | NCGMI | CSP | Postzygotic |
| --- | --- | --- | --- | --- |
| Premating | * | 3,736,612.5 | 10,951,245 | 0 |
| NCGMI | $< 1 \times 10^{-15}$ | * | 33,648,753.5 | 2 |
| CSP | $< 1 \times 10^{-15}$ | $< 1 \times 10^{-15}$ | * | 0 |
| Postzygotic | $< 1 \times 10^{-15}$ | $< 1 \times 10^{-15}$ | $< 1 \times 10^{-15}$ | * |

**TABLE S9.** Testing for the phylogenetic signature of reinforcing selection by comparing the magnitude of in two species triad. Each triad is composed by a sympatric pair and an allopatric pair. If reinforcing selection has acted, the magnitude of RI in the sympatric pair should be larger than in the allopatric pair. Significance of the difference between means was assessed using permutation tests. *Drosophila yakuba* and *D. teissieri* are sympatric, while *D. santomea* and *D. teissieri* are allopatric. *Drosophila melanogaster* and *D. simulans* are sympatric, while *D. melanogaster* and *D. sechellia* are allopatric.

|  | *D. yakuba/D. santomea/D. teissieri* |  |  |  | *D. simulans/D. sechellia/D. melanogaster* |  |  |  |
| --- | --- | --- | --- | --- | --- | --- | --- | --- |
|  | Mean<br>*yakuba-*<br>*teissieri* | Mean<br>*santomea-*<br>*teissieri* | Z-value | P | Mean<br>*simulans-*<br>*melanogaster* | Mean<br>*sechellia-*<br>*melanogaster* | Z-value | P |
| Premating | 0.9990<br>(0.0031) | 1(0) | 1 | 1 | 1(0) | 1(0) | NA | NA |
| NCGI | 0.8915 | 0.8618 | -1.9542 | 0.0508 | 0.9156<br>(0.0678) | 0.9522<br>(0.0430) | 1.742 | 0.08181 |
| CSP | 0.9811<br>(0.0166) | 0.9398<br>(0.0914) | -1.5613 | 0.1331 | 0.9647<br>(0.0346) | 0.9625<br>(0.0787) | -0.135 | 0.9156 |
| Postzygic<br>isolation | 0.9077<br>(0.0843) | 0.9019<br>(0.0757) | -0.16471 | 0.8794 | 0.6995<br>(0.2720) | 0.5162<br>(0.3140) | -<br>1.3618 | 0.1799 |
| Postzygic<br>isolation:<br>embryonic<br>lethality | 0.9625<br>(0.0301) | 0.9457<br>(0.0619) | 0.75184 | 0.5815 | 0.2701<br>(0.2750) | 0.2738<br>(0.2246) | 0.0342 | 0.9668 |
| Postzygic<br>isolation:<br>larval<br>lethality | 0.9526<br>(0.0539) | 0.9443<br>(0.0404) | 0.40326 | 0.7126 | 0.7349<br>(0.2477) | 0.6569<br>(0.1717) | -<br>0.8261 | 0.418 |
| Postzygic<br>isolation:<br>pupal<br>lethality | 0.9443<br>(0.0547) | 0.9526<br>(0.0472) | -0.6155 | 0.5528 | 0.9346<br>(0.0722) | 0.5418<br>(0.4310) | -2.14 | 0.0296 |

**TABLE S10.** Pairwise comparison between four RI barriers in the *Drosophila* genus using only phylogenetically independent crosses. Conventions are the same as in Table S10.

|  | Premating | NCGMI | CSP | Postzygotic |
| --- | --- | --- | --- | --- |
| Premating | * | 0 | 4,636,038.5 | 0 |
| NCGMI | $< 1 \times 10^{-15}$ | * | 0 | 814,100.5 |
| CSP | $< 1 \times 10^{-15}$ | $< 1 \times 10^{-15}$ | * | 0 |
| Postzygotic | $< 1 \times 10^{-15}$ | $< 1 \times 10^{-15}$ | $< 1 \times 10^{-15}$ | * |

**TABLE S11.**
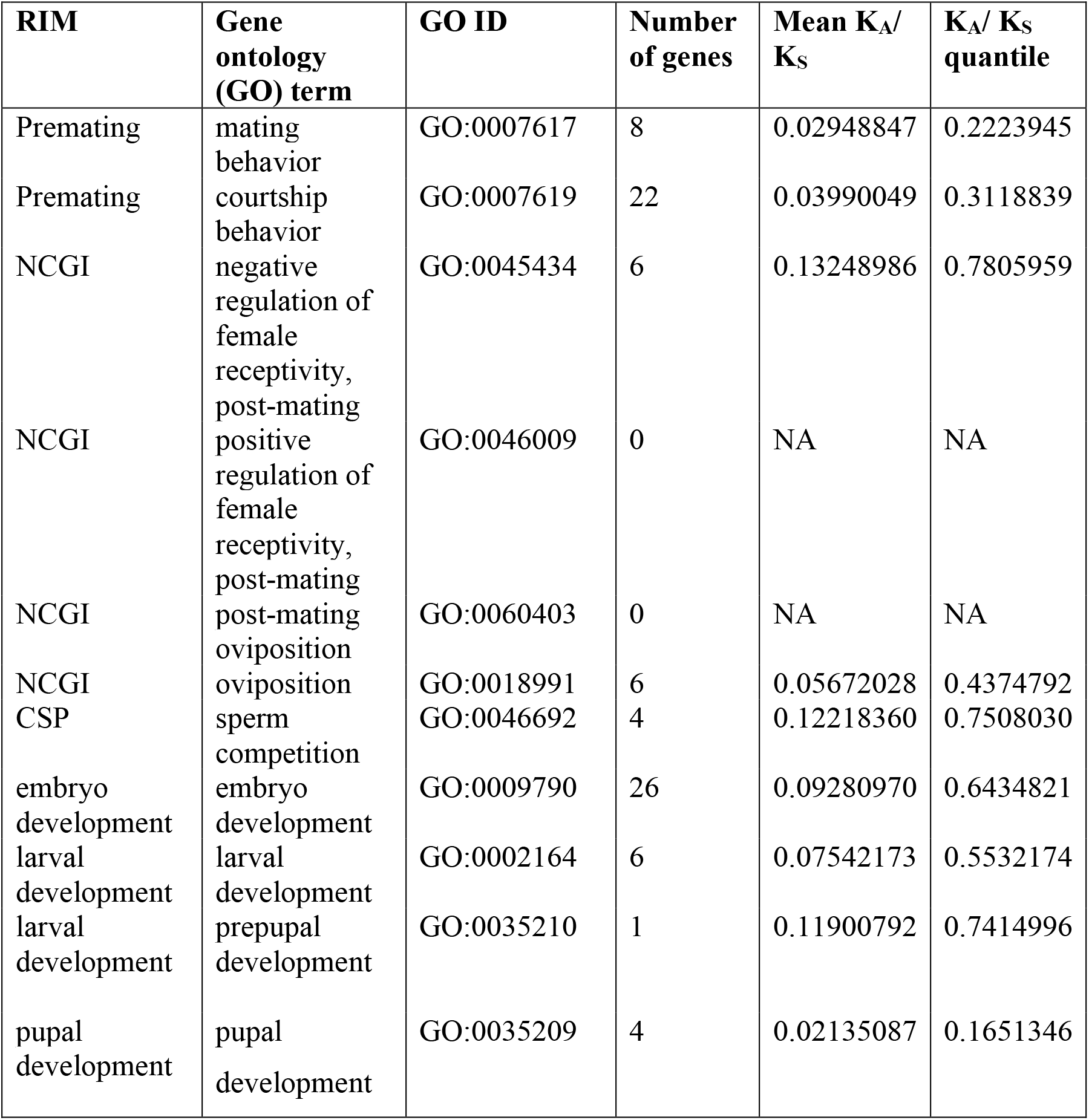
No evidence for pervasive selection at any GO term. We calculated the mean K_A_/ K_S_ for eleven GO terms associated to Reproductive isolating mechanisms. No GO term showed a notably large K_A_/ K_S_ value.

**TABLE S12.** Raw K_A_/ K_S_ estimates for all genes included in the GO analyses shown in Table S11.

| GO term number | GO term name | Gene | sequence length | $K_A/K_S$ |
| --- | --- | --- | --- | --- |
| GO:0002164 | larval development | CG5786 | 1353 | 0.037291095 |
| GO:0002164 | larval development | CG4620 | 1782 | 0.037665947 |
| GO:0002164 | larval development | CG44533 | 1587 | 0.044492441 |
| GO:0002164 | larval development | CG4141 | 3216 | 0.057672849 |
| GO:0002164 | larval development | CG1886 | 3612 | 0.06295307 |
| GO:0002164 | larval development | CG5671 | 267 | 0.212454986 |
| GO:0007617 | mating behavior | CG1725 | 18 | NA |
| GO:0007617 | mating behavior | CG8556 | 567 | 0 |
| GO:0007617 | mating behavior | CG32498 | 675 | 0.010807736 |
| GO:0007617 | mating behavior | CG14307 | 1458 | 0.013414634 |
| GO:0007617 | mating behavior | CG9533 | 4965 | 0.031213192 |
| GO:0007617 | mating behavior | CG9019 | 1974 | 0.040394248 |
| GO:0007617 | mating behavior | CG2647 | 1752 | 0.054991608 |
| GO:0007617 | mating behavior | CG3234 | 1818 | 0.055597867 |
| GO:0007619 | courtship behavior | CG12348 | 858 | 0 |
| GO:0007619 | courtship behavior | CG18069 | 936 | 0 |
| GO:0007619 | courtship behavior | CG8049 | 1365 | 0 |
| GO:0007619 | courtship behavior | CG10952 | 1002 | 0.003329634 |
| GO:0007619 | courtship behavior | CG32498 | 675 | 0.010807736 |
| GO:0007619 | courtship behavior | CG11094 | 669 | 0.011882998 |
| GO:0007619 | courtship behavior | CG4443 | 375 | 0.017262339 |
| GO:0007619 | courtship behavior | CG10118 | 1707 | 0.018227009 |
| GO:0007619 | courtship behavior | CG14039 | 1059 | 0.020346495 |
| GO:0007619 | courtship behavior | CG12073 | 1635 | 0.026519667 |
| GO:0007619 | courtship behavior | CG17228 | 3930 | 0.030338052 |
| GO:0007619 | courtship behavior | CG9533 | 4965 | 0.031213192 |
| GO:0007619 | courtship behavior | CG7925 | 363 | 0.032432433 |
| GO:0007619 | courtship behavior | CG10697 | 1314 | 0.033054914 |
| GO:0007619 | courtship behavior | CG8428 | 1086 | 0.036617262 |
| GO:0007619 | courtship behavior | CG6727 | 516 | 0.038233635 |
| GO:0007619 | courtship behavior | CG9019 | 1974 | 0.040394248 |
| GO:0007619 | courtship behavior | CG43368 | 4329 | 0.042032622 |
| GO:0007619 | courtship behavior | CG2647 | 1752 | 0.054991608 |
| GO:0007619 | courtship behavior | CG6070 | 1533 | 0.06211334 |
| GO:0007619 | courtship behavior | CG12390 | 1356 | 0.126022229 |
| GO:0007619 | courtship behavior | CG6917 | 1503 | 0.241991277 |
| GO:0009790 | embryo development | CG10293 | 1092 | 0.01226158 |
| GO:0009790 | embryo development | CG4533 | 555 | 0.026546865 |
| GO:0009790 | embryo development | CG11949 | 2040 | 0.02670692 |
| GO:0009790 | embryo development | CG43140 | 3303 | 0.029211956 |
| GO:0009790 | embryo development | CG8597 | 924 | 0.031774853 |
| GO:0009790 | embryo development | CG9842 | 582 | 0.046695794 |
| GO:0009790 | embryo development | CG6146 | 1374 | 0.046801872 |
| GO:0009790 | embryo development | CG4894 | 5319 | 0.048833189 |
| GO:0009790 | embryo development | CG3668 | 1317 | 0.052054795 |
| GO:0009790 | embryo development | CG11921 | 1098 | 0.055929782 |
| GO:0009790 | embryo development | CG5370 | 789 | 0.056584512 |
| GO:0009790 | embryo development | CG44436 | 3741 | 0.065331167 |
| GO:0009790 | embryo development | CG1945 | 7932 | 0.07361516 |
| GO:0009790 | embryo development | CG2950 | 1350 | 0.075516224 |
| GO:0009790 | embryo development | CG9885 | 1650 | 0.083745876 |
| GO:0009790 | embryo development | CG4035 | 729 | 0.08454927 |
| GO:0009790 | embryo development | CG11922 | 789 | 0.08603085 |
| GO:0009790 | embryo development | CG2189 | 1656 | 0.096002336 |
| GO:0009790 | embryo development | CG11129 | 1044 | 0.103975059 |
| GO:0009790 | embryo development | CG18402 | 6189 | 0.116952157 |
| GO:0009790 | embryo development | CG4965 | 1257 | 0.145536597 |
| GO:0009790 | embryo development | CG1264 | 627 | 0.15983843 |
| GO:0009790 | embryo development | CG9450 | 7299 | 0.165276432 |
| GO:0009790 | embryo development | CG9936 | 771 | 0.196078432 |
| GO:0009790 | embryo development | CG42572 | 1002 | 0.25483746 |
| GO:0009790 | embryo development | CG32562 | 3501 | 0.272364589 |
| GO:0018991 | oviposition | CG30446 | 1578 | 0.008635831 |
| GO:0018991 | oviposition | CG9019 | 1974 | 0.040394248 |
| GO:0018991 | oviposition | CG32484 | 1926 | 0.045749705 |
| GO:0018991 | oviposition | CG10733 | 690 | 0.068277715 |
| GO:0018991 | oviposition | CG7111 | 327 | 0.087740386 |
| GO:0018991 | oviposition | CG8887 | 6477 | 0.089523809 |
| GO:0035209 | pupal development | CG7998 | 996 | 0.007905983 |
| GO:0035209 | pupal development | CG3411 | 1035 | 0.013872374 |
| GO:0035209 | pupal development | CG12021 | 2520 | 0.021079881 |
| GO:0035209 | pupal development | CG7999 | 2934 | 0.042545251 |
| GO:0035210 | prepupal development | CG13176 | 1446 | 0.119007921 |
| GO:0045434 | negative regulation of female receptivity, post-mating | CG9659 | 1356 | 0.007367926 |
| GO:0045434 | negative regulation of female receptivity, post-mating | CG16752 | 723 | 0.020238621 |
| GO:0045434 | negative regulation of female receptivity, post-mating | CG7005 | 1773 | 0.043427621 |
| GO:0045434 | negative regulation of female receptivity, post-mating | CG5630 | 243 | 0.118793208 |
| GO:0045434 | negative regulation of female receptivity, post-mating | CG12558 | 861 | 0.255361268 |
| GO:0045434 | negative regulation of female receptivity, post-mating | CG4605 | 663 | 0.349750489 |
| GO:0046692 | sperm competition | CG9156 | 894 | 0.007387815 |
| GO:0046692 | sperm competition | CG17575 | 858 | 0.065161179 |
| GO:0046692 | sperm competition | CG1652 | 897 | 0.156889733 |
| GO:0046692 | sperm competition | CG1656 | 777 | 0.259295662 |

**TABLE S13.**
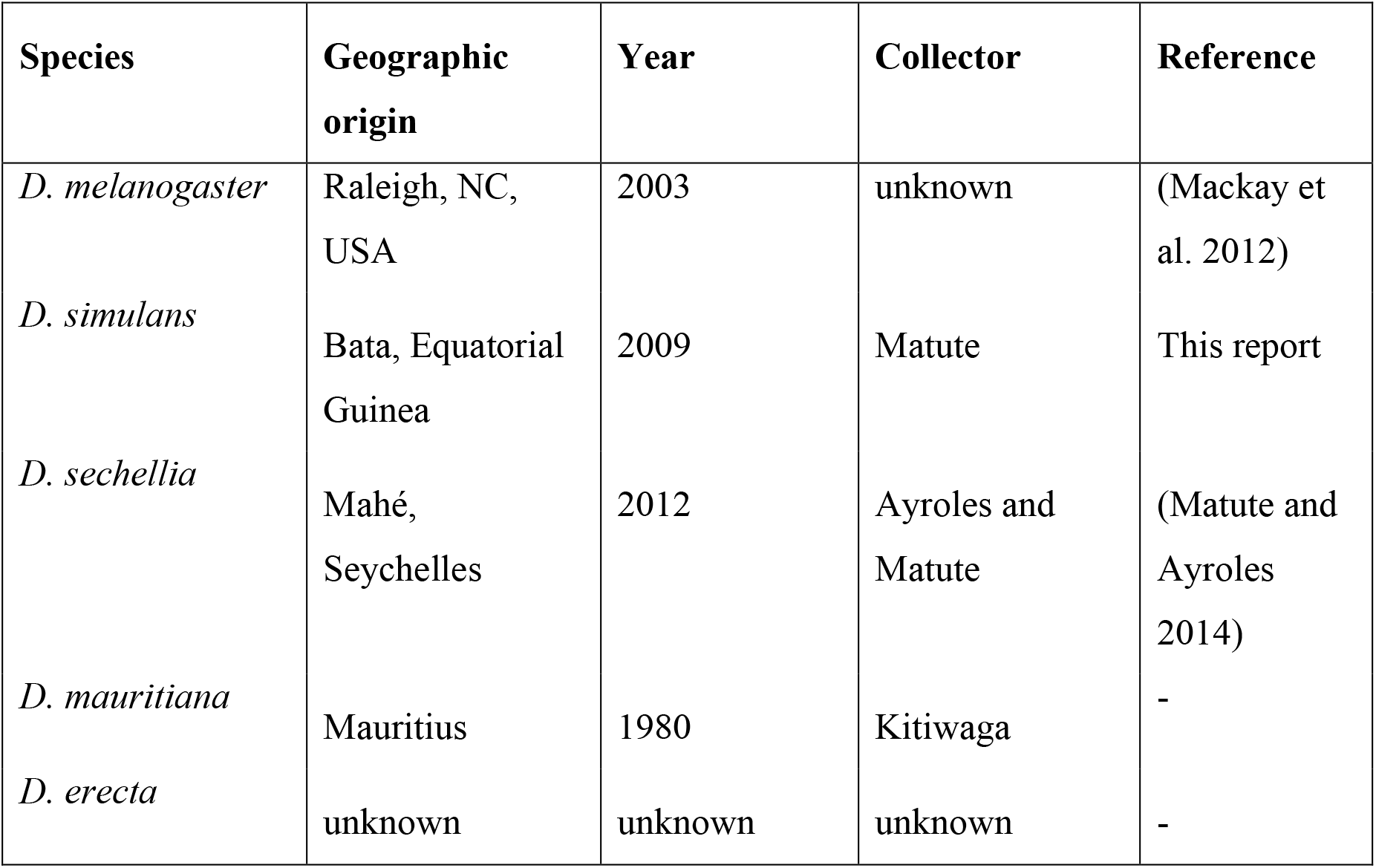
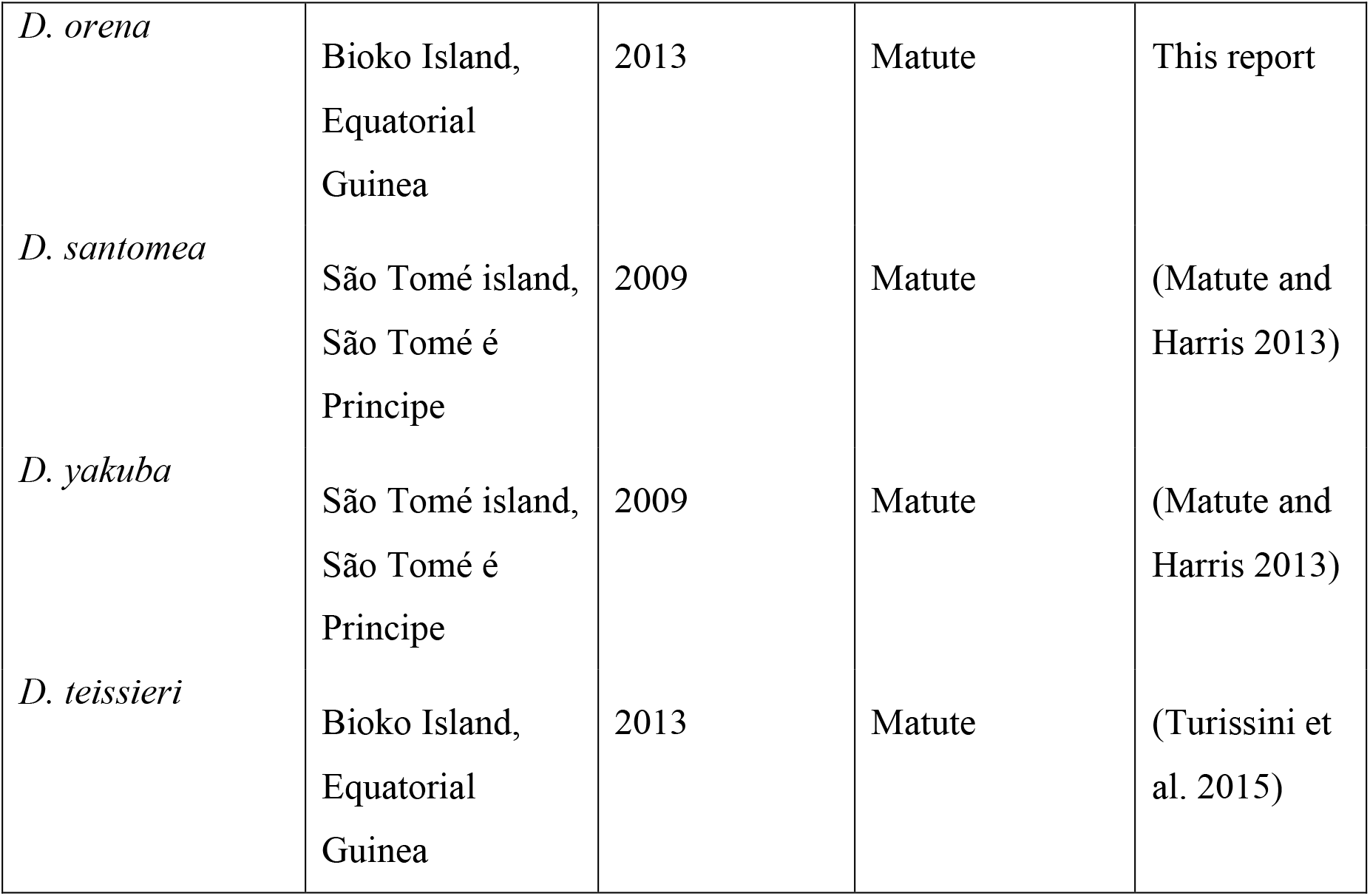
List of species, collection sites, and collector.

| Species | Geographic origin | Year | Collector | Reference |
| --- | --- | --- | --- | --- |
| *D. melanogaster* | Raleigh, NC, USA | 2003 | unknown | (Mackay et al. 2012) |
| *D. simulans* | Bata, Equatorial Guinea | 2009 | Matute | This report |
| *D. sechellia* | Mahé, Seychelles | 2012 | Ayroles and Matute | (Matute and Ayroles 2014) |
| *D. mauritiana* | Mauritius | 1980 | Kitiwaga | - |
| *D. erecta* | unknown | unknown | unknown | - |
| *D. orena* | Bioko Island,<br>Equatorial<br>Guinea | 2013 | Matute | This report |
| *D. santomea* | São Tomé island,<br>São Tomé é<br>Príncipe | 2009 | Matute | (Matute and<br>Harris 2013) |
| *D. yakuba* | São Tomé island,<br>São Tomé é<br>Príncipe | 2009 | Matute | (Matute and<br>Harris 2013) |
| *D. teissieri* | Bioko Island,<br>Equatorial<br>Guinea | 2013 | Matute | (Turissini et<br>al. 2015) |

**TABLE S14.** The number of scored individuals from each cross.

| **Cross** | **Total eggs** | **Dead Embryos** | **Egg cases** | **Pupae** | **Adults** |
| --- | --- | --- | --- | --- | --- |
| *mel* × *sim* | 96 | 0 | 93 | 48 | 45 |
| *mel* × *sim* | 109 | 1 | 99 | 45 | 38 |
| *mel* × *sim* | 102 | 2 | 94 | 50 | 44 |
| *mel* × *sim* | 112 | 3 | 92 | 52 | 51 |
| *mel* × *sim* | 69 | 0 | 64 | 29 | 23 |
| *mel* × *sech* | 69 | 2 | 54 | 22 | 6 |
| *mel* × *sech* | 59 | 4 | 51 | 32 | 11 |
| *mel* × *sech* | 78 | 8 | 65 | 30 | 12 |
| *mel* × *sech* | 69 | 10 | 55 | 29 | 16 |
| *mel* × *sech* | 79 | 2 | 43 | 22 | 14 |
| *mel* × *mau* | 111 | 6 | 101 | 57 | 56 |
| *mel* × *mau* | 99 | 1 | 93 | 47 | 43 |
| *mel* × *mau* | 140 | 1 | 122 | 70 | 66 |
| *mel* × *mau* | 45 | 3 | 40 | 40 | 19 |
| *mel* × *mau* | 88 | 4 | 76 | 45 | 40 |
| *mel* × *tei* | 14 | 8 | 6 | 6 | 6 |
| *mel* × *tei* | 22 | 9 | 13 | 10 | 9 |
| *mel* × *tei* | 41 | 25 | 16 | 8 | 5 |
| *mel* × *tei* | 34 | 23 | 11 | 12 | 15 |
| *mel* × *tei* | 20 | 8 | 12 | 12 | 12 |
| *mel* × *san* | 56 | 27 | 29 | 29 | 29 |
| *mel* × *san* | 97 | 45 | 50 | 46 | 44 |
| *mel* × *san* | 98 | 40 | 56 | 45 | 42 |
| *mel* × *san* | 60 | 32 | 25 | 23 | 23 |
| *mel* × *san* | 56 | 23 | 33 | 29 | 26 |
| *sim* × *mel* | 70 | 34 | 34 | 33 | 33 |
| *sim* × *mel* | 67 | 40 | 27 | 27 | 27 |
| *sim* × *mel* | 99 | 52 | 46 | 45 | 43 |
| *sim* × *mel* | 56 | 31 | 25 | 22 | 21 |
| *sim* × *mel* | 31 | 15 | 16 | 16 | 16 |
| *sim* × *sech* | 180 | 5 | 163 | 155 | 152 |
| *sim* × *sech* | 99 | 4 | 89 | 76 | 76 |
| *sim* × *sech* | 60 | 0 | 58 | 56 | 56 |
| *sim* × *sech* | 94 | 6 | 86 | 83 | 81 |
| *sim* × *sech* | 96 | 2 | 91 | 90 | 88 |
| *sim* × *mau* | 61 | 0 | 58 | 56 | 56 |
| *sim* × *mau* | 84 | 1 | 82 | 79 | 77 |
| *sim* × *mau* | 111 | 0 | 104 | 101 | 98 |
| *sim* × *mau* | 108 | 4 | 102 | 98 | 95 |
| *sim* × *mau* | 50 | 5 | 45 | 39 | 38 |
| *sim* × *tei* | 60 | 34 | 24 | 23 | 23 |
| *sim* × *tei* | 10 | 5 | 4 | 4 | 4 |
| *sim* × *tei* | 31 | 16 | 15 | 13 | 12 |
| *sim* × *tei* | 22 | 12 | 10 | 9 | 9 |
| *sim* × *tei* | 17 | 7 | 6 | 6 | 6 |
| *sim* × *san* | 43 | 24 | 14 | 13 | 13 |
| *sim* × *san* | 26 | 7 | 18 | 15 | 14 |
| *sim* × *san* | 19 | 9 | 8 | 8 | 8 |
| *sim* × *san* | 40 | 23 | 16 | 16 | 16 |
| *sim* × *san* | 31 | 14 | 16 | 16 | 15 |
| *sech* × *mel* | 67 | 28 | 31 | 23 | 23 |
| *sech* × *mel* | 50 | 17 | 32 | 28 | 28 |
| *sech* × *mel* | 48 | 22 | 25 | 19 | 19 |
| *sech* × *mel* | 56 | 34 | 20 | 16 | 16 |
| *sech* × *mel* | 33 | 18 | 14 | 12 | 12 |
| *sech* × *sim* | 21 | 3 | 17 | 16 | 16 |
| *sech* × *sim* | 22 | 1 | 21 | 18 | 15 |
| *sech* × *sim* | 13 | 0 | 13 | 12 | 11 |
| *sech* × *sim* | 15 | 1 | 14 | 14 | 14 |
| *sech* × *sim* | 19 | 1 | 17 | 16 | 16 |
| *sech* × *mau* | 24 | 1 | 23 | 23 | 23 |
| *sech* × *mau* | 22 | 3 | 17 | 16 | 15 |
| *sech* × *mau* | 45 | 0 | 43 | 37 | 34 |
| *sech* × *mau* | 26 | 1 | 24 | 20 | 19 |
| *sech* × *mau* | 25 | 2 | 19 | 19 | 18 |
| *sech* × *tei* | 19 | 9 | 10 | 8 | 5 |
| *sech* × *tei* | 12 | 6 | 6 | 5 | 4 |
| *sech* × *tei* | 19 | 10 | 9 | 5 | 4 |
| *sech* × *tei* | 14 | 6 | 7 | 4 | 3 |
| *sech* × *tei* | 22 | 14 | 8 | 8 | 9 |
| *sech* × *san* | 16 | 9 | 7 | 4 | 2 |
| *sech* × *san* | 5 | 1 | 4 | 0 | 0 |
| *sech* × *san* | 13 | 5 | 7 | 2 | 2 |
| *sech* × *san* | 23 | 12 | 11 | 2 | 2 |
| *sech* × *san* | 19 | 9 | 9 | 9 | 9 |
| *san* × *sech* | 31 | 15 | 14 | 12 | 0 |
| *san* × *sech* | 21 | 10 | 8 | 7 | 0 |
| *san* × *sech* | 17 | 11 | 6 | 5 | 0 |
| *san* × *sech* | 23 | 8 | 6 | 4 | 0 |
| *san* × *sech* | 30 | 14 | 10 | 8 | 0 |
| *mau* × *mel* | 23 | 11 | 11 | 10 | 10 |
| *mau* × *mel* | 99 | 38 | 51 | 46 | 41 |
| *mau × mel* | 29 | 13 | 14 | 14 | 14 |
| *mau × mel* | 96 | 38 | 51 | 40 | 22 |
| *mau × mel* | 74 | 28 | 40 | 37 | 31 |
| *mau × sim* | 71 | 4 | 67 | 67 | 67 |
| *mau × sim* | 114 | 5 | 104 | 102 | 100 |
| *mau × sim* | 67 | 2 | 61 | 59 | 56 |
| *mau × sim* | 104 | 4 | 100 | 93 | 91 |
| *mau × sim* | 58 | 2 | 55 | 55 | 55 |
| *mau × sech* | 52 | 3 | 48 | 47 | 45 |
| *mau × sech* | 96 | 1 | 94 | 89 | 89 |
| *mau × sech* | 80 | 1 | 74 | 72 | 67 |
| *mau × sech* | 55 | 0 | 53 | 49 | 43 |
| *mau × sech* | 37 | 1 | 34 | 34 | 34 |
| *mau × tei* | 13 | 6 | 7 | 5 | 5 |
| *mau × tei* | 16 | 8 | 8 | 6 | 6 |
| *mau × tei* | 49 | 25 | 22 | 20 | 19 |
| *mau × tei* | 14 | 5 | 8 | 5 | 4 |
| *mau × tei* | 40 | 23 | 16 | 15 | 15 |
| *mau × san* | 23 | 11 | 12 | 8 | 4 |
| *mau × san* | 19 | 9 | 10 | 7 | 6 |
| *mau × san* | 11 | 5 | 5 | 4 | 4 |
| *mau × san* | 19 | 10 | 7 | 7 | 7 |
| *mau × san* | 7 | 2 | 4 | 0 | 0 |
| *mau × yak* | 14 | 9 | 3 | 0 | 0 |
| *mau × yak* | 30 | 14 | 15 | 6 | 5 |
| *mau × yak* | 22 | 10 | 11 | 7 | 4 |
| *mau × yak* | 11 | 5 | 5 | 4 | 1 |
| *mau × yak* | 21 | 14 | 6 | 5 | 2 |
| *mau × ore* | 9 | 4 | 5 | 5 | 5 |
| *mau × ore* | 45 | 23 | 19 | 6 | 0 |
| *mau × ore* | 11 | 6 | 5 | 2 | 2 |
| *mau × ore* | 12 | 6 | 6 | 4 | 4 |
| *mau × ore* | 19 | 10 | 8 | 8 | 7 |
| *mau × ere* | 51 | 24 | 27 | 25 | 22 |
| *mau × ere* | 40 | 20 | 20 | 16 | 15 |
| *mau × ere* | 39 | 20 | 16 | 16 | 11 |
| *mau × ere* | 20 | 11 | 8 | 7 | 7 |
| *mau × ere* | 21 | 10 | 11 | 9 | 9 |
| *yak × mau* | 11 | 6 | 5 | 2 | 1 |
| *yak × mau* | 15 | 7 | 5 | 4 | 4 |
| *yak × mau* | 14 | 6 | 6 | 3 | 2 |
| *yak × mau* | 19 | 10 | 9 | 6 | 4 |
| *yak × mau* | 15 | 8 | 7 | 4 | 3 |
| *yak* × *san* | 109 | 0 | 109 | 109 | 109 |
| *yak* × *san* | 103 | 2 | 99 | 95 | 95 |
| *yak* × *san* | 102 | 0 | 99 | 98 | 98 |
| *yak* × *san* | 33 | 5 | 27 | 25 | 24 |
| *yak* × *san* | 103 | 4 | 99 | 98 | 98 |
| *yak* × *tei* | 56 | 6 | 48 | 45 | 41 |
| *yak* × *tei* | 44 | 1 | 38 | 31 | 29 |
| *yak* × *tei* | 78 | 2 | 74 | 70 | 66 |
| *yak* × *tei* | 49 | 3 | 46 | 45 | 45 |
| *yak* × *tei* | 50 | 1 | 42 | 39 | 34 |
| *san* × *mau* | 17 | 5 | 5 | 5 | 5 |
| *san* × *mau* | 36 | 2 | 20 | 14 | 11 |
| *san* × *mau* | 15 | 9 | 2 | 0 | 0 |
| *san* × *mau* | 21 | 1 | 11 | 10 | 6 |
| *san* × *mau* | 16 | 8 | 5 | 5 | 4 |
| *san* × *yak* | 61 | 5 | 52 | 49 | 47 |
| *san* × *yak* | 66 | 4 | 62 | 61 | 60 |
| *san* × *yak* | 96 | 0 | 92 | 91 | 89 |
| *san* × *yak* | 78 | 0 | 67 | 61 | 56 |
| *san* × *yak* | 77 | 4 | 71 | 69 | 65 |
| *san* × *tei* | 94 | 4 | 90 | 89 | 88 |
| *san* × *tei* | 22 | 5 | 16 | 16 | 14 |
| *san* × *tei* | 41 | 1 | 39 | 37 | 33 |
| *san* × *tei* | 52 | 2 | 48 | 45 | 41 |
| *san* × *tei* | 30 | 3 | 26 | 26 | 26 |
| *tei* × *mau* | 45 | 29 | 15 | 14 | 14 |
| *tei* × *mau* | 26 | 10 | 15 | 14 | 12 |
| *tei* × *mau* | 14 | 2 | 11 | 4 | 0 |
| *tei* × *mau* | 29 | 9 | 16 | 12 | 11 |
| *tei* × *mau* | 26 | 8 | 15 | 9 | 8 |
| *tei* × *yak* | 80 | 1 | 77 | 75 | 75 |
| *tei* × *yak* | 109 | 4 | 100 | 98 | 98 |
| *tei* × *yak* | 72 | 1 | 67 | 67 | 67 |
| *tei* × *yak* | 53 | 2 | 49 | 46 | 45 |
| *tei* × *yak* | 102 | 3 | 96 | 88 | 87 |
| *tei* × *san* | 68 | 2 | 64 | 64 | 64 |
| *tei* × *san* | 95 | 4 | 89 | 82 | 78 |
| *tei* × *san* | 84 | 3 | 77 | 72 | 69 |
| *tei* × *san* | 109 | 2 | 104 | 99 | 98 |
| *tei* × *san* | 103 | 1 | 99 | 87 | 74 |

**TABLE S15.**
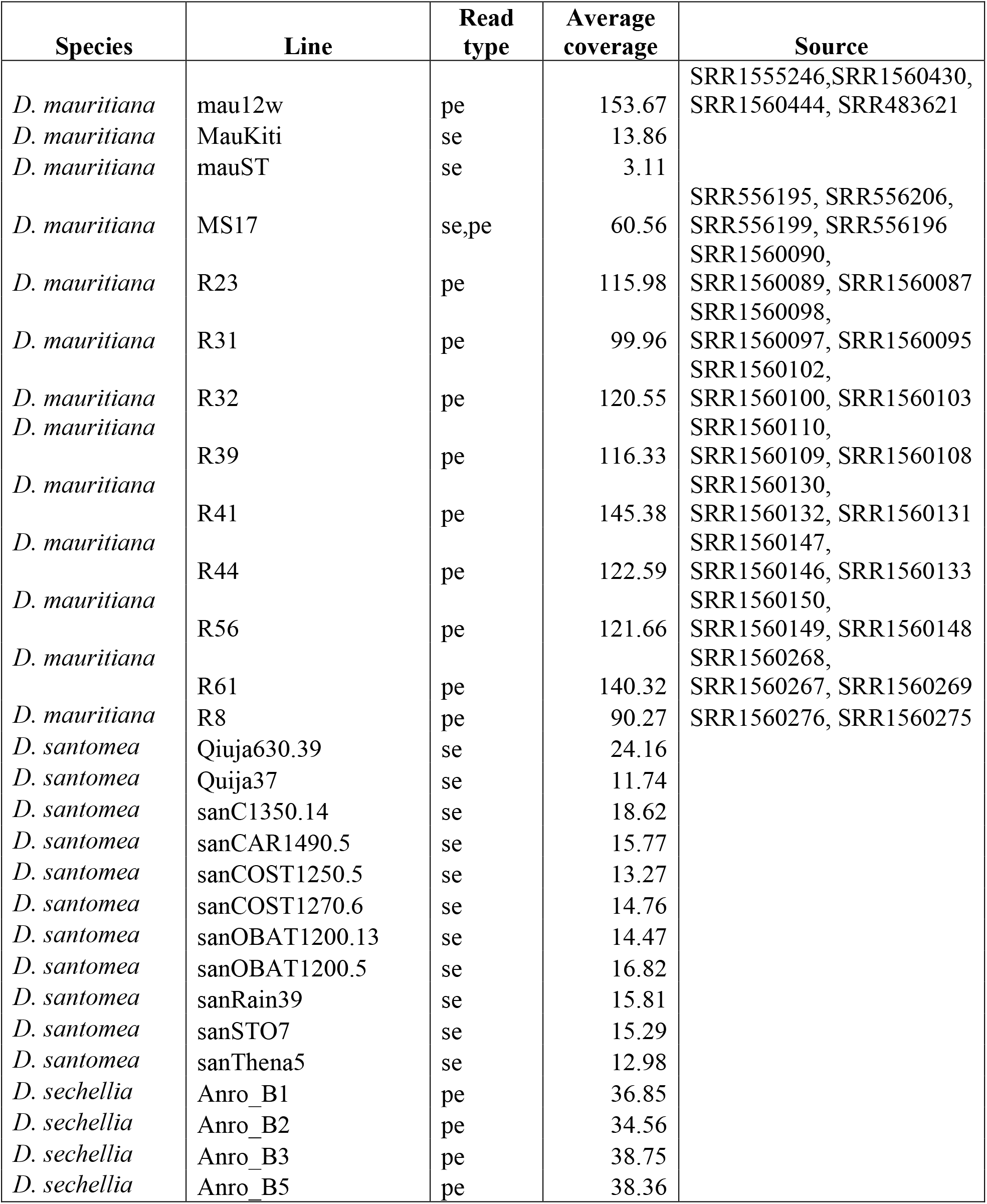
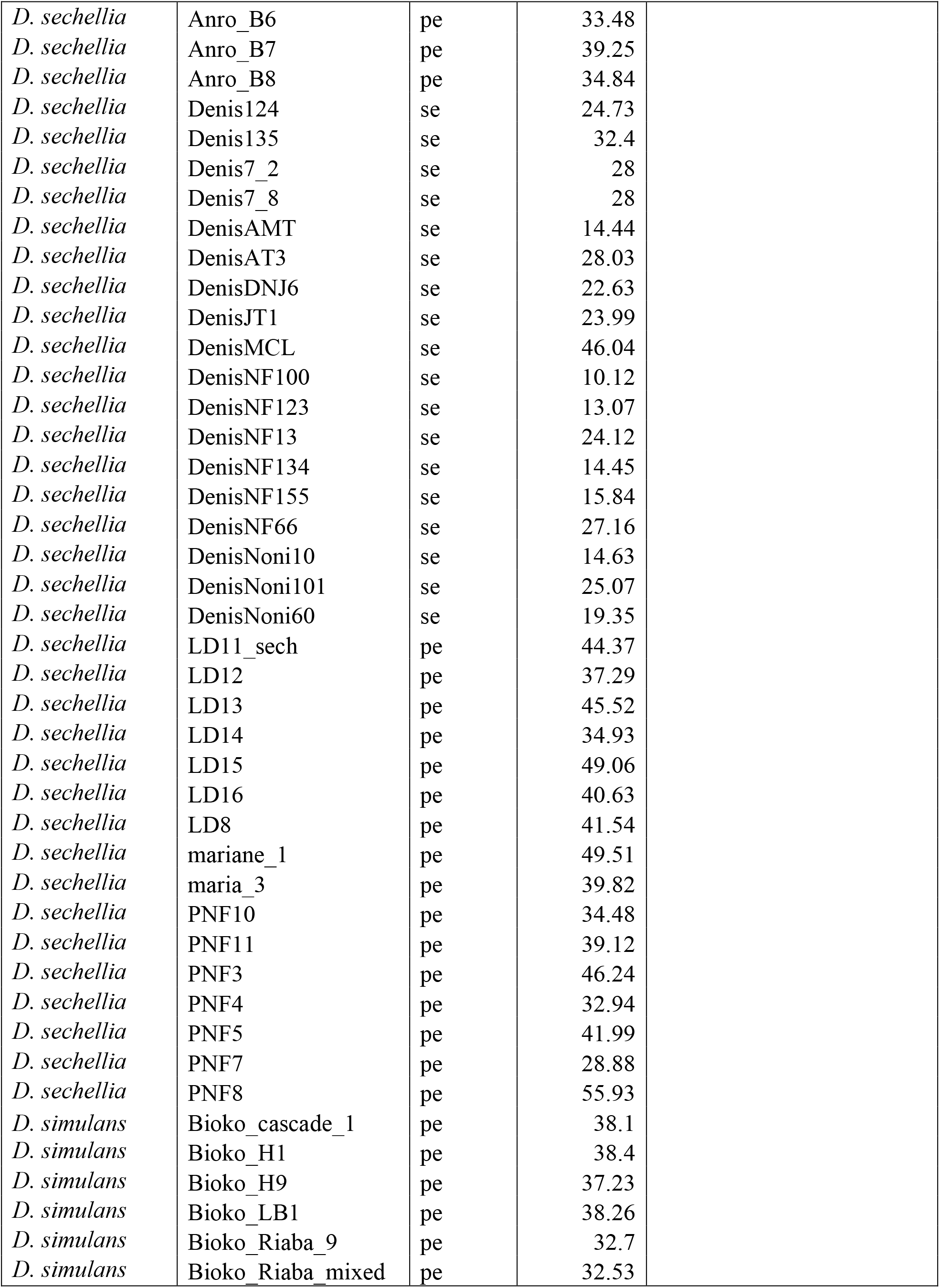
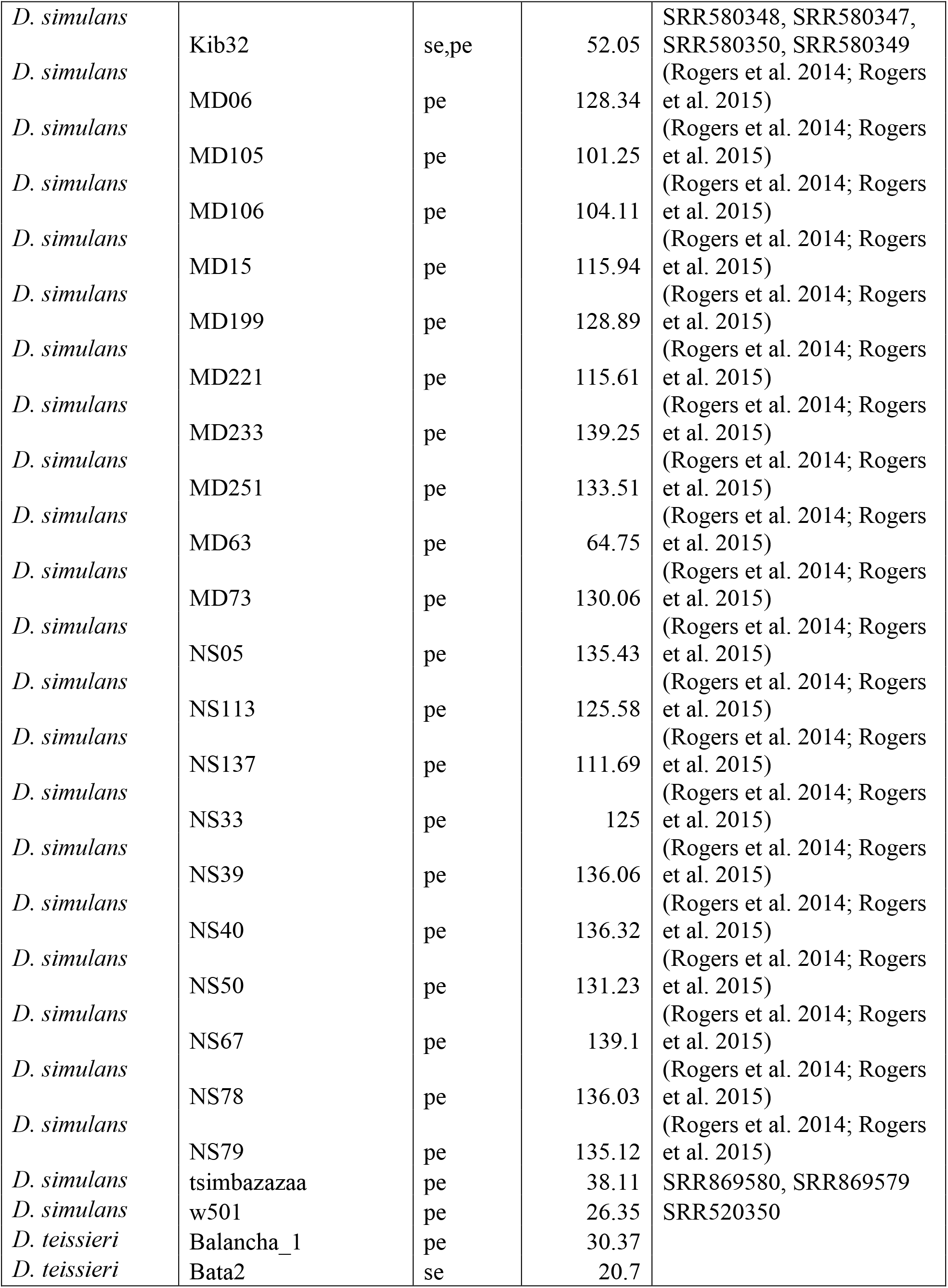
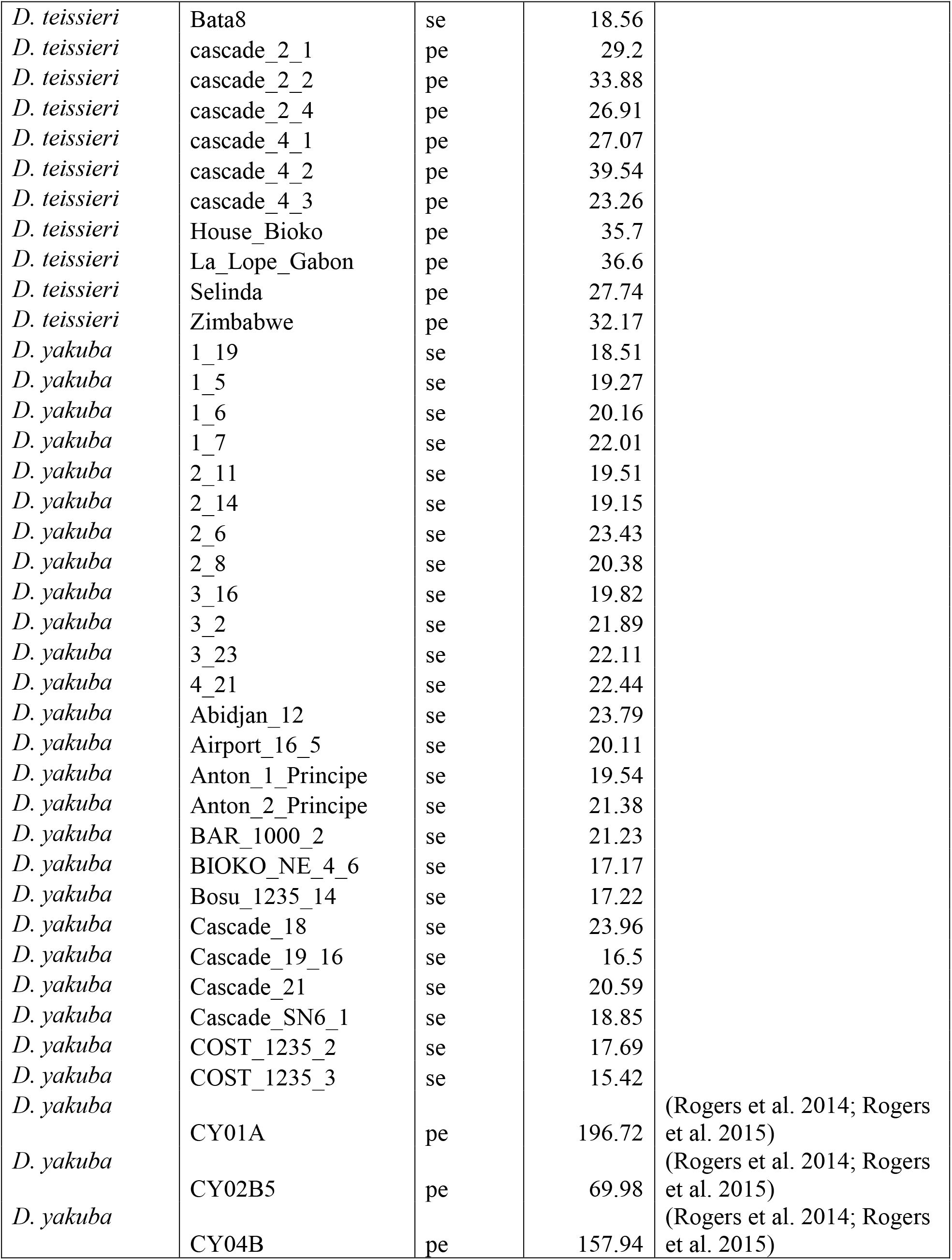
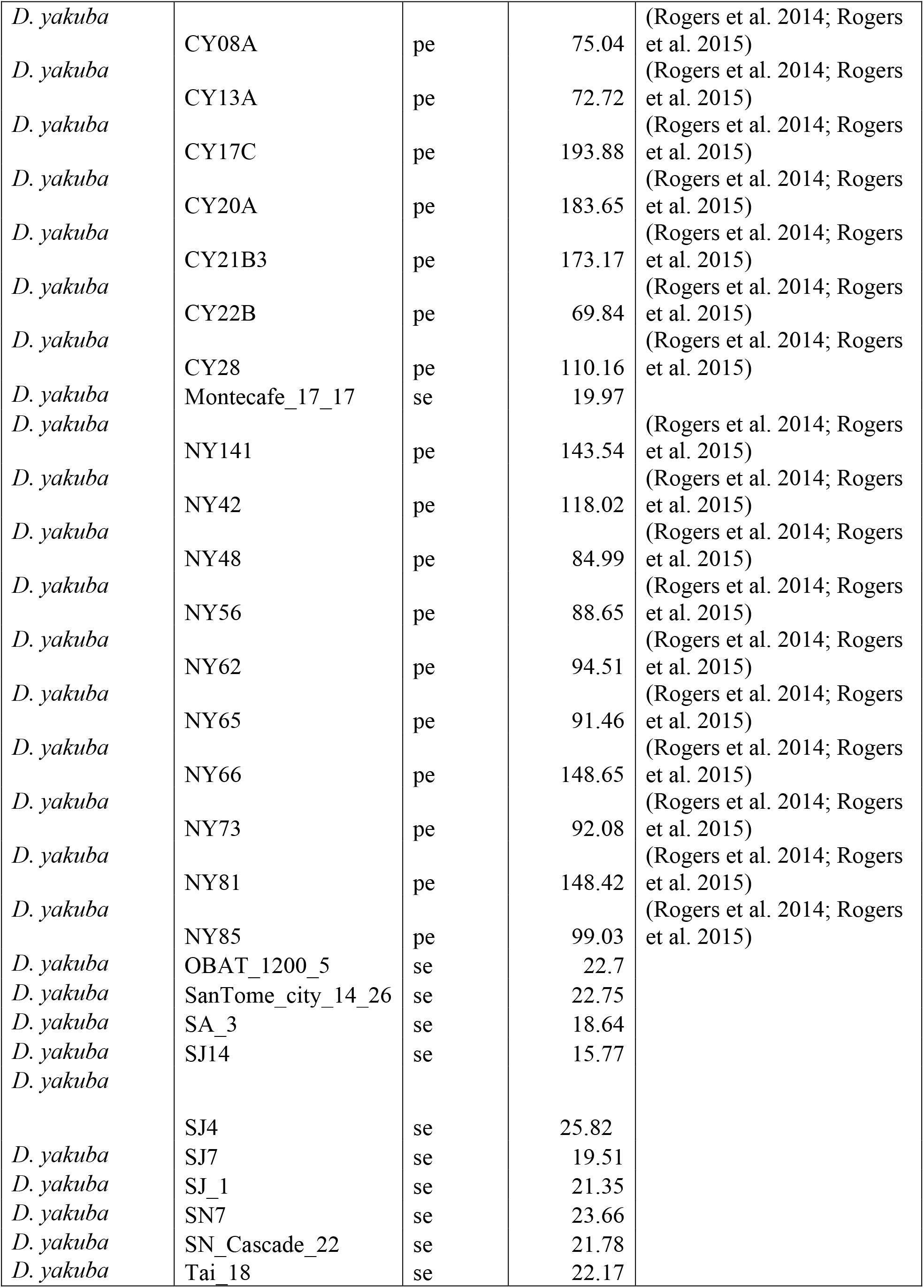
Sequencing type (se: Single end; pe: Paired end), and coverage for each isofemale line.

**TABLE S16.**
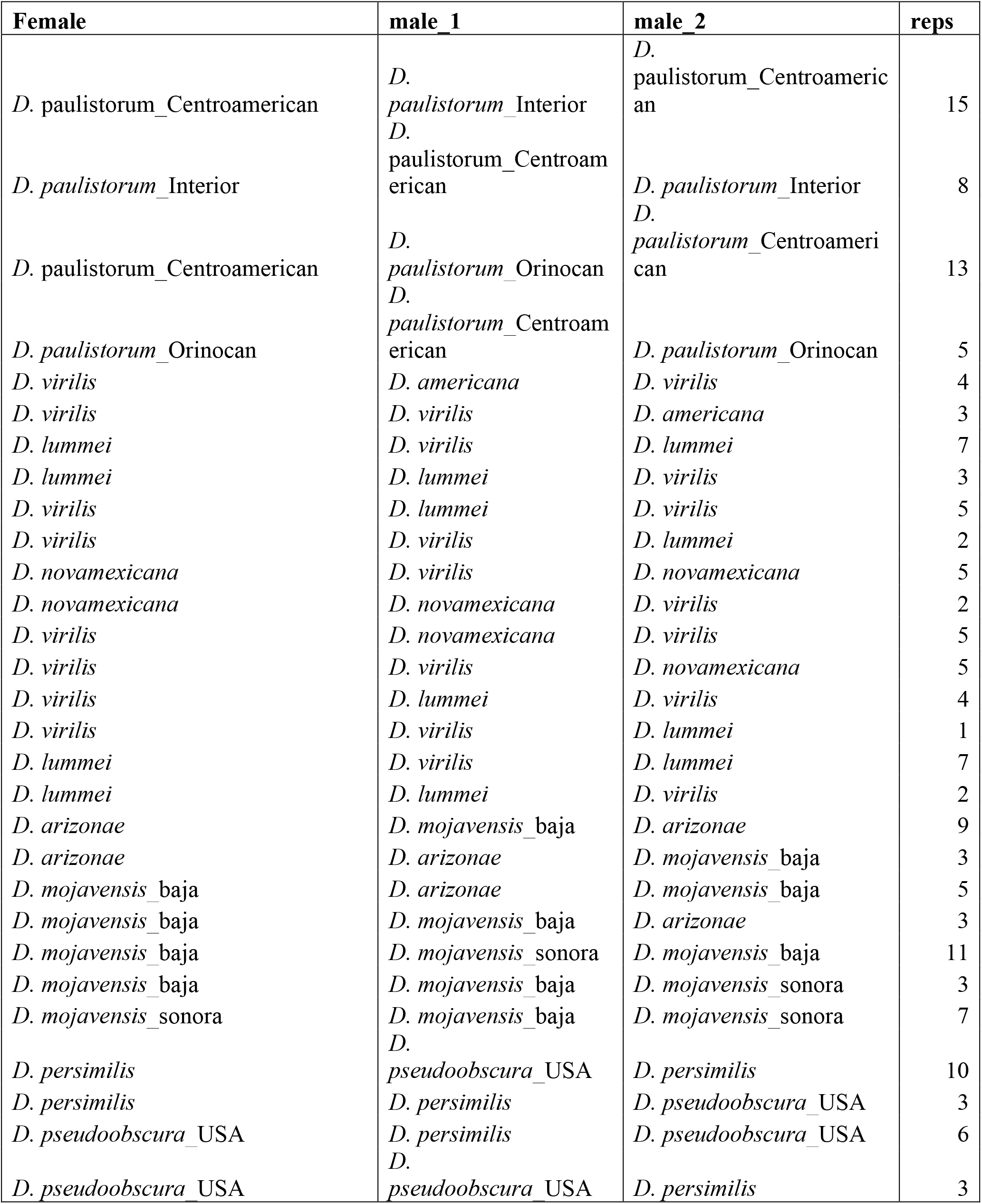
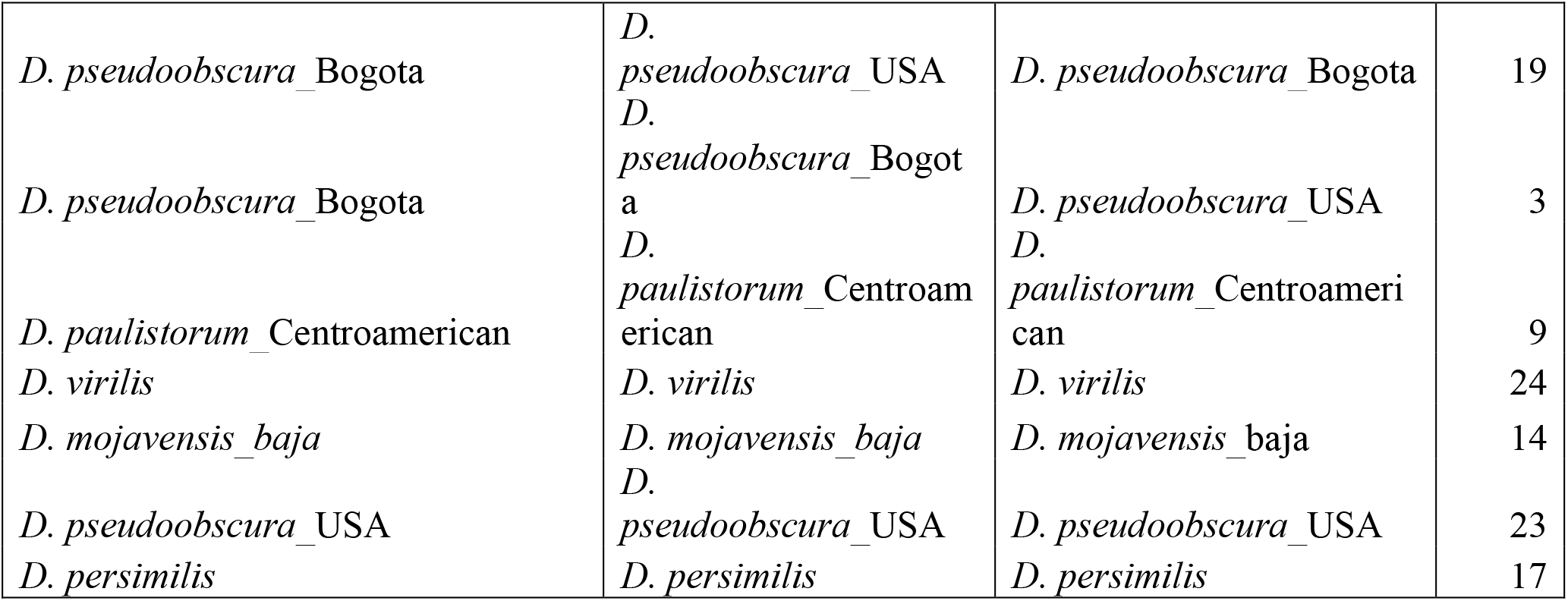
Sample sizes of conspecific sperm precedence experiments in species subgroups different from *melanogaster* (shown in Table S2).

